# Discovery and Optimization of Narrow Spectrum Inhibitors of Tousled Like Kinase 2 (TLK2) Using Quantitative Structure Activity Relationships

**DOI:** 10.1101/2023.12.28.573261

**Authors:** Christopher R. M. Asquith, Michael P. East, Tuomo Laitinen, Carla Alamillo-Ferrer, Erkka Hartikainen, Carrow I. Wells, Alison D. Axtman, David H. Drewry, Graham J. Tizzard, Antti Poso, Timothy M. Willson, Gary L. Johnson

**Author notes:** Corresponding authors.; (C. R. M. Asquith) (G. L. Johnson) E-mail address (C. R. M. Asquith) and (G. L. Johnson).

## Abstract

The oxindole scaffold has been the center of several kinase drug discovery programs, some of which have led to approved medicines. A series of two oxindole matched pairs from the literature were identified where TLK2 was a potent off-target kinase. The oxindole as long been considered a promiscuous inhibitor template, but across these 4 specific literature oxindoles TLK2 activity was consistent, while the kinome profile was radically different from narrow to broad spectrum coverage. We synthesized a large series of analogues and through quantitative structure-activity relationship (QSAR) analysis, water mapping of the kinase ATP binding sites, small-molecule x-ray structural analysis and kinome profiling, narrow spectrum, sub-family selective, chemical tool compounds were identified to enable elucidation of TLK2 biology.

## 1. Introduction

Tousled-like kinase 2 (TLK2) is a ubiquitously expressed, nuclear serine/threonine kinase. Kinase activity of TLK2 and its closest human paralog TLK1 is activated by autophosphorylation, hetero- and homo-dimerization, and oligomerization through conserved coiled-coiled domains.^1^ TLK kinase activity is highest during S-phase where TLKs phosphorylate ASF1 histone chaperones.^2^ DNA rapidly associates with histones during replication and ASF1 chaperones regulate the soluble pool of histones H3 and H4 during this process.^3^ Phosphorylation of ASF1 chaperones by TLKs stabilizes ASF1 protein,^4^ and promotes interactions between ASF1 and histones as well as histones with downstream chaperones CAF1 and HIRA1. TLK2, but not TLK1, was also shown to regulate G2/M checkpoint recovery after DNA damage through an ASF1A dependent mechanism.^5^ TLKs have ASF1 independent functions during DNA replication through the regulation of replication forks.^6^ Depletion of TLKs results in replication fork stalling, the accumulation of single-stranded DNA, and ultimately, DNA damage and cell cycle arrest in G1. These phenotypes were rescued by ectopic expression of WT TLKs but not kinase dead TLKs. Analysis of chromatin accessibility using the Assay for Tranposase-Accessible Chromatin using sequencing (ATAC-seq) following knockdown of TLK1 or TLK2 increased accessibility and transcription of heterochromatin. As a result, loss of TLKs induced alternate lengthening of telomeres thereby activating the cGAS-STING-TBK1 mediated immune response. Consistent with this function, high levels of TLK2 in patients were associated with poor innate and adaptive immune responses in tumors and, potentially, immune evasion.^7^ Thus, TLKs, and more specifically their kinase activity, are essential for proper chromatin assembly, DNA replication, and maintenance of heterochromatin.

The cellular functions of TLK1 and TLK2 are predominantly overlapping but their clinical implications are more diverse which may suggest other distinct functions. In the present study, we focus on TLK2. TLK2 has been implicated in neurodevelopmental disease and intellectual deficiency.^8-10^ Neurodevelopmental defects manifested with TLK2 haploinsuffiency and were more severe in a single, homozygous case. TLK2 mutations observed in these studies resulted in loss of TLK2 kinase activity.^11^ In cellular models of latent gammaherpesvirus infections, knockdown of TLK1 or TLK2 was sufficient to reactivate latent Epstein-Barr virus infection whereas only knockdown of TLK2 resulted in reactivation of latent Kaposi sarcoma-associated herpesvirus.^12^ Latency allows viral infections to evade the immune system and antiviral therapies resulting in lifelong, incurable infection. Reactivation of virus in combination with antiviral therapy is a promising approach known as “shock and kill” that could potentially cure these latent viral infections.^13^ TLK2 has also been implicated in breast cancer and glioblastoma where TLK2 is amplified or over-expressed in a high percentage of patients and where higher expression levels are associated with poor patient outcomes.^14-16^ In cell lines from both cancer types, ectopic expression of TLK2 lead to enhanced aggressiveness and activated SRC signalling whereas knockdown or pharmacological inhibition of TLK2 slowed growth and inhibited invasion. Knockout of TLK2 in mice resulted in late embryonic lethality due to defective trophoblast differentiation and placental failure.^17^ Conditional knockout of TLK2 to bypass the placental defect led to healthy mice suggesting that TLK2 was otherwise dispensable for development and viability. These findings suggest that inhibition of TLK2 may be tolerable in normal, healthy cells in patients. Thus, TLK2 is a very promising target for drug development in multiple cancer types and as a latency reversing agent in shock and kill approaches for latent viral infections.^13-14^

While some recent advances have been made around TLK2 chemical tools, with a recent advancement based on indirubin based derivatives, these analogues still inhibit both TLK1 and TLK2 with limited selectivity information.^18^ There are currently no potent and selective TLK2 inhibitors as chemical probes to investigate TLK2 biology.^19^ This despite several concerted efforts to map and identify leads within existing ATP-competitive kinase inhibitors and to define inhibitor chemical space.^19-28^

## 2. Results

Currently the only small molecule inhibitors available to modulate TLK2 are compounds with very broad kinome coverage including staurosporine, Syk inhibitor R406 and RTK inhibitor sunitinib, all of which inhibit TLK2 as an undesired off-target with low relative potency (**Figure 1**). However, the sunitinib oxindole based scaffold has demonstrated tractability on TLK2 where small modifications to afford SU14813 still maintain potency on TLK2 even when coverage has been reduced from 187 to 149 kinases. This is further supported by additional results of an enzyme assay-based panel identifying two further oxindoles, GW506033X and SU9516 that had 84% and 57% inhibition of TLK2 at 0.5 μM, respectively.^23^ The narrower kinome profile of SU9516 provided an attractive starting point for progress towards the development of a selective chemical probe for TLK2. Herein we report the synthesis, modelling, and characterization of oxindole based inhibitors as potent narrow-spectrum TLK2 inhibitors.

**Figure 1.**
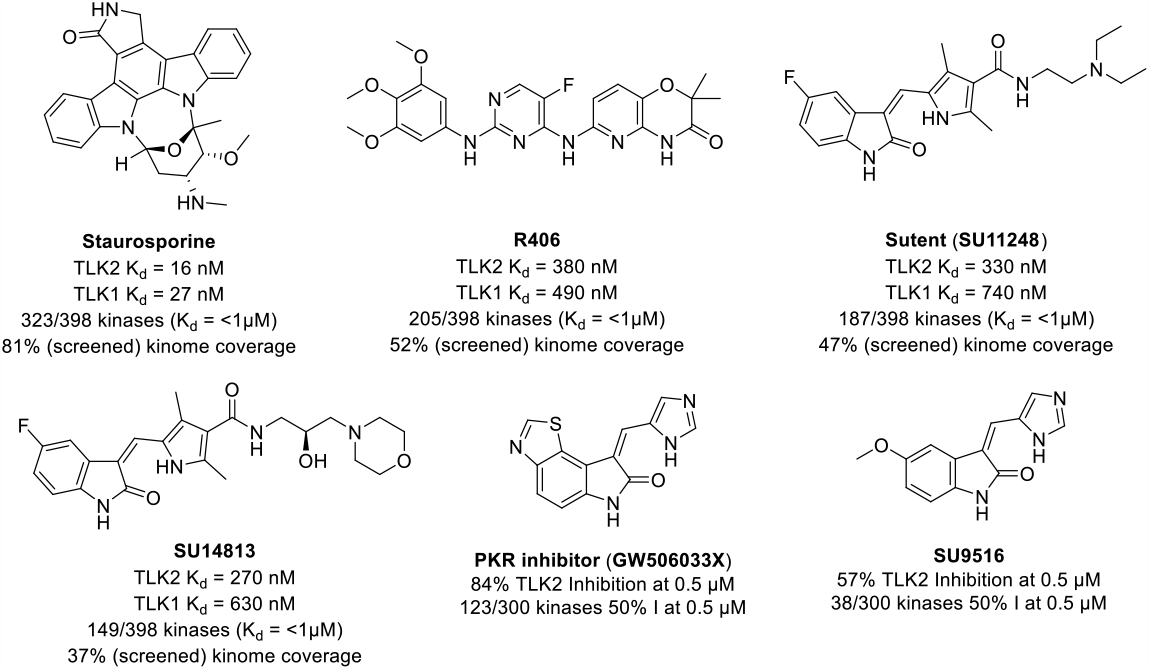
Previously reported kinase inhibitors with off-target of TLK2 inhibition, including two matched pair oxindoles.

Having identified the oxindole scaffold as a promising inhibitor chemotype, we first screened 21 commercially available oxindole based kinase inhibitors (**Figure 2**)^29-35^ in a TLK2 enzyme assay (**Table S1-S2**).^23^ This enabled us to quickly map the TLK2 vs the close kinome landscape and to prioritize our efforts around the smaller SU9516 and GW506033X oxindoles.

**Figure 2.**
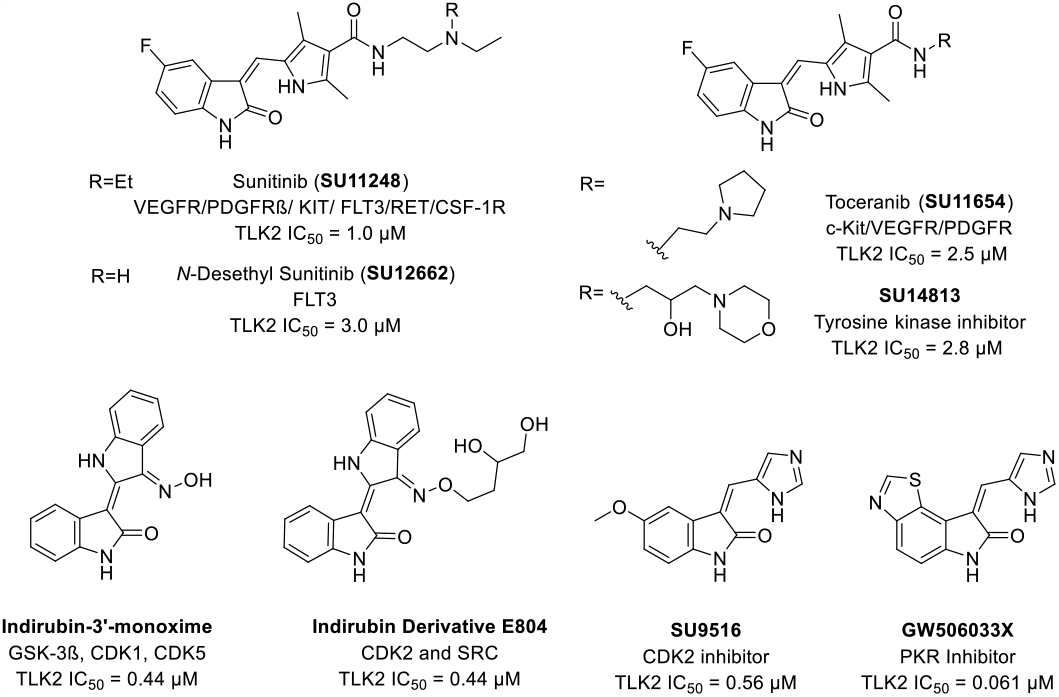
Screening of literature inhibitors with the oxindole scaffold with activities below 10 μM (see figure S1 for additional screening and structures). Format: Name, primary target, TLK2 IC_50_

Several interchangeable matched pairs including sunitinib (SU11248), toceranib (SU11654), SU12662 and SU14813 all demonstrated consistent low single digit micromolar activity against TLK2. While all these compounds have promiscuous kinome profiles,^20-22^ this data increased our confidence that the oxindole may provide a good starting point for TLK2 probe development. The two indirubin derivatives had the same activity against TLK2 (IC_50_ = 0.44 μM) indicating that oxindoles were bound in the traditional binding mode, with space to grow the solvent exposed region of the molecule.^29-35^ The only other two oxindoles that showed activity below 1 micromolar were SU9516 and GW506033X, both containing an imidazole head group. The narrow kinome spectrum of SU9516 (10 kinases inhibited to 80% at 1 μM) combined with the potency of GW506033X (IC_50_ = 0.061 μM) afforded a promising indication that a potent narrow spectrum compound was possible.^23^ We also noted that the same major kinase off-targets namely, fms-like tyrosine kinase 3 (FLT3) and tropomyosin receptor kinase C (TRKC) were recurring themes and likely a prognostic marker for further promiscuity across the kinome. In addition to screening for TLK2 potency, we screened these two off-targets as a micro-panel to check for selectivity between molecules before kinome-wide screening.^36-38^

Having decided to focus our attention on SU9516 (**1**) and related GW506033X, we first sought to explore the structural features required for TLK2 inhibition (**Scheme 1**). Initially we synthesized a series of direct SU9516 analogs (**1**-**35**) using commercially available 5- and/or 6-substituted oxindoles and the Knoevenagel condensation.^39-41^ The oxindoles along with a corresponding aldehyde were suspended in ethanol and treated with piperidine under reflux to afford condensation products **1**-**35** in varying yields (15-85%). The products were isolated by direct crystallization from the crude reaction mixture or following chromatographic purification. A second set of analogues was synthesized in a two-step procedure: first using a Friedel Craft acylation by treatment of unsubstituted oxindole with aluminum (III) chloride and the corresponding acid chloride in dichloromethane to afford the 5-position ketone substituted products **36**-**39** in acceptable to good yields (30-74%).^42-44^ Second, the ketone products (**36**-**39**) were then treated with the corresponding aldehyde, piperidine under reflux to afford condensation products **40**-**48** in acceptable to good yields (30-74%). To afford a series of sulphonamide analogs, we first treated the unsubstituted oxindole with neat chlorosulphonic acid at -10 °C, affording **49** in good yield (68%).^45^ The sulfonyl chloride **49** was treated with the corresponding amine to afford the sulphonamide building blocks **50**-**51** in excellent yields, 78% and 85% respectively. The sulphonamides **50**-**51** were then treated with the previous Knoevenagel conditions with the respective aldehyde to afford the condensation products **52**-**55** in good yield (41-75%).

To access a novel 5-position *iso*propyl ether oxindole, a standard Mitsunobu reaction was performed on the 5-hydroxy oxindole with diethyl azadicarboxylate (DEAD), triphenylphosphine, *iso*propyl alcohol in THF at 0 °C to afford 5-*iso*propoxyindolin-2-one (**56**) in 30% yield.^46-48^ The ether **56** was then subject to the Knoevenagel conditions with the corresponding aldehyde to afford the desired condensation products **57**-**59** in good yields (53-62%). Finally, we prepared a series of substituted oxindole amides, furnished with a standard HATU coupling,^49^ in the presence of DIPEA and THF. The first set (**60**-**64**) was afforded in good yields (63-78 %), followed by a series of condensations under standard conditions to render a series of 5-postion substituted amide final products (**65**-**76**) in a range of yields (15-71%). The second set of amide substitutions (**77**-**88**) were also afforded in good to excellent yields (48-91%) followed by standard Knoevenagel conditions with the appropriate aldehyde to produce the condensation products (**89**-**107**) in a range of yields (11-71%).

Having prepared a series of oxindole analogues, these were screened in a TLK2 enzyme assay in a 10-point dose dependent format for TLK2 to determine an IC_50_ value. The maximum concentration used was 20 μM with an ATP concentration of 10 μM. The corresponding collateral target evaluation was on FLT3 and TRKC which were screened at a 0.5 μM, consistent with previous reports.^23^ The most advanced compounds were then subjected to a Kinase-Glo assay and additional TLK2 assay. The lead compounds were then screened in a DiscoverX kinome panel assay (>400 kinases) to assess their selectivity across the kinome.^20-22^

In the first series of analogues we followed up on the initial hit compound (*Z*)-3-((1*H*-imidazol-5-yl)methylene)-5-methoxyindolin-2-one (**1**) which when resynthesized produced a compound with consistent potency on TLK2 (IC_50_ = 240 nM) (**Table 1**). The substitution of the imidazole with the pyrrole (**2**) resulted in a 6-fold drop in potency on TLK2. Introduction of the sterically hindered dimethyl group (**3**) afforded no activity on TLK2 at the maximum concentration tested but still showed activity on FLT3 and TRKC. The introduction of a pyrazole group (**4**) lead to a 36-fold reduction of TLK2 activity with respect to **1**. The 2-substituted imidazole (**5**) afforded a compound that was near equipotent with **1** against TLK2. However, none of the simple 5 membered analogues (**2**-**5**) improved activity on TLK2. A switch to a 6-member 2-pyridyl (**6**) and 3-pyridyl (**7**) head group ring system demonstrated no activity against TLK2 at the maximum concentration tested but also no activity against TRKC and FLT3. Removal of the 5-methoxy group to afford the unsubstituted oxindole (**8**) lead to a 2-fold drop in potency against TLK2. Interestingly the pyrrole substitution **9** shows the same potency as **8** unlike the corresponding drop observed between **1** and **2**.

**Table 1.**
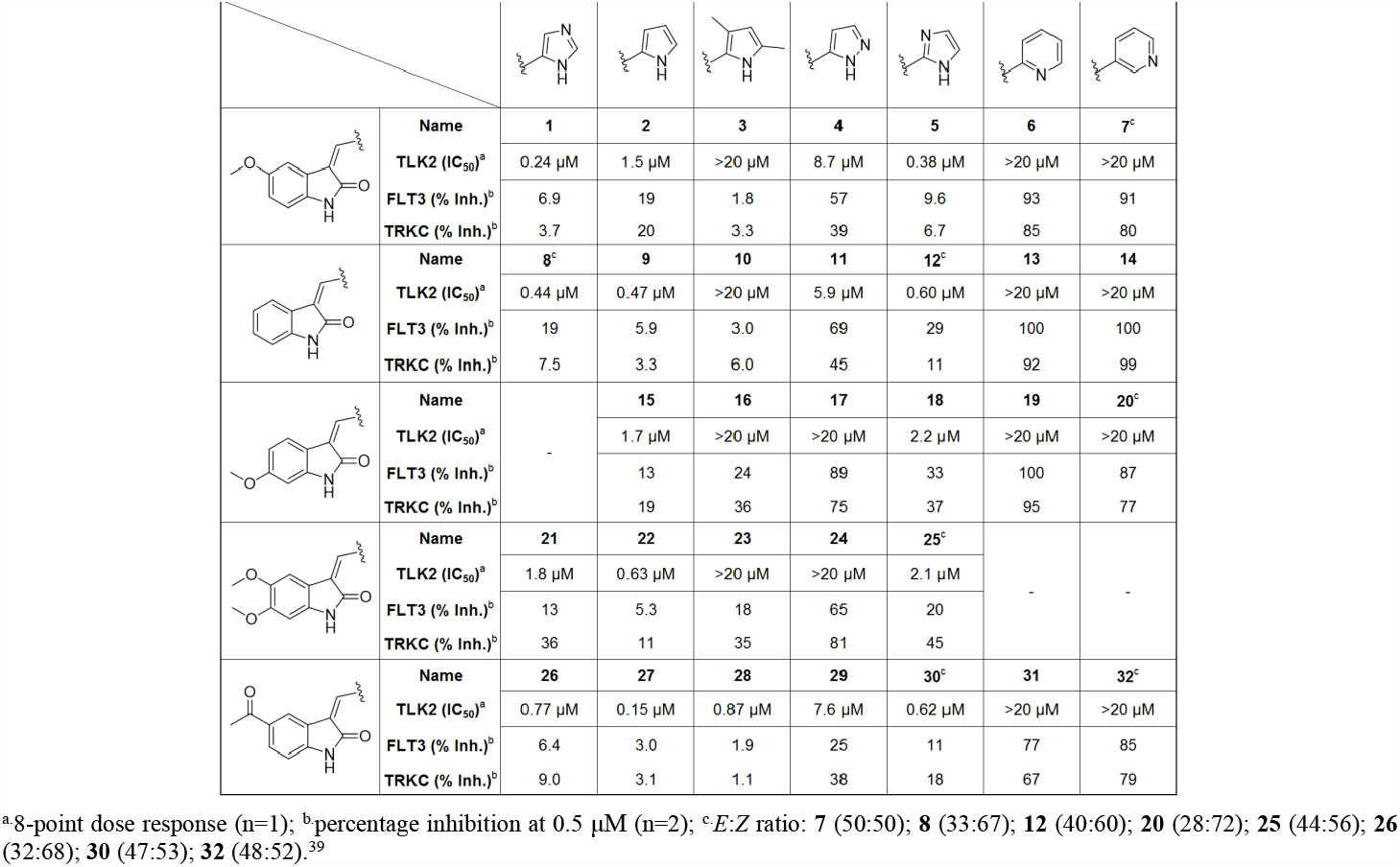
Matrix array of SU9516 derivatives with 5,6-oxindole substitutions and different heterocyclic head groups.

The corresponding 3,5-dimethyl pyrrole substitution **10** was inactive against TLK2, while the pyrazole **11** showed a 13-fold drop compared with **8**. The 2-substituted imidazole analogue **12** was equipotent with **8**, while the two pyridyl analogues **13** and **14** demonstrated no activity against TLK2 or TRKC/FLT3. Switching to the 6-methoxy oxindole substitution **15**-**20** lead to a net reduction of around 10-fold in activity on nearly all analogues with only the pyrrole maintaining TLK2 inhibition at the same level as **2**.

**Scheme 1.**
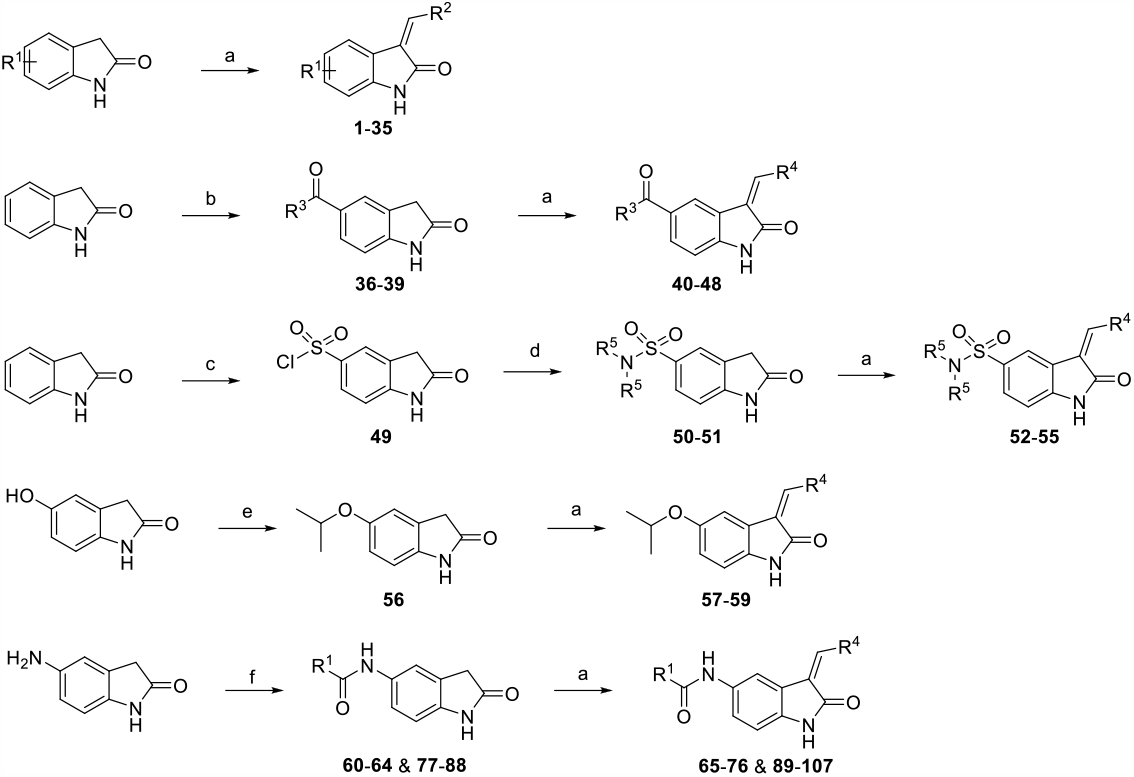
Synthetic route to prepare the oxindole analogues - *reagents and conditions* a) R_2_CHO, piperidine, EtOH, reflux, 4-24 h; b) R^1^COCl, AlCl_3_, CH2Cl_2_, rt, 48 h; c) ClSO_3_H, -5 °C, 2 h; d) (R^1^)_2_NH, EtN(CH(CH_3_)_2_)_2_ THF, rt, 12 h; e) R^1^OH, diethyl azadicarboxylate (DEAD), PPh_3_, rt, -5 °C to rt; f) R^1^COCl, HATU, EtN(CH(CH_3_)_2_)_2_, THF, rt, 12 h.

Given the decrease in inhibition of TLK2 activity at the 6-position, we decided to explore the 5,6-dimethxoyoxindole core **21**-**25** to assess potential synergistic effects of a double substitution pattern. The results of the 5 analogues were consist with the mono-substituted 6-methoxy except for the pyrrole **22** that showed a 3-fold jump in potency against TLK2. However, **22** also showed a boost in TRKC and FLT3 inhibition, meaning it was also likely an overall increase in kinome inhibition rather than something solely related to TLK2. We decided to focus our attention on the 5-postion as this seemed to be preferable for TLK2 inhibition. Switching the core to the 5-acetyloxindole **26**-**32** afforded our first compound **27** with more potency against TLK2 IC_50_ = 150 nM, than the original starting point (**1**) (TLK2 IC_50_ = 240 nM). The imidazole analogue was 3-fold less active against TLK2, but the pyrrole was 60% more potent against TLK2. The 3,5-dimethylpyrolle analogue (**28**) also demonstrated inhibition of TLK2 for the first time with an IC_50_ = 870 nM. The pyrazole **29** was significantly weaker nearly 10-fold less potent than the imidazole parent **26**. The 2-substituted imidazole (**30**) is as active as the parent while the pyridyl analogues **31** and **32** showed no activity against TLK2 at the maximum concentration, but there was an increase in TRKC and FLT3 inhibition compared to the other pyridyl analogues synthesized.

There are clear trends observed between pyrrole, 5-position-imidazole, 4-position-imidazole, unsubstituted, 5-position methoxy and acetyl group. The docking of **1, 3, 27** and **28** in the TLK2 protein crystal structure,^11^ provided an insight into these trends (**Figure 3**). The differences between **1** and **3** can be simply explained by the steric clash between the 3,5-dimethylpyrrole on However, the reduced solvation by removal of the additional nitrogen is also contributing to the reduced potency of 3 on TLK2 (**Figure 3A-B**). However, the acetyl substitution of **27** and **28** allowed for a shift in overall binding position that is able to compensate for this steric clash (**Figure 3C-D**) and enables a closing of the gap of potency from >40-fold between **1** and **3** to equipotent between **26** and **28**. We used this knowledge in the design of our next series of ketone-based substituted oxindole analogues.

**Figure 3.**
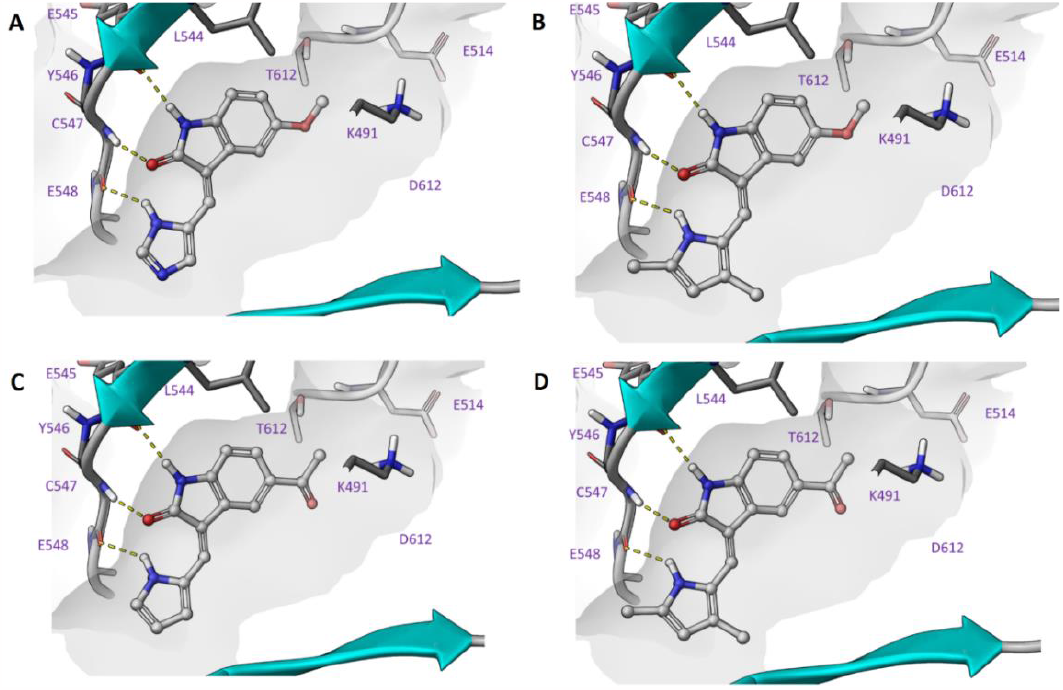
Inhibitor docking of A) **1**, B) **3**, C) **27**, D) **28** in TLK2 (PDB: 5O0Y), demonstrating key hinge binding interactions and space in the hydrophobic pocket.

The (*Z*)-3-((1*H*-pyrrol-2-yl)methylene)-5-acetylindolin-2-one (**27**) was the most potent analogue from the initial matrix array (**Table 1**). The acetyl showed a good trend towards TLK2 inhibition even enabling the 3,5-dimethyl pyrrole substitution **28** to have activity against TLK2 below 1 μM. These two results together with a TLK2 inhibition preference for the pyrrole, unsubstituted 5-position and 4-position imidazole prompted us to focus on the acetyl derivatives (**Table 2**).

**Table 2.** Investigation of derivatives of **27** with pyrrole and imidazole heterocyclic head groups.

| Name | R<sup>1</sup> | X<sup>1</sup> | X<sup>2</sup> | TLK2 IC<sub>50</sub> (nM)<sup>a</sup> | FLT3 (%)<sup>b</sup> | TRKC (%)<sup>b</sup> |
| --- | --- | --- | --- | --- | --- | --- |
| 33 |  | C-H | C-H | 62 | 5.8 | 9.3 |
| 34<sup>c</sup> |  | N | C-H | 140 | 33 | 46 |
| 35 |  | C-H | N | 26 | 30 | 37 |
| 40 |  | C-H | C-H | 220 | 3.6 | 2.5 |
| 41 |  | N | C-H | 230 | 8.9 | 4.3 |
| 42 |  | C-H | N | 250 | 14 | 8.7 |
| 43 |  | C-H | C-H | 870 | 5.9 | 1.9 |
| 44 |  | N | C-H | 700 | 17 | 2.8 |
| 45 |  | C-H | C-H | 340 | 3.5 | 1.0 |
| 46 |  | N | C-H | 340 | 15 | 1.9 |
| 47 |  | C-H | C-H | 100 | 0.8 | 0.9 |
| 48 |  | C-H | N | 340 | 9.6 | 5.2 |

<sup>a</sup>8-point dose response (n=1); <sup>b</sup>percentage inhibition at 0.5 μM (n=2); <sup>c</sup>E:Z ratio: **33** (30:70).<sup>39</sup>

First, we looked at the simple carboxylic acid derivatives **33-35**. The direct replacement of the acetyl to form the carboxylic acid **33** demonstrated a 2-fold improvement in TLK2 inhibition, while the imidazole **34** was equipotent. However, the 2-substituted imidazole **35** showed a 6-fold boost in potency against TLK2 (IC_50_ = 26 nM). Interestingly both imidazole substituted compounds also had good selectivity over the two off-targets FLT3 and TRKC. To pursue the aim of a TLK2 probe quality compound with single digit nanomolar activity against TLK2, we hence investigated larger substituents. The *n*-butyl substituent analogues **40-42** showed a slight decrease in inhibition compared to **27** with flat SAR across the three analogues. The larger phenyl analogues **43-44** showed no improvement with an overall 5-6-fold decrease compared with **27**. Introduction of a 4-fluoro substitution **45-46** mitigated some of the decrease in TLK2 inhibition to 2-3-fold compared with **27** with the same off-target FLT3/TRKC off-target profile. Switching from a phenyl to a furan afforded compounds **47-48** that were near equipotent with **27**. Despite decreased ligand efficiency compared with **27**, these furan analogues **47-48** presented a potential expansion point for further optimization.

The docking of **44, 46** and **47** supported our initial observations with **27** (**Figure 4**). The aromatic groups that are able to interact with the catalytic lysine in **44, 46** and **47** appeared to be preferable to a simple alkyl when attempting to drive potency on TLK2 below IC_50_ = 100 nM (**Figure 4**). Comparing the unsubstituted **44** to the 4-fluoro substituted **46**, we observed a potential halogen bond between the fluorine and a backbone carbonyl which may explain the 3-fold boost in potency against TLK2 (**Figure 4A-B**).^50-52^ However, further analysis of the binding and water network of **35** highlighted untapped potential within a network of interactions in the hydrophobic pocket (**Figure 5**). The carboxylic acid motif of **35** was able to align with the Thr612 which in-turn coordinated a main chain amide, which can be used in the next phase of development. The water network is not disrupted by this series of interactions, but this simulation highlights that the oxindole occupies an optimal hinge binding position.

**Figure 4.**
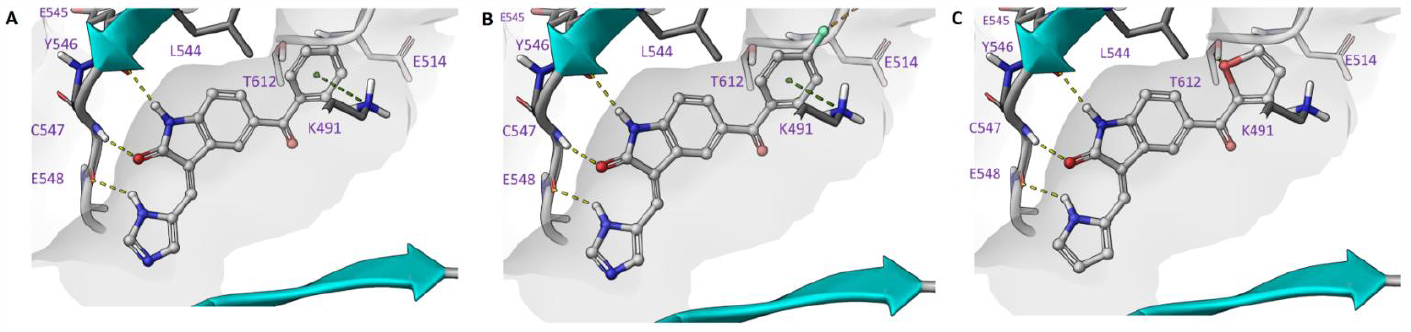
Inhibitor docking comparing of A) **44**, B) **46** and C) **47** in TLK2 (PDB: 5O0Y) all demonstrating key hinge binding interactions and an expansion in the hydrophobic pocket.

**Figure 5.**
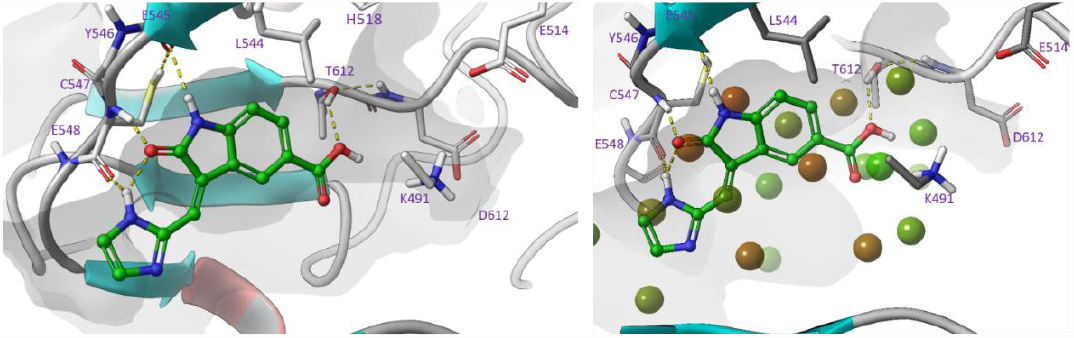
Left: favorable docking pose of compound **35**. Right: visualization of hydration sites from a WaterMap simulation close to docked compound **35**.

The relatively flat SAR of the ketone substitution towards TLK2 inhibition led us to expand our search to other 5-postion substitution patterns including two different sulphonamides (**52-55**) and an *iso*propyl ether (**57-59**) (**Table 3**). Surprisingly, despite the relatively high potency of the acetyl (**27**), the sulphonamides (**60-65**) were inactive on TLK2 at the top concentration. The 5-*iso*propyl ether oxindole showed some potential. The pyrrole analog (**57**) was 3-fold more potent for TLK2 than the 5-methyl ether version (**2**). The imidazole (**58**) was 7-fold less potent on TLK2 than the corresponding 5-methyl ether (**1**). While the 2-subsitituted imidazole (**59**) was the only analog to show an improvement over the methyl counterpart with a 2-fold increase in potency against TLK2.

**Table 3.** Investigation of sulphonamide and ether derivatives.

| Name | R<sup>1</sup> | X<sup>1</sup> | X<sup>2</sup> | TLK2 IC<sub>50</sub> (nM)<sup>a</sup> | FLT3 (%)<sup>b</sup> | TRKC (%)<sup>b</sup> |
| --- | --- | --- | --- | --- | --- | --- |
| 52 |  | C-H | C-H | > 20000 | 31 | 1.6 |
| 53 |  | N | C-H | > 20000 | 55 | 12 |
| 54 |  | C-H | C-H | > 20000 | 27 | 3.0 |
| 55 |  | N | C-H | > 20000 | 62 | 27 |
| 57 |  | C-H | C-H | 560 | 8.3 | 2.8 |
| 58 |  | N | C-H | 1600 | 33 | 11 |
| 59 |  | C-H | N | 160 | 25 | 5.9 |

<sup>a</sup>8-point dose response (n=1); <sup>b</sup>percentage inhibition at 0.5 μM (n=2)

However, these limited improvements meant we switched to the 5-amide substituted oxindoles **65-76** in hope of driving down potency on TLK2 (**Table 4**). We first screened a direct *n*-propyl analog **65** that showed similar potency to **27** with a reduction in off-target potency of FLT3 and TRKC. The addition of a pendent ethyl group to form a pentan-3-yl substitution **66-68** dropped potency by around 10-fold against TLK2 compared to the corresponding *n*-propyl analog **65**. The cyclohexane substitution **69-71** was broadly similar in potency with **27** on TLK2, but with a significant improvement on the off-target profile with a reduction in off-target potency of FLT3 and TRKC compared to **27**. The unsubstituted phenyl **72-74** has an asymmetric structure activity relationship (SAR) for TLK2 with the pyrrole **72** 5-fold weaker than **27**. While the imidazole **73** is 4-fold more potent and both improved off-target selectivity against FLT3 and TRKC. The 2-subsitituted imidazole **74**, is 3-fold less potent than **73** and equipotent to **27**. The furan substitution analogs **75** and **76** were equipotent with **27** on TLK2 but both demonstrated improved off-target selectivity against FLT3 and TRKC.

**Table 4.** Initial investigation of amide derivatives.

| Name | R<sup>1</sup> | X<sup>1</sup> | X<sup>2</sup> | TLK2 IC<sub>50</sub> (nM)<sup>a</sup> | FLT3 (%)<sup>b</sup> | TRKC (%)<sup>b</sup> |
| --- | --- | --- | --- | --- | --- | --- |
| **65** |  | C-H | C-H | 170 | 11 | 4.9 |
| **66** |  | C-H | C-H | 2400 | 25 | 2.3 |
| **67** |  | N | C-H | 2300 | 61 | 6.0 |
| **68** |  | C-H | N | 2500 | 75 | 15 |
| **69** |  | C-H | C-H | 150 | 28 | 11 |
| **70** |  | N | C-H | 100 | 46 | 16 |
| **71** |  | C-H | N | 130 | 58 | 32 |
| **72** |  | C-H | C-H | 820 | 21 | 19 |
| **73** |  | N | C-H | 34 | 44 | 30 |
| **74** |  | C-H | N | 110 | 48 | 33 |
| **75** |  | C-H | C-H | 120 | 30 | 28 |
| **76** |  | C-H | N | 130 | 42 | 33 |

<sup>a</sup>8-point dose response (n=1); <sup>b</sup>percentage inhibition at 0.5 μM (n=2)

The unsubstituted phenyl **72**, is about an order of magnitude less active than close derivatives **73** and **74**. This is likely due to the water network coordination as the 5-membered 1,2-diazole, 1,3-diazole and pyrrole ring system are all pointing towards the solvent exposed region of the TLK2 ATP binding site. The interactions in the hydrophobic pocket region were identical for all three compounds, with the designed interaction of Thr612 and the cation-pi interaction between Lys491 and the phenyl ring observed in each of the docked examples (**Figure 6A-C**). However, we observed through WaterMap that analog **72** was not able to make a stable hydrogen bond network with first solvation shell waters (**Figure 7**), likely leading to weaker binding.^53-56^ The lack of solvent coordination coupled with the ability of **74** to hydrophobically stack with Leu468 (**Figure 7**), which was not possible with **72** and **73**, provided a rational explanation for the results we observed.

**Figure 6.**
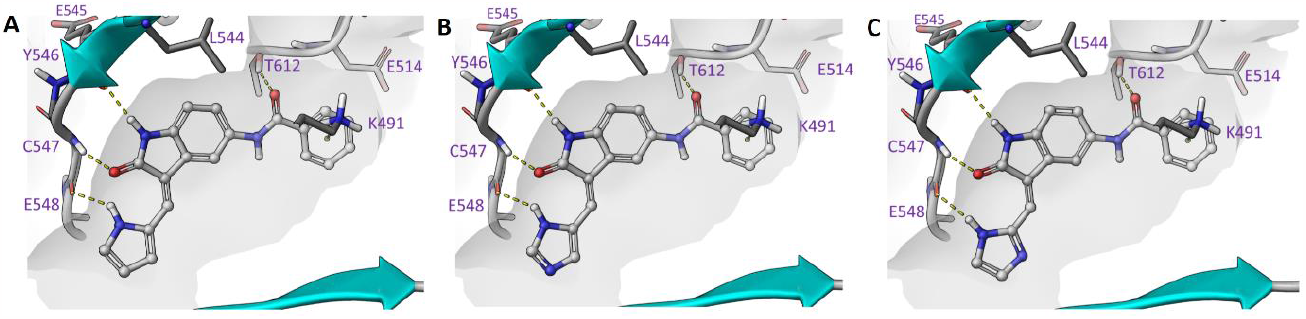
Inhibitor docking comparing A) **72**, B) **73**, C) **74** in TLK2 (PDB: 5O0Y) demonstrating key hinge binding interactions and an expansion in the hydrophobic pocket.

**Figure 7.**
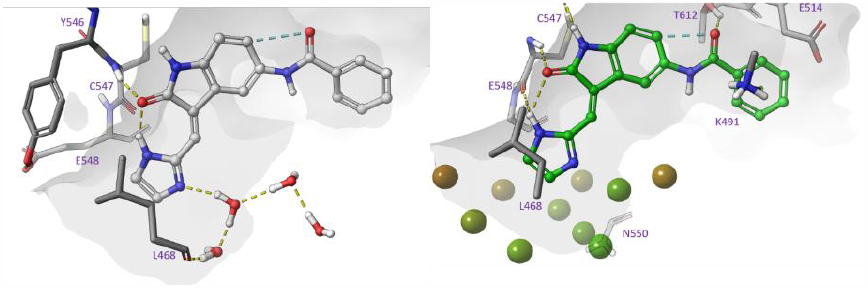
Snapshot of trajectory waters from WaterMap simulation illustrating favourable solvent interactions of **73**. For clarity, waters and hydration sites are shown only at the mouth of the pocket.

To build on the encouraging results of **73** and **74**, we decided to explore the amide derivatives with a series of modifications to the phenyl ring system **88-106** and focused on the imidazole and 2-subsitituted imidazole head group (**Table 5**). First, we probed the 4-position of the phenyl ring, by introducing a nitrile group with an imidazole head group to afford the most potent compound **88** against TLK2 observed so far (IC_50_ = 12 nM); 13-fold more potent than **27**, with good selectivity over FLT3/TRKC. The 3-position nitrile with the imidazole head group **89** afforded 8-fold less potency for TLK2 than **88**, but with slightly improved selectivity over FLT3/TRKC, however the 2-subsitituted imidazole **90** only had a 4-fold drop in potency for TLK2. The 2-position nitrile was not well tolerated with 2-subsitituted imidazole **91** having an IC_50_ over 1 μM on TLK2. The 4-methoxy substitution analogs **92-93** were 7-9-fold less potent on TLK2 than **89**. The 3-methoxy with the imidazole head group **94** was consistent with **92-93**, but the 2-subsitituted imidazole **95** was 28-fold less potent on TLK2 than **94**. In contrast to **91**, the 2-methoxy derivatives **96-97** were potent against TLK2 with the imidazole head group **96** being single digit nanomolar (IC_50_ = 9.1 nM), the most potent compound to date. The 2-subsitituted imidazole derivative **97** was 7-fold less potent on TLK2 but had good selectivity over FLT3/TRKC. The fluorine substitutions around the phenyl ring system **98-102** had relatively flat SAR for TLK2. The 2-position fluoro imidazole **102** was the most potent against TLK2 (IC_50_ = 110 nM) and all analogs **98-102** had good selectivity against FLT3 and TRKC. The larger trifluoromethyl substitution in the 4-postion was partly tolerated with the imidazole head group **103** showing only a 3-fold drop with respect to **102**. However, the 2-subsitituted imidazole **104** showed limited to no activity against TLK2 at the highest concentration tested, potentially due to the inactive geometric *E*-isomer contribution. The 2-position analogs **105-106** were consistent with **98-102**, with equipotency on TLK2 and good selectivity over FLT3. Interestingly, **105-106** both display reduced selectivity over TRKC compared with **103**.

**Table 5.** Advanced investigation of amide derivatives.

| Name | R<sup>1</sup> | X<sup>1</sup> | X<sup>2</sup> | TLK2 IC<sub>50</sub> (nM)<sup>a</sup> | FLT3 (%)<sup>b</sup> | TRKC (%)<sup>b</sup> |
| --- | --- | --- | --- | --- | --- | --- |
| **88<sup>c</sup>** | 4-CN | N | C-H | 12 | 39 | 47 |
| **89<sup>c</sup>** | 3-CN | N | C-H | 95 | 56 | 52 |
| **90** | 3-CN | C-H | N | 48 | 60 | 46 |
| **91** | 2-CN | C-H | N | 4000 | 75 | 33 |
| **92** | 4-OMe | N | C-H | 88 | 36 | 45 |
| **93** | 4-OMe | C-H | N | 110 | 44 | 50 |
| **94** | 3-OMe | N | C-H | 100 | 42 | 48 |
| **95<sup>c</sup>** | 3-OMe | C-H | N | 2800 | 84 | 80 |
| **96** | 2-OMe | N | C-H | 9.1 | 25 | 43 |
| **97** | 2-OMe | C-H | N | 67 | 40 | 34 |
| **98** | 4-F | N | C-H | 190 | 41 | 39 |
| **99** | 4-F | C-H | N | 140 | 46 | 53 |
| **100** | 3-F | N | C-H | 200 | 34 | 33 |
| **101<sup>c</sup>** | 3-F | C-H | N | 490 | 70 | 60 |
| **102** | 2-F | C-H | N | 110 | 26 | 24 |
| **103** | 4-CF<sub>3</sub> | N | C-H | 330 | 58 | 57 |
| **104<sup>c</sup>** | 4-CF<sub>3</sub> | C-H | N | 6100 | 88 | 83 |
| **105<sup>c</sup>** | 2-CF<sub>3</sub> | N | C-H | 400 | 60 | 17 |
| **106<sup>c</sup>** | 2-CF<sub>3</sub> | C-H | N | 190 | 62 | 20 |

<sup>a</sup>8-point dose response (n=1); <sup>b</sup>percentage inhibition at 0.5 μM (n=2); <sup>c</sup>*E*:*Z* ratio: **88** (50:50); **89** (43:57); **95** (37:63); **101** (35:65); **104** (34:66); **105** (38:62); **106** (34:66).<sup>39</sup>

The docking of **73, 88** and **96** provided further evidence of the key interaction at the Thr612 residue with the amide bond carbonyl and the catalytic lysine (Lys491) and afford high potency low double and single digit nanomolar compounds against TLK2 (**Figure 8**). The high affinity was dependent on an aromatic ring system close to the active site but away from the hinge. We looked to exploit these traits to further optimize the analogs towards TLK2 inhibition and potentially increase kinome specificity in the process.

**Figure 8.**
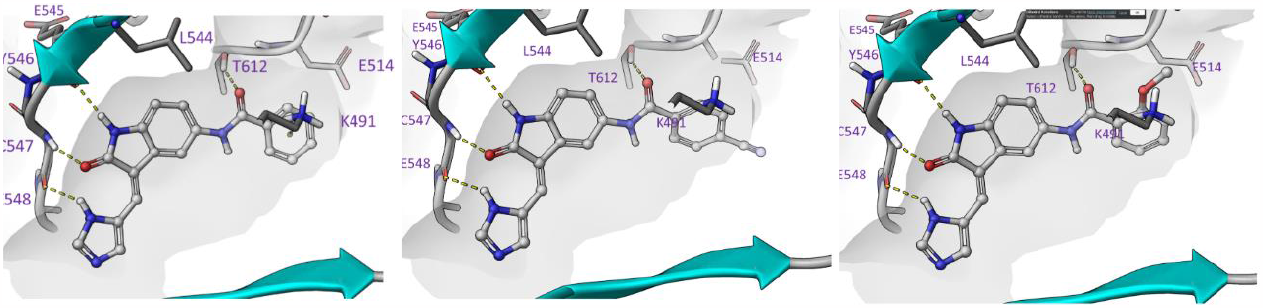
Inhibitor docking in TLK2 (PDB: 5O0Y) comparing A) **73**, B) **88** and C) **96**. All demonstrating key hinge binding interactions and a refined expansion in the hydrophobic pocket.

The recently released solved X-ray structures of TLK2 together with our molecular docking simulations, were also helpful in the design of more effective TLK2 inhibitors.^10^ A 3D-QSAR model was constructed so that the main fields could be overlaid with co-crystallized ligands and verifiable favourable docking poses. We constructed a 3D-QSAR model for TLK2 using the field based QSAR functionality of Schrödinger Maestro (**Table 6** and **7**). The flexible ligand alignments were performed using a Bemis-Mucko method.^52^ The 3D-QSAR model allowed recognition of the largest common scaffold and utilized a favorable docking pose of structurally representative derivatives as alignment templates. Torsion angles of larger substituents, including methoxy groups, were manually adjusted when required. The models were constructed with 80% of compounds defined randomly as training sets with the rest of the compounds as a test set (**Figure 9**). The field values were calculated utilizing a grid spacing approach of 1.0 Å extended 3.0 Å beyond training set limits. Variables with |T-value| < 2 were eliminated from final model building.

**Table 6.** Validation of 3D-QSAR models for TLK2

| # Factors | SD | R<sup>2</sup> | R<sup>2</sup> CV | R<sup>2</sup> Scramble | Stability | F | P | RMSE | Q<sup>2</sup> | Pearsson-r |
| --- | --- | --- | --- | --- | --- | --- | --- | --- | --- | --- |
| 1 | 0.74 | 0.43 | 0.38 | 0.055 | 0.997 | 56.1 | 1.01E-10 | 0.85 | 0.15 | 0.54 |
| 2 | 0.50 | 0.74 | 0.63 | 0.18 | 0.96 | 105.0 | 2.48E-27 | 0.57 | 0.61 | 0.83 |
| 3 | 0.22 | 0.80 | 0.68 | 0.27 | 0.94 | 98.5 | 1.43E-29 | 0.54 | 0.65 | 0.82 |
| 4 | 0.28 | 0.85 | 0.70 | 0.36 | 0.92 | 104.2 | 3.60E-30 | 0.52 | 0.68 | 0.83 |
| 5 | 0.33 | 0.89 | 0.71 | 0.43 | 0.92 | 116.9 | 7.54E-34 | 0.46 | 0.75 | 0.86 |
| 6 | 0.30 | 0.91 | 0.71 | 0.48 | 0.88 | 122.9 | 3.73E-38 | 0.47 | 0.74 | 0.86 |

**Table 7.**
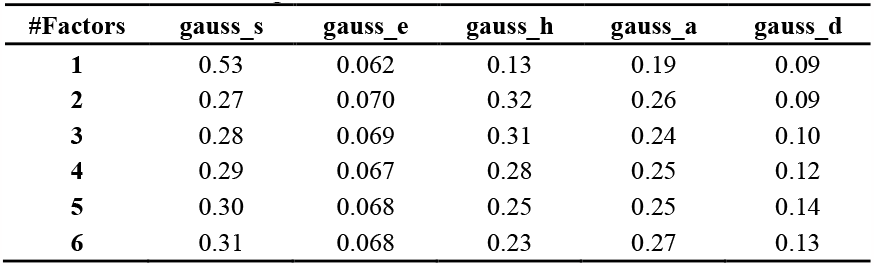
Regression Statistics for TLK2 QSAR model

**Figure 9.**
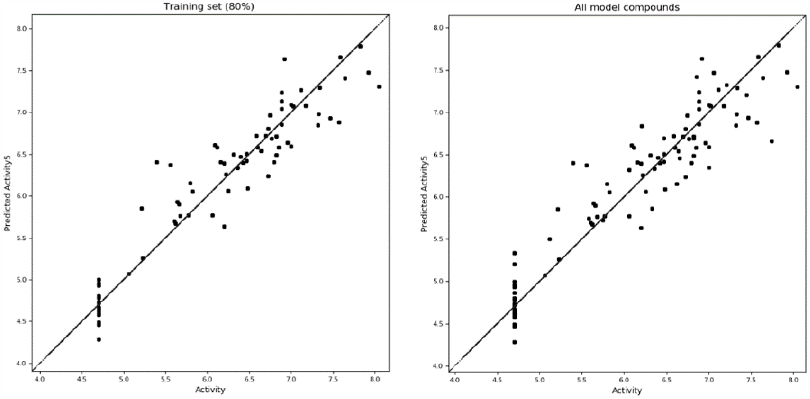
Scatter plots of field-based 3D-QSAR models showing predicted versus measured binding affinities (nM) of active compounds printed at logarithmic scale with the TLK2 training set (left) and TLK2 all compounds (right).

The R-squared value of the TLK2 model with all four partial least squares regression (PLS) components were 0.75. The model had above average internal predictivities with a Q-square value of 0.75, which was determined by leave-one out (LOO) cross-validation. The stability of the model was 0.92 with four (PLS) components having demonstrated a robust predictivity of structure to activity output.

The shape of the oxindole series had a good alignment in the TLK2 ATP binding pocket with a strong interaction with hinge residues with semi-optimized back pocket QSAR fields (**Figure 10A-F**). The pyrrole/imidazole head group assisted the hinge interacting with an additional anchor point. The oxindole scaffold was able to form two direct hydrogen bond interactions *via* the oxindole amide functionality to the main chain amide of Tyr546 and Cys547. The N-H of the pyrrole/imidazole was then able to form an interaction with Glu548. The optimized amide derivatives were also able to interact *via* the carbonyl of the amide with Thr612 and had a cation-pi interaction with Lys491 forming with the pendent heterocycle. The internal conformational restriction of the oxindole scaffold between the carbonyl and amine of the pendant heterocycle was also assisting in the optimization of TLK2 binding, not only minimizing the entropic loss associated flexibility but also allowed the ligand to adopt a preferred conformation for binding.^39,57-58^

**Figure 10.**
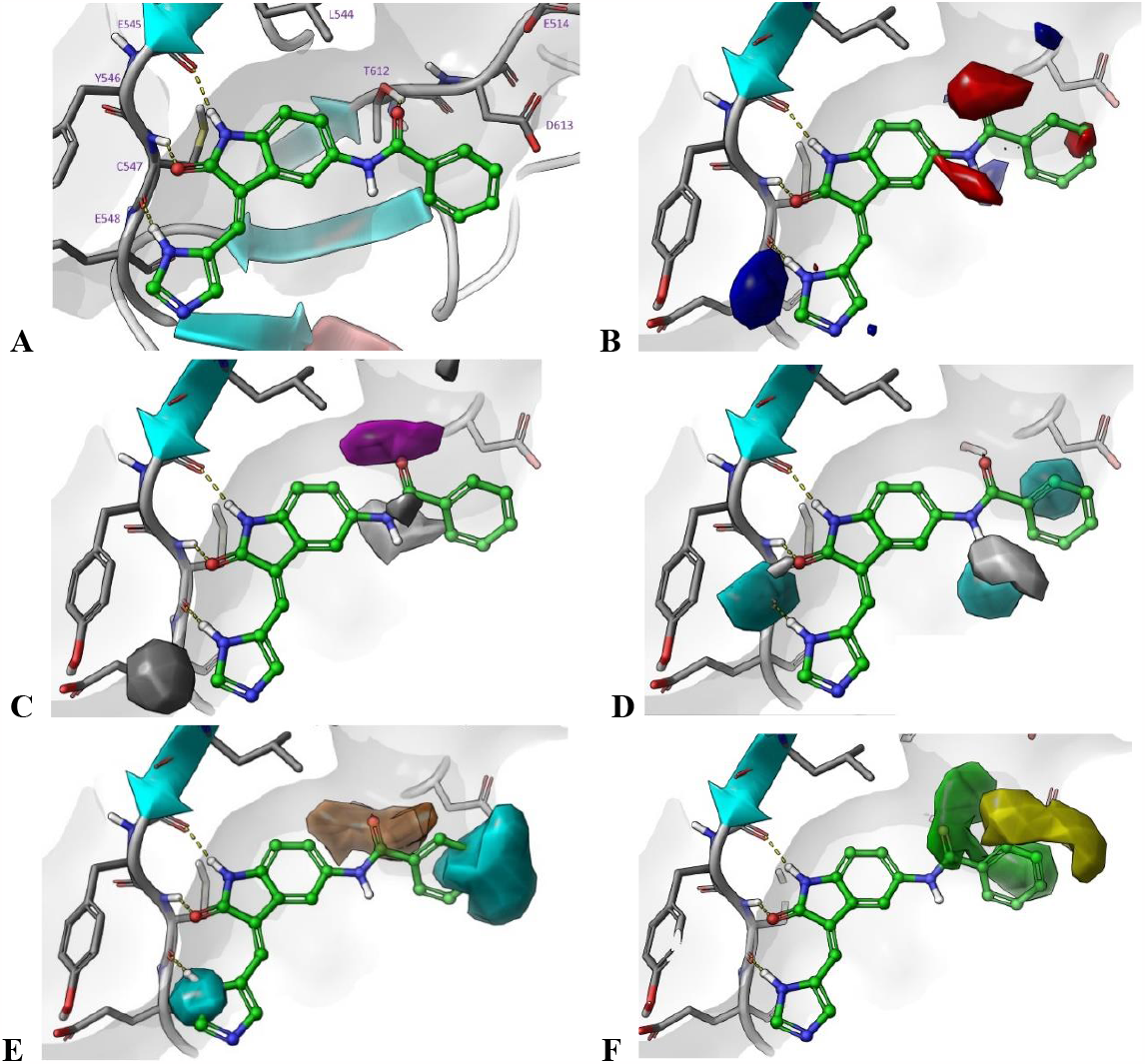
QSAR modelling on **73** template: A) Standard docking of **73** in TLK2 (PDB:5O0Y); B) Electrostatics (blue = positive and red = negative); C) hydrogen bonding acceptor fields (magenta = positive and grey = negative); D) hydrogen bonding donor fields (cyan = positive and grey = negative); E) hydrophobic fields (orange = positive and cyan = negative); F) Steric fields (green = positive and yellow = negative).

We further explored the rigidity within the oxindole core with the solving of a series of small molecule crystal structures **1, 2, 15, 19, 72**, and **74** (**Figure 11, Table S3**) and **4, 9, 11, 13** and **16** (**Figure S1, Table S3**). Structures **1, 2, 16** and **19** were planar due to intramolecular hydrogen bonding between the pyrrole **15** and imidazole **1** and **2** ring and the oxindole carbonyl oxygen atom (**Figure 11**). The steric hindrance of the pyridine substituent of **19** and lack of hydrogen bond donor resulted in a deviation from planarity for this structure. Whilst the methylene pyrrole/imidazole rings were planar with the oxindole moiety, the benzamide substituents of **72** and **74** were not (**Figure 11**). The methylene imidazole substituted structures **1** and **15** included a hydrogen bonded dimer between the amine of the imidazole and the oxindole carbonyl instead except **74** which formed flat 1D tapes along the crystallographic *-ac* plane *via* oxindole amine, imidazole imine H-bonding. These tapes formed H-bonds to those above and below *via* hydrogen bonds between the amide functionalities. The methylene pyrazole substituted structures showed no distinct packing motifs with **2** forming interleaved, corrugated tapes *via* oxindole amine, imidazole imine H-bonding and **16** forming a H-bonded, tetramer structure comprising two crystallographically independent molecules.

**Figure 11.**
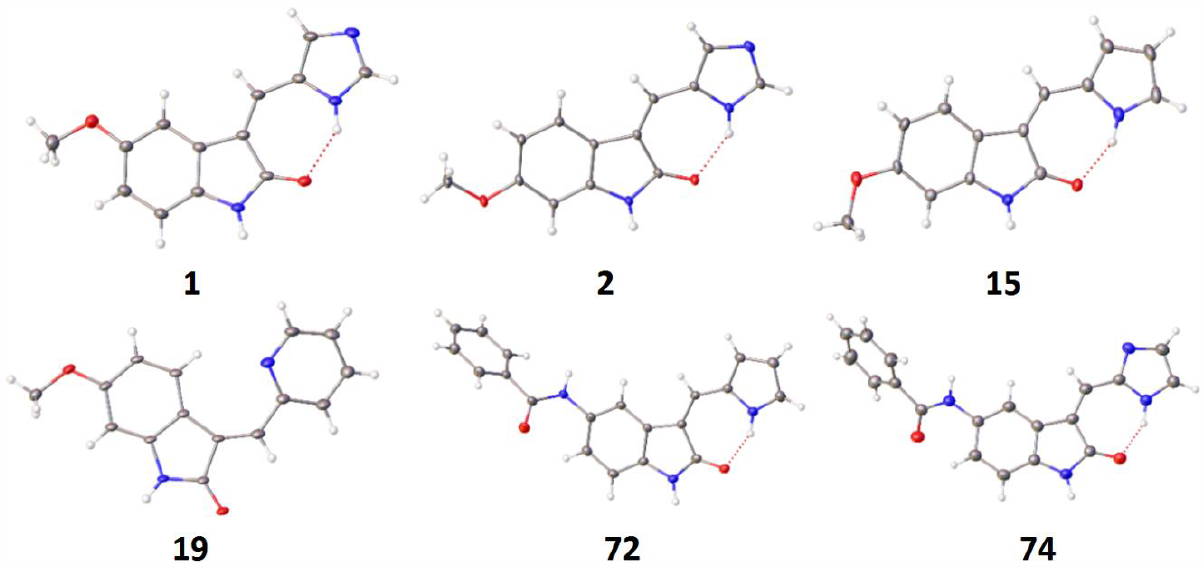
Small molecule crystal structures of **1, 2, 15, 19, 72** and **74**.

A series of oxindole derivatives were designed based on the quantitative structure-activity relationships (QSAR) QSAR model utilizing prior SAR knowledge and the small molecule crystal structure information. These were synthesized and tested using the same protocol as described above, with the addition of TLK1 screened at a 0.5 μM concentration, consistent with previous reports.^23^

We synthesized the series of predicted analogs utilizing similar conditions as before (**Scheme 1**) with a few alterations in starting materials. The amide intermediates **107-129** were afforded by a routine HATU coupling to the respective 5-aminooxindole in good yields (57-84 %). The amide was then treated with the previous Knoevenagel conditions with the respective aldehyde to afford the condensation products **117-129** in a range of yields (21-71 %) (**Scheme 2**).

**Scheme 2.**
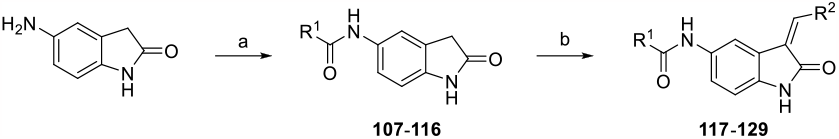
Synthetic route to prepare predicted oxindole analogs - *reagents and conditions* a) R^1^COCl, EtN(CH(CH_3_)_2_)_2_, THF, rt, 12 h; b) R^2^CHO, piperidine, EtOH, reflux, 4-24 h.

The QSAR modelling suggested that the phenyl ring could be reduced to a 5-member ring and coupled with earlier knowledge of the furan **75** and **76**, so we synthesized the thiophenes **117-121** (**Table 8**). The activities observed were broadly in line with the QSAR model. The unsubstituted thiophene with the imidazole head group **117** was potent on TLK2 (IC_50_ = 23 nM) but lacked selectivity over FLT3/TRKC, the 2-subtitued imidazole **118** was equipotent but slightly more selective against FLT3/TRKC. The introduction of a 4-methoxy group onto the thiophene **119** reduced potency for TLK2 by 3-fold compared to **117**. However, the 4-ethoxy group **120** was equipotent with **117** with equivalent selectivity over FLT3/TRKC. Interestingly **120** was the only compound to show any significant activity on TLK1 in the initial screening (54% inhibition at 1 µM). The final compound in this series, the 4-fluoro **121** was 8-fold less potent against TLK2 compared to **117**.

**Table 8.**
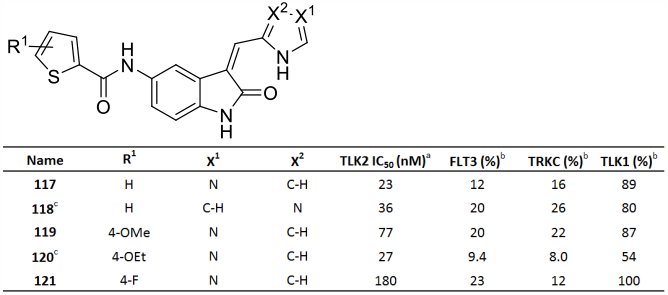
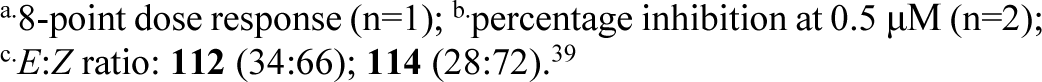
Results of initial set of QSAR predicted compounds.

| Name | $R^1$ | $X^1$ | $X^2$ | | | | |
| --- | --- | --- | --- | --- | --- | --- | --- |
| | | | | TLK2 $IC_{50}$ (nM)<sup>a</sup> | FLT3 (%)<sup>b</sup> | TRKC (%)<sup>b</sup> | TLK1 (%)<sup>b</sup> |
| **117** | H | N | C-H | 23 | 12 | 16 | 89 |
| **118<sup>c</sup>** | H | C-H | N | 36 | 20 | 26 | 80 |
| **119** | 4-OMe | N | C-H | 77 | 20 | 22 | 87 |
| **120<sup>c</sup>** | 4-OEt | N | C-H | 27 | 9.4 | 8.0 | 54 |
| **121** | 4-F | N | C-H | 180 | 23 | 12 | 100 |

<sup>a</sup>8-point dose response (n=1); <sup>b</sup>percentage inhibition at 0.5 μM (n=2); <sup>c</sup>E:Z ratio: **112** (34:66); **114** (28:72).<sup>39</sup>

We then optimized these initial compounds to afford a series of more advanced amide derivatives **122-127** (**Table 9**). The fused thiophene, **122** was equipotent with **117**, with no difference in selectivity. This larger 5,5-system was tolerated, so an expansion of the phenyl substitution patterns was synthesized to include 3-trifluoromethyl, 5-fluoro **123** and the fused 2,3-catacol **124**, unfortunately these analogs both lost significant activities against TLK2 compared with **117**. We also looked at smaller 5-member rings, thiazole **125**, oxazole **126** and isothiazole **127** afforded potent compounds against TLK2 with the oxazole approaching towards single digit nanomolar against TLK2.

**Table 9.**
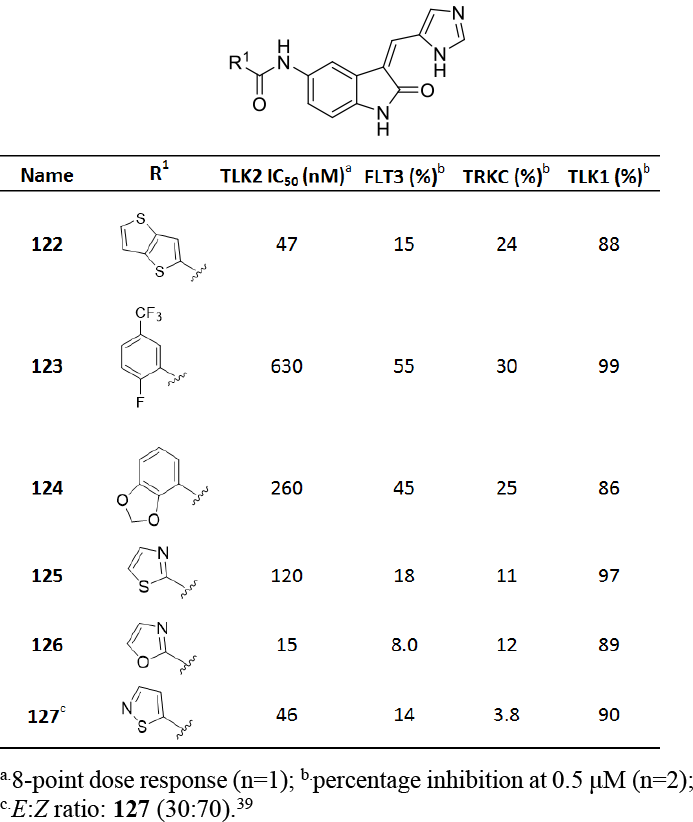
Optimization of compounds based on QSAR.

|  |  |  |  |  |  |
| --- | --- | --- | --- | --- | --- |
| Name | R<sup>1</sup> | TLK2 IC<sub>50</sub> (nM)<sup>a</sup> | FLT3 (%)<sup>b</sup> | TRKC (%)<sup>b</sup> | TLK1 (%)<sup>b</sup> |
| **122** |  | 47 | 15 | 24 | 88 |
| **123** |  | 630 | 55 | 30 | 99 |
| **124** |  | 260 | 45 | 25 | 86 |
| **125** |  | 120 | 18 | 11 | 97 |
| **126** |  | 15 | 8.0 | 12 | 89 |
| **127<sup>c</sup>** |  | 46 | 14 | 3.8 | 90 |

<sup>a</sup>8-point dose response (n=1); <sup>b</sup>percentage inhibition at 0.5 μM (n=2); <sup>c</sup>E:Z ratio: **127** (30:70).<sup>39</sup>

In a further effort to validate the model and test the limits of the two additional compounds **128** and **129** were synthesised (**Table 10**). These were based on the cores structure of **73** and tested the predictive power of the model. The addition of the 2-methyl (**128**) to the imidazole head group afforded an 8-fold increase in potency against TLK2 with no loss of selectivity against FLT3 and TRKC. The model was accurate for the pyrazole **129**, the introduction of the extra nitrogen led to an over 15-fold drop in TLK2 inhibition, consistent with previous SAR in this series.

**Table 10.**
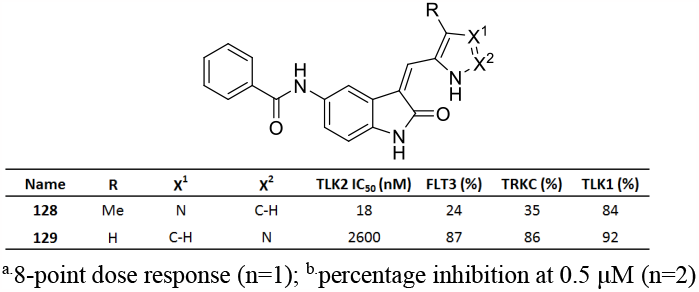
Optimization of compounds based on QSAR.

|  |  |  |  |  |  |  |  |
| --- | --- | --- | --- | --- | --- | --- | --- |
| Name | R | X<sup>1</sup> | X<sup>2</sup> | TLK2 IC<sub>50</sub> (nM) | FLT3 (%) | TRKC (%) | TLK1 (%) |
| **128** | Me | N | C-H | 18 | 24 | 35 | 84 |
| **129** | H | C-H | N | 2600 | 87 | 86 | 92 |

<sup>a</sup>8-point dose response (n=1); <sup>b</sup>percentage inhibition at 0.5 μM (n=2)

Aligning the results of the prediction with the experimental results, we found a good concordance (**Table 11**).

**Table 11.** Compound results based on QSAR prediction.

| Compound | TLK2 Inhibition Activity (nM) |  |  |  |
| --- | --- | --- | --- | --- |
|  | Predicted | Found | Log Predicted | log Actual |
| **117** | 51 | 23 | 7.29 | 7.64 |
| **118** | 56 | 36 | 7.25 | 7.44 |
| **119** | 50 | 77 | 7.30 | 7.11 |
| **120** | 370 | 27 | 6.44 | 7.57 |
| **121** | 68 | 180 | 7.17 | 6.74 |
| **122** | 180 | 47 | 6.75 | 7.33 |
| **123** | 1800 | 630 | 5.76 | 6.20 |
| **124** | 40 | 260 | 7.40 | 6.59 |
| **125** | 8 | 120 | 8.08 | 6.92 |
| **126** | 8 | 15 | 8.12 | 7.82 |
| **127** | 37 | 46 | 7.44 | 7.34 |
| **128** | 180 | 18 | 6.74 | 7.74 |
| **129** | 960 | 2600 | 6.02 | 5.59 |

In addition to assessing the selectivity profile on the two near off-targets FLT3 and TRKC; we also investigated the wider kinome selectivity profile of ten of the most promising analogs using KinomeSCAN^®^ at 1 μM across over 400 kinases (**Table 12, Figure 12**).^20-22^ The original starting point (**1**) was moderately selective across the kinome hitting 15 kinases over 90%. The 5-acetyl analog **27**, was the most promiscuous analog with 62 kinases over 90%, possibly due to the small flexible ketone orientated favourably to towards the common catalytic lysine motif (**Figure 3D**). Interestingly, **1, 27** and **44** had similar hit kinases above 90%. Common kinases included MST2, CDKL2, ERK8, TRKA, JAK3(JH1domain-catalytic) and TNIK. The amide substitution in the 5-position not only increased selectivity across the kinome but also changed the kinases inhibited. Compounds **72-74** consistently hit MYLK4, BIKE, GRK4, SRPK3 above 90%, another four PRP4, AAK1, PIP5K1A, SRPK2 are revealed as consistent hits if the threshold is decreased to 80% inhibition, in addition to other kinases (see supporting information). The most narrow spectrum inhibitor across the kinome tested in the KinomeSCAN^®^ assay was **97** which hit only 7 kinases above 90%; including BIKE, DRAK2, JAK1(JH2domain-pseudokinase), MAST1, PIP5K1A, GRK4, and TYK2 (JH2domain-pseudokinase). The selectivity of the designed thiophene amide **111** was moderate with 15 kinases inhibited. The addition of a methyl **128** onto the 5-position of the imidazole head group of **73** changed the kinome profile subtly at 90% inhibition, with inhibition reduced on TYK2(JH2domain-pseudokinase), AAK1 but also increased on DRAK2, PIP5K1A, RIOK1, MAST1, LKB1.

**Table 12.**
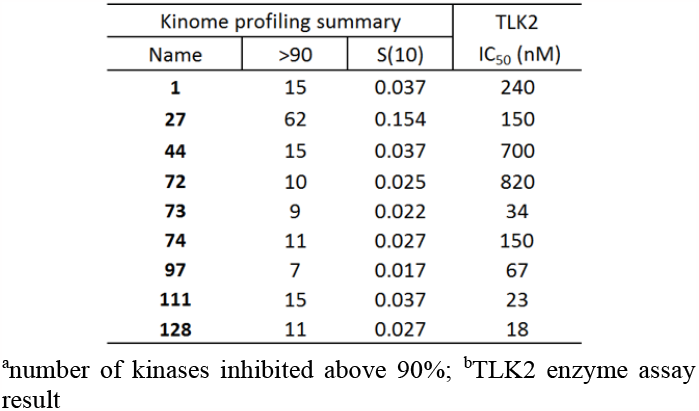
Kinome selectivity profiling

| Kinome profiling summary |  |  | TLK2 IC<sub>50</sub> (nM) |
| --- | --- | --- | --- |
| Name | >90 | S(10) |  |
| **1** | 15 | 0.037 | 240 |
| **27** | 62 | 0.154 | 150 |
| **44** | 15 | 0.037 | 700 |
| **72** | 10 | 0.025 | 820 |
| **73** | 9 | 0.022 | 34 |
| **74** | 11 | 0.027 | 150 |
| **97** | 7 | 0.017 | 67 |
| **111** | 15 | 0.037 | 23 |
| **128** | 11 | 0.027 | 18 |

<sup>a</sup>number of kinases inhibited above 90%; <sup>b</sup>TLK2 enzyme assay result

**Figure 12.**
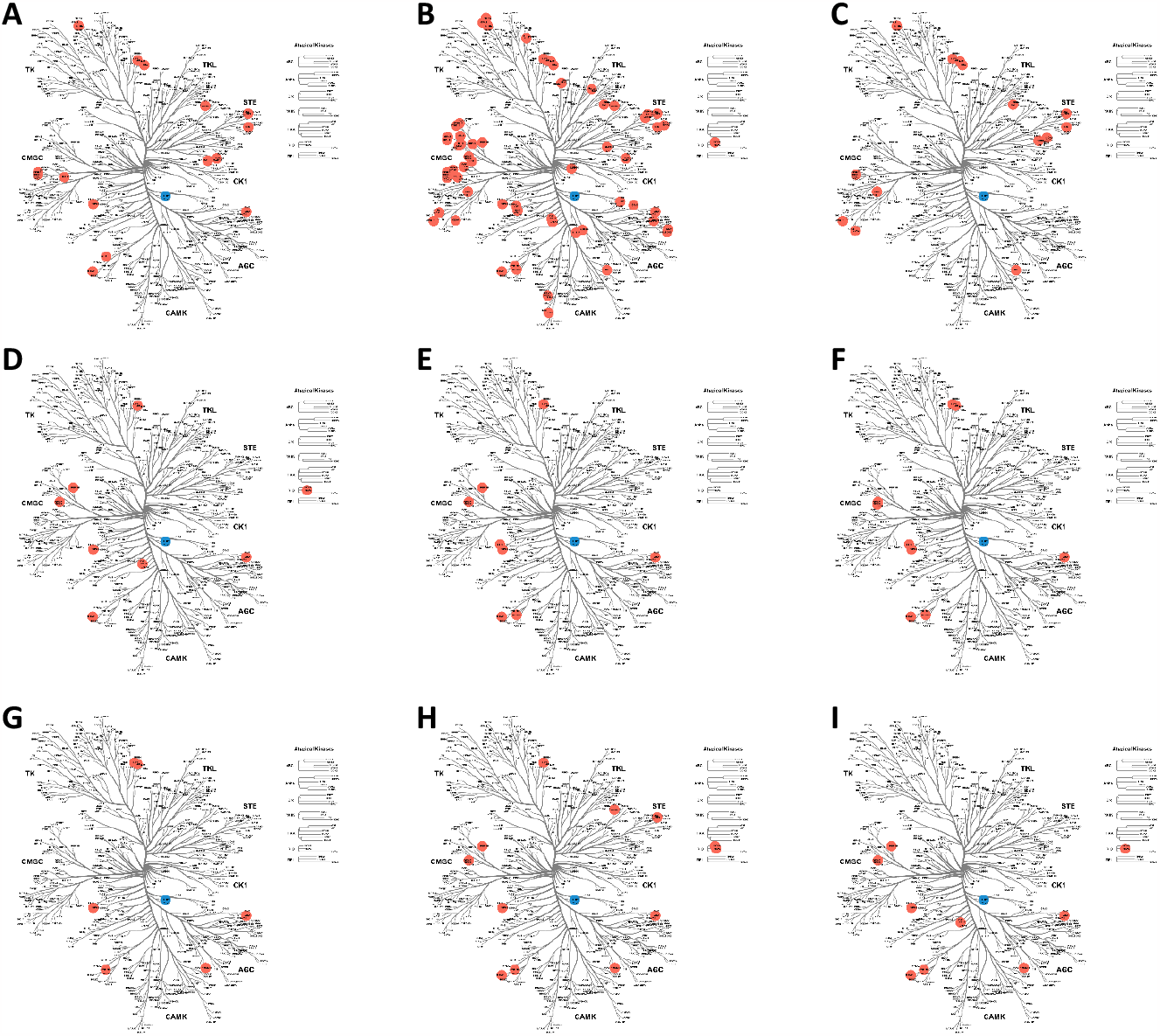
Kinome tree image of KinomeScan data for A) **1**, B) **27**, C) **44**, D) **72**, E) **73**, F) **74**, G) **97**, H) **111**, I) **128**

Finally, we screened the most promising compounds **97, 117, 128**, the negative control compound **129**, along with the starting point compound **1** in an orthogonal assay measuring the conversion of ATP to ADP.^59-61^ The results were consistent with both the weak and potent TLK2 binding of the five compounds in the enzyme assay (**Figure 13** and **Table 13**). The potent TLK2 activities of **97, 117** and **128** in addition to their narrow spectrum kinome selectivity makes them attractive starting points for further optimization.

**Table 13.** TLK2 Functional Inhibition Kinase-Glo Results

| Name | TLK2 IC<sub>50</sub> (nM) |  |
| --- | --- | --- |
|  | Enzyme | Kinase Glo |
| **1** | 240 | 280 |
| **97** | 67 | 450 |
| **117** | 15 | 97 |
| **128** | 18 | 250 |
| **129** | 2700 | 7800 |

**Figure 13.**
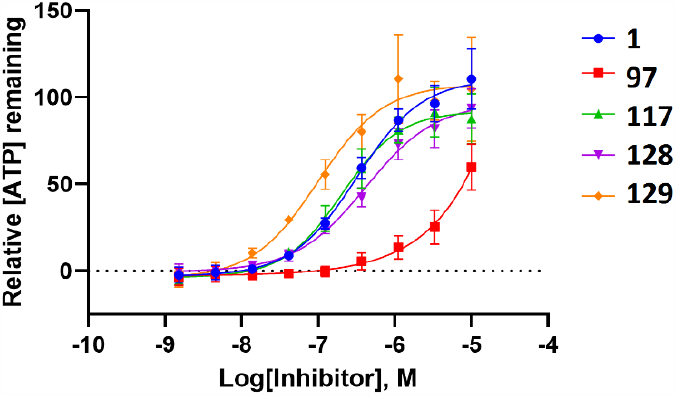
TLK2 Kinase-Glo curves (n=3) for **1, 97, 117, 128, 129**.

## 3. Discussion

TLK2 represents an attractive target for therapeutic development in a variety of cancer types and in “shock-and-kill” approaches for antiviral therapies. The published TLK2 inhibitors to date only offer compounds with moderate potency and poor kinome selectivity.^18,62-63^ In the present study, we demonstrate optimization of a chemotype for increased potency and selectivity for the clinically important TLK2 kinase. Since we achieved selectivity over the closest human paralog, TLK1, this work also provides important tool compounds for better characterization of the specific cellular functions of TLK2.^64-65^

The original target for the oxindole scaffold was receptor tyrosine kinases namely VEGFR and PDGFR.^66^ However, the scaffold demonstrated varying affinities for a large portion of the kinome.^20-23^ There have been a number of priority cancer targets including TrkA,^67^ CDK2,^68-70^ PDK1,^71^ and Src,^72^ among others,^73^ all inhibited by the oxindole scaffold. While the primary kinase targets were well defined, the oxindole chemotype has historically been considered as a broad-spectrum inhibitor chemotype. However, a LRRK2 program at Novartis by Troxler *et al*,^74^ suggested that the replacement of fluorine in the 5-position with a methoxy significantly improved kinome wide selectivity with the oxindole scaffold. This was supported by the narrow spectrum of the initial 5-methoxy substituted hit compound SU9516, when compared to the matched pair oxindoles GW5060033X and Sunitinib.^20-23^

The two oxindole matched pairs in the literature (**Figure 1**) demonstrated a tractable affinity to TLK2.^20-23^ We found several key structural features that could be tuned within the scaffold including: 1) The electronics of the oxindole head group and the internal hydrogen bond; 2) the electronics and sterics of the amide substitution pattern and 3) formation of the key Lys491 interaction with the amide linker.

The use of conformational restriction has been an effective tool in drug discovery.^75^ This methodology has been utilized to orient ATP competitive inhibitors in the preferred conformation. This can be used to not only increase potency, but also to enhance selectivity over other kinases/targets. There are several examples within kinase inhibitor design including Cyclin G-Associated Kinase (GAK),^55,76^ Spleen tyrosine kinase (Syk)^77^ and Cardiac troponin I-interacting kinase (TNNI3K)^78-79^ where addition or removal of a single nitrogen to alter the inhibitor conformation resulted in an increase by 60-160-fold on target.^80^ Another method employed in an EGFR inhibitor program involved fixing the pendent substituents with hydrogen bonds between an alcohol and a carbonyl.^56^ There are a number of other non-kinase examples including the development of a human histamine H4 receptor (hH4R) inhibitor agonist,^81^ and design of a selective CB2 cannabinoid receptor agonist.^82^

The adding or removing of a nitrogen has been proven to have a significant effect on activities across different systems, not only by altered conformation, but also in reduced de-solvation energy potential and as a hydrogen bond donor/acceptor.^76^ The oxindole scaffold has multiple potential contact points within the TLK2 ATP binding site. However, we found that that the most crucial nitrogen was on the pendant head group contained within the pyrrole/imidazole and formed not only an interaction at the hinge with Glu548 but also, an internal hydrogen bond with the carbonyl of the oxindole. This internal hydrogen bond pre-organizes the orientation of the molecule as an optimal ATP memetic (**Figure 14**). The solvent exposed shell is a key modulator of activity, with differing profiles depending on the nitrogen location on the ‘head group’ 5-member ring. We can rationally explain these differences by modelling the first water network solvent shell.^55,84-85^ Comparing **Figure 14D** to **Figure 14E-F** using WaterMap, both imidazoles are able to coordinate using the second nitrogen, while the pyrrole is not leading to a 10-20 fold drop in potency on TLK2.

**Figure 14.**
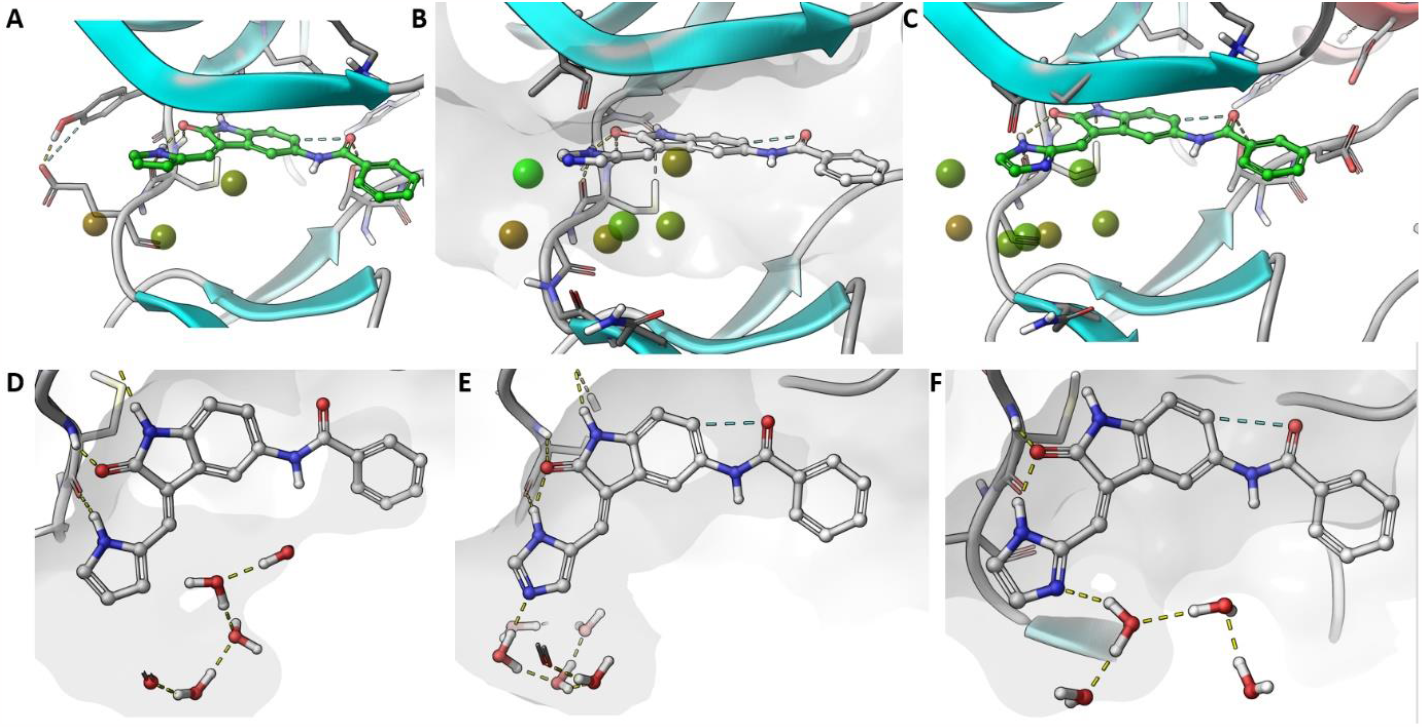
Representative oxindoles with three key head groups showing first shell solvent front interactions using WaterMap A) **72**, B) **73**, C) **74**, D) **72**, E), **73** and F) **74** in TLK2 (PDB: 5O0Y). Waters are presented with two different representations balls A-C (green - low energy and red – higher energy) and full formulae D-F.

**Table 8**. Toxicity profiling of different oxygen substituted anilines based on the scaffold of analogue **72** (**101-112**).^a,b^

In addition to the conformation and head group nitrogen and oxindole hinge interactions our modelling highlights that to achieve a high affinity compound, a favourable fit into the hydrophobic back pocket is required. The key hydrogen bond interaction to the catalytic lysine (Lys491) affords a considerable boost in potency. The challenge then is to maintain selectivity across the kinome, this is achieved in part by a series of cationic-pi interactions also donated by Lys491, but also favourable fine-tuned π–π stacking interactions. This is demonstrated with a highlighted section of the 5-position substitution and the possibility to form a cationic-π interaction with Lys491. The small molecule crystal structures (**Figure 4 & Table S10**) support the theoretical observations with the TLK2 ATP binding site.

## Conclusions

Oxindole based inhibitors are often considered to be promiscuous across the kinome, this mainly derived from several well-known broad-spectrum inhibitors including sunitinib and GW506033X.^86^ Here we demonstrate fine-tuning and optimisation to derive a narrow spectrum oxindole based TLK2 inhibitor, utilizing cutting edge modelling to prospectively design inhibitors. The result was compound **126** and **128** that were 10 to 15-fold more potent than the starting point **1** with maintenance of kinome selectivity and selectivity over the sub-family member TLK1. The QSAR and water network modelling provided new insights into the development of selective chemical probes for TLK2 and beyond. The comprehensive data sets also afford a more complete level of knowledge of the oxindole previously unavailable to the medicinal chemist’s toolbox.

## Experimental

### Molecular Modelling

Ligand preparation: Structures of small molecules were parametrized and minimized using the LigPrep module of Schrodinger 2019-4 suite employing OPLS3e force field.

Protein preparation: The TLK2 kinase a crystal structure (PDB:5O0Y) co-crystallized with substrate analog (phosphothiophosphoric acid-adenylate ester) was downloaded from RCSB-databanks. The structure was H-bond optimized and minimized using standard protein preparation procedure of Schrödinger suite.

TLK2 Molecular docking: The ligand docking was performed using SP settings of Schrodinger docking protocol with softened vdw potential (scaling 0.6). The initial docking of resulted quite poor convergence, thus oxindole-indole scaffold can bind in several conformations. To ensure correct docking mode of the oxindole (2-indolone) scaffold to kinase folds was confirmed using LPDB module of the Schrödinger package and oxindole as a query. After visual inspection of PDB:2X2K, 2X2M, 4QMO and 1PF8. To further improve convergence of docking poses a hydrogen bond constraint to mainchain NH of hinge residue was required, as experimentally observed in the case of other oxindole containing scaffolds (NH of C507 in the case of TLK2). The grid box was centered using coordinate center of the core structure of corresponding x-ray ligand as template. Graphical illustrations were generated using, Maestro software of Schrödinger. Flexible ligand alignments were performed using a Bemis-Mucko method, enabling recognition of the largest common scaffolds using a favorable docking pose of structurally representative derivative as alignment template (**73**). Torsion angles of larger substituents, such as methoxy groups, were manually adjusted to fit template structures when needed.

TLK2 3D-QSAR model: The model for the TLK2 kinase was constructed using the field based QSAR functionality of Schrödinger Maestro 2019-4. The models were constructed by defining 80% of compounds randomly as training sets and rest of the compounds were considered as test set. Grid spacing of 1.0 Å extended 3.0 beyond training set limits was used to the calculate field values. Subsequently, predicted versus measured affinities were plotted to identify outliers. The statistical metrics of models: R-square values with four PLS components were 0.79. The model shows acceptable internal predictivity (Q-square value 0.70), determined by leave-one-out (LOO) cross-validation. Stability value above 0.9 with four partial least squares regression (PLS) components let us to suggest that the quality and predictivity of the model inside the studied applicability domain was robust.

### Crystallography

Each compound had a suitable crystal was selected and mounted on a MITIGEN holder (MiTeGen, Ithaca, NY, USA) in perfluoroether oil on a Rigaku FRE+ diffractometer equipped with a Mo target, VHF Varimax confocal mirrors, an AFC12 goniometer and HyPix 6000 detector (**1, 2, 15, 19, 72** and **74**) or a Rigaku 007HF diffractometer equipped with a Cu target, HF Varimax confocal mirrors, an AFC11 goniometer and HyPix 6000 detector (**15, 74**). The crystal was kept at a steady *T* = 100(2) K during data collection. CrysAlisPro^87^ was used to record images, process all data and apply empirical absorption corrections. Unit cell parameters were refined against all data. The structures were solved by the ShelXT^88^ structure solution program using the using dual methods solution method and using Olex2^89^ as the graphical interface. The models were refined with version 2018/3 of ShelXL,^88^ using full matrix least squares minimization on *F*^2^ minimization. All non-hydrogen atoms were refined anisotropically. Hydrogen atom positions were calculated geometrically and refined using the riding model, except for those bonded to N-atoms which were located in the difference map and refined with a riding model. Structure **72** crystallised as an ethanol solvate. The data for **72** were processed as a 2-component non-merohedral twin (rotation (UB1, UB2) = 179.9982° around [0.00 0.71 0.71] (reciprocal) or [-0.39 0.85 0.35] (direct) lattice vectors). CCDC deposit numbers: 2309069-2309077, see supporting information.

### Chemistry

#### General chemistry

All reactions were performed using flame-dried round-bottomed flasks or reaction vessels. Where appropriate, reactions were carried out under an inert atmosphere of nitrogen with dry solvents, unless otherwise stated. Yields refer to chromatographically and spectroscopically pure isolated yields. Reagents were purchased at the highest commercial quality and used without further purification. Reactions were monitored by thin-layer chromatography carried out on 0.25-mm. Merck silica gel plates (60_F-254_) using ultraviolet light as a visualizing agent. NMR spectra were recorded on a Varian Inova 400 (Varian, Palo Alto, CA, USA) or Inova 500 spectrometer (Varian, Palo Alto, CA, USA) and calibrated using residual undeuterated solvent as an internal reference (CDCl_3_: ^1^H NMR = 7.26, ^13^C NMR = 77.16). The following abbreviations or combinations thereof were used to explain the multiplicities observed: s = singlet, d = doublet, t = triplet, q = quartet, m = multiplet, br = broad. Liquid chromatography (LC) and high resolution mass spectra (HRMS) were recorded on a ThermoFisher hybrid LTQ FT (ICR 7T) (ThermoFisher, Waltham, MA USA). Compound spectra of **1-129** and mass spectrometry method are included in the Supplementary Materials. All compounds were >98% pure by ^1^H/^13^C NMR and LCMS.

**3-[(3,5-Dimethyl-1H-pyrrol-2-yl)methylene]-1,3-dihydro-2H-indol-2-one** (SU5416/Semaxanib) (**10**) was consistent with commercial product and previous report.^90^ **2-oxoindoline-5-sulfonyl chloride** (**49**) was prepared consistent with previous report.^45^ **5-*iso*propoxyindolin-2-one** (**56**) was prepared consistent with previous report.^74^

### General Procedures

#### Procedure A. Knoevenagel condensation

The oxindole (1.0 equiv), aldehyde (1.1 equiv), piperidine (2.4 equiv) were suspended in ethanol at reflux overnight. The solvent was removed and the resulting solid re-suspended DCM/EtOAc 1/1 and flushed through a plug of silica (DCM/EtOAc 1/1). The product was the purified by silica gel flash column chromatography (hexane:EtOAc) to afford the title compound.

#### Procedure B. Friedel-Crafts acylation

To a suspension of AlCl_3_ (6.0 equiv) in CH_2_Cl_2_ was added the appropriate acyl chloride (3.0 equiv) and the slurry was stirred at room temperature for 30 min, followed by the addition of oxindole (1.0 equiv) in CH_2_Cl_2_. The resulting mixture was stirred at room temperature for 2 days. Upon completion, the solution was cooled to 0 °C before quenching the reaction with ice/water, followed extraction of the organics with CH_2_Cl_2_. The combined organics were washed with saturated solution of NaHCO_3_, dried over Na_2_SO_3_ and the solvent was removed by rotary evaporation before purification by silica gel flash column chromatography (cyclohexane:EtOAc 9:1 to 1:1) to afford the title compound.

#### Procedure C. Friedel-Crafts acylation

The appropriate acyl chloride (3.0 equiv) was added to grinded AlCl_3_ (5.0 equiv) and the slurry was heated to 150 °C, then oxindole (1.0 equiv) was added portionwise. The resulting mixture was heated at 185 °C for 5 min. Upon completion, the mixture was cooled to 0 °C and was quenched with ice/water. The acylated product precipitated and was collected by filtration. Purification by silica gel flash column chromatography (cyclohexane:EtOAc 9:1 to 1:1) afforded the title compound.

#### General Procedure D. Sulphonamide coupling

To a solution of 2-oxoindoline-5-sulfonyl chloride (1.0 equiv) in THF the corresponding amine (1.1 equiv) and DIPEA (3.0 equiv) at room temperature, and the resulting mixture was stirred for 18 h at room temperature. Upon completion, the mixture was diluted with EtOAc and saturated solution of NaHCO_3_ and layers were separated. The organics were further extracted from the aqueous with EtOAc and the combined organics were washed with brine, dried over Na_2_SO_4_, filtered and the solvent was removed by rotary evaporation. Purification by silica gel flash column chromatography (hexane:EtOAc or CH_2_Cl_2_:MeOH mixtures) afforded the desired compound.

#### General Procedure E. Mitsunobu reaction

A dry three-neck flask was charged with 5-hydroxyindolin-2-one (1.0 equiv), PPh_3_ (1.4 equiv) and the appropriate alcohol (1.4 equiv) in anhydrous THF (0.25M). The solution was cooled to 0 °C and diethyl azadicarboxylate (1.4 equiv) in PhMe was added dropwise. The resulting mixture was warmed to room temperature and stirred for 18 h. Upon completion, the solvent reduced *in vacuo*, before purification by silica gel flash column chromatography (hexane:EtOAc 8:2) to afford the title compound.

#### General Procedure F. Amide HATU coupling

To a solution of amine (1.0 equiv) in THF, HATU (1.3 equiv), the corresponding carboxylic acid (1.1 equiv) and DIPEA (3.0 equiv) at room temperature, and the resulting mixture was stirred for 18 h at room temperature. Upon completion, the mixture was diluted with EtOAc and saturated solution of NaHCO_3_ and layers were separated. The organics were further extracted from the aqueous with EtOAc and the combined organics were washed with brine, dried over Na_2_SO_4_, filtered and the solvent reduced *in vacuo*. Purification by silica gel flash column chromatography (hexane:EtOAc or CH_2_Cl_2_:MeOH mixtures) afforded the desired compound.

##### (3Z)-3-[(1H-imidazol-5-yl)methylidene]-5-methoxy-2,3-dihydro-1H-indol-2-one (1)

Compound **1** was prepared by procedure A to afford a purple solid (177 mg, 0.734 mmol, 80%). MP 210-212 °C; ^1^H NMR (400 MHz, DMSO-*d*_6_) δ 13.81 (s, 1H), 10.82 (s, 1H), 8.02 (s, 1H), 7.88 (s, 1H), 7.61 (s, 1H), 7.36 – 7.35 (m, 1H), 6.80 – 6.76 (m, 2H), 3.76 (s, 3H). HRMS m/z [M+H]^+^ calcd for C_13_H_12_N_3_O_2_: 242.0930, found 242.0918, LC t_R_ = 2.94 min, > 98% Purity. Consistent with previous report.^85^

##### (3Z)-5-methoxy-3-(1H-pyrrol-2-ylmethylidene)-2,3-dihydro-1H-indol-2-one (2)

Compound **2** was prepared by procedure A to afford a dark orange solid (168 mg, 0.699 mmol, 76%). MP 220-222 °C; ^1^H NMR (400 MHz, DMSO-*d*_6_) δ 13.16 (s, 1H), 10.84 (s, 1H), 7.56 (s, 1H), 7.52 (d, *J* = 8.4 Hz, 1H), 7.29 (q, *J* = 2.3 Hz, 1H), 6.75 (dt, *J* = 3.6, 1.7 Hz, 1H), 6.58 (dd, *J* = 8.4, 2.3 Hz, 1H), 6.45 (d, *J* = 2.3 Hz, 1H), 6.34 – 6.29 (m, 1H), 3.76 (s, 3H). ^13^C NMR (100 MHz, DMSO-*d*_6_) δ 169.8, 159.3, 140.3, 129.6, 124.7, 124.2, 119.6, 119.1, 117.9, 117.1, 111.1, 107.0, 96.1, 55.3. HRMS m/z [M+H]^+^ calcd for C_14_H_13_N_2_O_2_: 241.0977, found 241.0961, LC t_R_ = 4.38 min, > 98% Purity.

##### (3Z)-3-[(3,5-dimethyl-1H-pyrrol-2-yl)methylidene]-5-methoxy-2,3-dihydro-1H-indol-2-one (3)

Compound **3** was prepared by procedure A to afford a red solid (200 mg, 0.745 mmol, 81%). MP 231-233 °C; ^1^H NMR (400 MHz, DMSO-*d*_6_) δ 13.45 (s, 1H), 10.57 (s, 1H), 7.58 (s, 1H), 7.40 (d, *J* = 2.4 Hz, 1H), 6.76 (d, *J* = 8.4 Hz, 1H), 6.67 (dd, *J* = 8.4, 2.4 Hz, 1H), 5.99 (d, *J* = 2.5 Hz, 1H), 3.77 (s, 3H), 2.31 (s, 6H). ^13^C NMR (100 MHz, DMSO-*d*_6_) δ 169.5, 154.8, 135.6, 132.0, 131.6, 126.9, 126.6, 123.7, 113.3, 112.5, 111.9, 109.6, 104.1, 55.6, 13.5, 11.4. HRMS m/z [M+H]^+^ calcd for C_16_H_17_N_2_O_2_: 269.1290, found 269.1282, LC t_R_ = 4.79 min, > 98% Purity.

##### (3Z)-5-methoxy-3-(1H-pyrazol-5-ylmethylidene)-2,3-dihydro-1H-indol-2-one (4)

Compound **4** was prepared by procedure A to afford a red solid (160 mg, 0.663 mmol, 72%). MP 192-194 °C; ^1^H NMR (400 MHz, DMSO-*d*_6_) δ 10.93 (s, 1H), 7.87 (s, 2H), 7.73 (s, 1H), 7.40 (d, *J* = 2.2 Hz, 1H), 6.86 – 6.77 (m, 3H), 3.77 (s, 3H). ^13^C NMR (100 MHz, DMSO-*d*_6_) δ 170.0, 154.8, 136.6, 134.5, 130.3, 126.0, 123.2, 114.9, 113.6, 111.0, 109.8, 106.3, 55.9. HRMS m/z [M+H]^+^ calcd for C_13_H_12_N_3_O_2_: 242.0930, found 242.0920, LC t_R_ = 3.85 min, > 98% Purity.

##### (3Z)-3-(1H-imidazol-2-ylmethylidene)-5-methoxy-2,3-dihydro-1H-indol-2-one (5)

Compound **5** was prepared by procedure A to afford a red solid (164 mg, 0.680 mmol, 74%). MP >300 °C; ^1^H NMR (400 MHz, DMSO-*d*_6_) δ 10.97 (s, 1H), 7.86 (s, 1H), 7.55 (t, *J* = 1.5 Hz, 2H), 7.34 (d, *J* = 1.2 Hz, 1H), 6.82 (d, *J* = 1.4 Hz, 2H), 3.77 (s, 3H). ^13^C NMR (100 MHz, DMSO-*d*_6_) δ 168.9, 155.2, 143.7, 133.7, 132.6, 125.0, 124.7, 124.1, 120.6, 115.1, 110.5, 106.0, 55.6. HRMS m/z [M+H]^+^ calcd for C_13_H_12_N_3_O_2_: 242.0930, found 242.0917, LC t_R_ = 2.80 min, > 98% Purity.

##### (3Z)-5-methoxy-3-(pyridin-2-ylmethylidene)-2,3-dihydro-1H-indol-2-one (6)

Compound **6** was prepared by procedure A to afford a dark red solid (176 mg, 0.698 mmol, 76%). MP 176-178 °C; ^1^H NMR (400 MHz, DMSO-*d*_6_) δ 10.42 (s, 1H), 8.96 – 8.86 (m, 1H), 8.82 (d, *J* = 2.7 Hz, 1H), 7.96 (td, *J* = 7.6, 1.8 Hz, 1H), 7.88 (dt, *J* = 7.8, 1.1 Hz, 1H), 7.56 (s, 1H), 7.47 (ddd, *J* = 7.6, 4.7, 1.2 Hz, 1H), 6.88 (dd, *J* = 8.4, 2.7 Hz, 1H), 6.77 (d, *J* = 8.4 Hz, 1H), 3.77 (s, 3H). ^13^C NMR (100 MHz, DMSO-*d*_6_) δ 169.3, 154.2, 153.2, 149.6, 137.3, 137.3, 133.7, 129.9, 128.6, 124.2, 122.3, 116.3, 114.0, 109.8, 55.3. HRMS m/z [M+H]^+^ calcd for C_15_H_13_N_2_O_2_: 253.0977, found 253.0966, LC t_R_ = 3.88 min, > 98% Purity.

##### (3Z)-5-methoxy-3-(pyridin-3-ylmethylidene)-2,3-dihydro-1H-indol-2-one (7)

Compound **7** was prepared by procedure A to afford a red solid (104 mg, 0.412 mmol, 45%, ratio of E:Z - 50:50). MP 206-208 °C; (*Z*)-5-Methoxy-3-(pyridin-3-ylmethylene)indolin-2-one. ^1^H NMR (500 MHz, DMSO-*d*_6_) δ 10.5 (d, *J* = 6.0 Hz, 1H), 9.19 (dd, *J* = 1.8, 1.1 Hz, 1H), 8.91 (dddd, *J* = 8.1, 2.3, 1.6, 0.6 Hz, 1H), 8.58 (dd, *J* = 4.8, 1.7 Hz, 1H), 7.84 (s, 1H), 7.49 (ddd, *J* = 8.0, 4.9, 0.9 Hz, 1H), 7.41 (d, *J* = 2.5 Hz, 1H), 6.82 (dd, *J* = 8.4, 2.5 Hz, 1H), 6.74 (dd, *J* = 8.4, 0.5 Hz, 1H), 3.77 (s, 3H). ^13^C NMR (125 MHz, DMSO-*d*_6_) δ 167.1, 154.7, 152.4, 150.3, 137.9, 134.9, 133.0, 129.9, 129.3, 125.3, 123.2, 115.4, 110.1, 106.3, 55.6. (*E*)-5-Methoxy-3-(pyridin-3-ylmethylene)indolin-2-one. ^1^H NMR (500 MHz, DMSO-*d*_6_) δ 10.5 (d, *J* = 6.0 Hz, 2H), 8.862 – 8.857 (m, 1H), 8.65 (ddd, *J* = 4.9, 1.7, 0.6 Hz, 1H), 8.12 (dddd, *J* = 7.9, 2.4, 1.7, 0.8 Hz, 1H), 7.62 (s, 1H), 7.56 (ddd, *J* = 7.9, 4.7, 0.5 Hz, 1H), 6.89 – 6.89 (m, 1H), 6.86 (dd, *J* = 8.4, 2.5 Hz, 1H), 6.80 (dd, *J* = 8.4, 0.5 Hz, 1H), 3.60 (s, 3H). ^13^C NMR (125 MHz, DMSO-*d*_6_) δ 168.1, 154.0, 150.2, 149.7, 136.9, 136.3, 132.3, 130.5, 129.9, 123.6, 121.4, 115.6, 110.7, 108.9, 55.3. HRMS m/z [M+H]^+^ calcd for C_15_H_13_N_2_O_2_: 253.0977, found 253.0966, LC t_R_ = 3.10 min, > 98% Purity.

##### (3Z)-3-(1H-pyrazol-5-ylmethylidene)-2,3-dihydro-1H-indol-2-one (8)

Compound **8** was prepared by procedure A to afford a bright yellow solid (156 mg, 0.739 mmol, 49%, ratio of E:Z - 33:67). MP 249-251 °C; (*Z*)-3-((1*H-*Pyrazol-5-yl)methylene)indolin-2-one. ^1^H NMR (600 MHz, DMSO-*d*_6_) δ 10.39 (bs, 1H), 9.31 (d, *J* = 7.6 Hz, 1H), 8.00 (s, 1H), 7.90 (s, 1H), 7.47 (s, 1H), 7.16 (td, *J* = 7.6, 1.4 Hz, 1H), 6.96 (td, *J* = 7.6, 1.1 Hz, 1H), 6.82 (d, *J* = 8.5 Hz, 1H). ^13^C NMR (125 MHz, DMSO-*d*_6_) δ 169.9, 141.5, 138.1, 137.0, 128.2, 126.9, 126.4, 126.1, 122.6, 121.28, 120.6, 108.7. (*E*)-3-((1*H-*Pyrazol-5-yl)methylene)indolin-2-one. ^1^H NMR (600 MHz, DMSO-*d*_6_) δ 7.99 (s, 1H), 7.83 (s, 1H), 7.66 (d, *J* = 6.9 Hz, 1H), 7.21–7.15 (m, 1H), 7.01 (td, *J* = 7.6, 1.0 Hz, 1H), 6.89 (d, *J* = 7.7 Hz, 1H). ^13^C NMR (125 MHz, DMSO-*d*_6_) δ 168.6, 139.8, 139.0, 138.1, 127.9, 124.3, 123.9, 121.3, 120.6, 120.2, 119.2, 109.5. HRMS m/z [M+H]^+^ calcd for C_12_H_10_N_3_O: 212.0824, found 212.0814, LC t_R_ = 2.82 min, > 98% Purity.

##### (3Z)-3-(1*H*-pyrrol-2-ylmethylidene)-2,3-dihydro-1H-indol-2-one (9)

Compound **9** was prepared by procedure A to afford a dark orange solid (159 mg, 0.756 mmol, 67%). MP 218-220 °C; ^1^H NMR (400 MHz, DMSO-*d*_6_) δ 13.34 (s, 1H), 10.89 (s, 1H), 7.74 (s, 1H), 7.63 (dt, *J* = 7.4, 0.9 Hz, 1H), 7.35 (td, *J* = 2.6, 1.4 Hz, 1H), 7.15 (td, *J* = 7.6, 1.2 Hz, 1H), 6.99 (td, *J* = 7.6, 1.1 Hz, 1H), 6.88 (dt, *J* = 7.8, 0.9 Hz, 1H), 6.83 (dt, *J* = 3.6, 1.7 Hz, 1H), 6.35 (dt, *J* = 3.7, 2.3 Hz, 1H). ^13^C NMR (100 MHz, DMSO-*d*_6_) δ 169.2, 139.0, 129.6, 126.9, 126.3, 125.6, 125.2, 121.2, 120.3, 118.5, 116.8, 111.4, 109.5. HRMS m/z [M+H]^+^ calcd for C_13_H_11_N_2_O: 211.0871, found 211.0865, LC t_R_ = 4.47 min, > 98% Purity.

##### (3Z)-3-(1H-pyrazol-5-ylmethylidene)-2,3-dihydro-1H-indol-2-one (11)

Compound **11** was prepared by procedure A to afford an orange solid (90 mg, 0.426 mmol, 38%). MP 190-192 °C; ^1^H NMR (600 MHz, DMSO-*d*_6_) δ 13.56 (s, 1H), 10.52 (s, 1H), 7.94 (s, 1H), 7.72 (d, *J* = 7.5 Hz, 1H), 7.48 (s, 1H), 7.27 – 7.24 (m, 1H), 7.00 (td, *J* = 7.6, 1.1 Hz, 1H), 6.87 (d, *J* = 7.7 Hz, 1H), 6.85 – 6.80 (m, 1H). ^13^C NMR (150 MHz, DMSO-*d*_6_) δ 169.5, 146.3, 142.5, 140.4, 129.7, 126.2, 125.8, 125.0, 121.9, 121.0, 110.6, 109.4. HRMS m/z [M+H]^+^ calcd for C_12_H_10_N_3_O: 212.0824, found 212.0812, LC t_R_ = 3.85 min, > 98% Purity.

##### (3Z)-3-(1H-imidazol-2-ylmethylidene)-2,3-dihydro-1H-indol-2-one (12)

Compound **12** was prepared by procedure A to afford a bright yellow solid (195 mg, 0.923 mmol, 82% ratio of E:Z - 40:60 - not able to clearly distinguish *E*/Z isomers). MP >300 °C; Major diastereoisomer. 3-((1*H-*Imidazol-2-yl)methylene)indolin-2-one. ^1^H NMR (600 MHz, DMSO-*d*_6_) δ 10.52 (s, 1H), 9.40 (dd, *J* = 7.7, 1.3 Hz, 1H), 7.45 (s, 2H), 7.39 (s, 1H), 7.24 – 7.22 (m, 1H), 7.00 (td, *J* = 7.6, 1.1 Hz, 1H), 6.85 (d, *J* = 7.7 Hz, 1H). ^13^C NMR (150 MHz, DMSO-*d*_6_) δ 169.6, 143.4, 142.4, 129.6, 128.9, 127.1, 124.3, 121.8, 121.0, 120.9, 120.2, 109.1. Minor diastereoisomer. 3-((1*H-*Imidazol-2-yl)methylene)indolin-2-one. ^1^H NMR (600 MHz, DMSO-d_6_) δ 14.10 (s, 1H), 7.83 (d, *J* = 7.4 Hz, 1H), 7.80 (s, 1H), 7.55 (bs, 1H), 7.34 (s, 1H), 7.26 – 7.23 (m, 1H), 7.03 (td, *J* = 7.6, 1.0 Hz, 1H), 6.92 (d, *J* = 7.8 Hz, 1H). 13C NMR (150 MHz, DMSO-*d*_6_) δ 168.9, 143.7, 140.1, 132.5, 127.1, 124.4, 124.0, 123.7, 122.0, 121.0, 120.7, 110.0. HRMS m/z [M+H]^+^ calcd for C_12_H_10_N_3_O: 212.0824, found 212.0813, LC t_R_ = 2.60 min, > 98% Purity.

##### (3Z)-3-(pyridin-2-ylmethylidene)-2,3-dihydro-1H-indol-2-one (13)

Compound **13** was prepared by procedure A to afford a dark orange solid (160 mg, 0.720 mmol, 64%). MP 191-193 °C; ^1^H NMR (400 MHz, DMSO-*d*_6_) δ 10.62 (s, 1H), 9.03 – 8.97 (m, 1H), 8.88 (ddd, *J* = 4.8, 1.8, 0.8 Hz, 1H), 7.94 (td, *J* = 7.7, 1.8 Hz, 1H), 7.86 (dt, *J* = 7.9, 1.1 Hz, 1H), 7.57 (s, 1H), 7.46 (ddd, *J* = 7.5, 4.7, 1.2 Hz, 1H), 7.28 (td, *J* = 7.7, 1.3 Hz, 1H), 6.98 (td, *J* = 7.7, 1.1 Hz, 1H), 6.87 (dt, *J* = 7.7, 0.9 Hz, 1H). ^13^C NMR (100 MHz, DMSO-*d*_6_) δ 169.3, 153.2, 149.7, 143.6, 137.2, 133.7, 130.8, 129.3, 128.4, 127.9, 124.1, 121.5, 121.2, 109.6. HRMS m/z [M+H]^+^ calcd for C_14_H_11_N_2_O: 223.0871, found 223.0860, LC t_R_ = 3.85 min, > 98% Purity.

##### (3Z)-3-(pyridin-3-ylmethylidene)-2,3-dihydro-1H-indol-2-one (14)

Compound **14** was prepared by procedure A to afford a light yellow solid (45 mg, 0.203 mmol, 18%). MP 155-157 °C; ^1^H NMR (400 MHz, DMSO-*d*_6_) δ 10.66 (s, 1H), 8.87 (dt, *J* = 1.9, 0.9 Hz, 1H), 8.65 (dd, *J* = 4.9, 1.6 Hz, 1H), 8.12 (dddd, *J* = 7.9, 2.4, 1.7, 0.8 Hz, 1H), 7.62 (s, 1H), 7.55 (ddd, *J* = 7.9, 4.8, 0.9 Hz, 1H), 7.37 (dd, *J* = 7.8, 1.1 Hz, 1H), 7.27 – 7.23 (m, 1H), 6.91 – 6.85 (m, 2H). ^13^C NMR (100 MHz, DMSO-*d*_6_) δ 168.1, 150.1, 149.8, 143.2, 136.4, 132.1, 130.6, 130.6, 129.4, 123.7, 122.3, 121.3, 120.6, 110.3. HRMS m/z [M+H]^+^ calcd for C_14_H_11_N_2_O: 223.0871, found 223.0861, LC t_R_ = 2.60 min, > 98% Purity.

##### (3Z)-6-methoxy-3-(1H-pyrrol-2-ylmethylidene)-2,3-dihydro-1H-indol-2-one (15)

Compound **15** was prepared by procedure A to afford an orange solid (137 mg, 0.570 mmol, 62%). MP 225-226 °C; ^1^H NMR (400 MHz, DMSO-*d*_6_) δ 13.16 (s, 1H), 10.84 (s, 1H), 7.56 (s, 1H), 7.52 (d, *J* = 8.4 Hz, 1H), 7.29 (td, *J* = 2.6, 1.4 Hz, 1H), 6.75 (dt, *J* = 3.6, 1.7 Hz, 1H), 6.58 (dd, *J* = 8.4, 2.3 Hz, 1H), 6.45 (d, *J* = 2.3 Hz, 1H), 6.32 (dt, *J* = 3.7, 2.4 Hz, 1H), 3.76 (s, 3H). ^13^C NMR (100 MHz, DMSO-*d*_6_) δ 169.8, 159.3, 140.3, 129.6, 124.7, 124.2, 119.6, 119.1, 117.9, 117.1, 111.1, 107.0, 96.1, 55.3. HRMS m/z [M+H]^+^ calcd for C_14_H_13_N_2_O_2_: 241.0977, found 241.0962, LC t_R_ = 4.41 min, > 98% Purity.

##### (3Z)-3-[(3,5-dimethyl-1H-pyrrol-2-yl)methylidene]-6-methoxy-2,3-dihydro-1H-indol-2-one (16)

Compound **16** was prepared by procedure A to afford an orange solid (200 mg, 0.745 mmol, 81%). MP 231-233 °C; ^1^H NMR (400 MHz, DMSO-*d*_6_) δ 13.14 (s, 1H), 10.72 (s, 1H), 7.60 (d, *J* = 8.4 Hz, 1H), 7.40 (s, 1H), 6.56 (dd, *J* = 8.4, 2.3 Hz, 1H), 6.45 (d, *J* = 2.3 Hz, 1H), 5.96 (d, *J* = 2.5 Hz, 1H), 3.75 (s, 3H), 2.30 (s, 3H), 2.27 (s, 3H). ^13^C NMR (100 MHz, DMSO-*d*_6_) δ 169.9, 158.5, 139.4, 134.4, 130.1, 126.5, 121.6, 119.1, 118.6, 113.1, 112.0, 106.7, 95.8, 55.2, 13.5, 11.3. HRMS m/z [M+H]^+^ calcd for C_16_H_17_N_2_O_2_: 269.1290, found 269.1272, LC t_R_ = 4.67 min, > 98% Purity.

##### (3Z)-6-methoxy-3-(1H-pyrazol-5-ylmethylidene)-2,3-dihydro-1H-indol-2-one (17)

Compound **17** was prepared by procedure A to afford an orange solid (189 mg, 0.783 mmol, 85%). MP 209-211 °C; ^1^H NMR (400 MHz, DMSO-*d*_6_) δ 11.06 (s, 1H), 7.80 (s, 1H), 7.64 (dd, *J* = 19.1, 8.0 Hz, 3H), 6.85 (s, 1H), 6.62 (d, *J* = 8.4 Hz, 1H), 6.47 (s, 1H), 3.78 (s, 3H). ^13^C NMR (100 MHz, DMSO-*d*_6_) δ 169.3, 160.8, 141.8, 140.2, 138.0, 125.0, 121.3, 118.6, 116.5, 110.7, 107.6, 96.5, 55.4. HRMS m/z [M+H]^+^ calcd for C_13_H_12_N_3_O_2_: 242.0930, found 242.0917, LC t_R_ = 3.54 min, > 98% Purity.

##### (3Z)-3-(1H-imidazol-2-ylmethylidene)-6-methoxy-2,3-dihydro-1H-indol-2-one (18)

Compound **18** was prepared by procedure A to afford an orange solid (126 mg, 0.522 mmol, 57%). MP >300 °C; ^1^H NMR (600 MHz, DMSO-*d*_6_) δ 13.94 (s, 1H), 11.33 (s, 1H), 7.73 (d, *J* = 8.4 Hz, 1H), 7.61 (s, 1H), 7.50 (s, 1H), 7.29 (s, 1H), 6.60 (dd, *J* = 8.4, 2.3 Hz, 1H), 6.48 (d, *J* = 2.3 Hz, 1H), 3.78 (s, 3H). ^13^C NMR (150 MHz, DMSO-*d*_6_) δ 169.6, 160.6, 143.9, 141.7, 132.0, 123.8, 121.9, 121.5, 120.1, 116.7, 107.5, 96.4, 55.4. HRMS m/z [M+H]^+^ calcd for C_13_H_12_N_3_O_2_: 242.0930, found 242.0916, LC t_R_ = 2.82 min, > 98% Purity.

##### (3Z)-6-methoxy-3-(pyridin-2-ylmethylidene)-2,3-dihydro-1H-indol-2-one (19)

Compound **19** was prepared by procedure A to afford a mustard solid (151 mg, 0.599 mmol, 65%). MP 182-184 °C; δ ^1^H NMR (400 MHz, DMSO-*d*_6_) δ 10.57 (s, 1H), 8.98 (d, *J* = 8.6 Hz, 1H), 8.89 – 8.78 (m, 1H), 7.92 (td, *J* = 7.7, 1.9 Hz, 1H), 7.81 (dt, *J* = 8.0, 1.1 Hz, 1H), 7.42 (ddd, *J* = 7.6, 4.7, 1.2 Hz, 1H), 7.39 (s, 1H), 6.55 (dd, *J* = 8.7, 2.5 Hz, 1H), 6.42 (d, *J* = 2.4 Hz, 1H), 3.79 (s, 3H). ^13^C NMR (100 MHz, DMSO-*d*_6_) δ 170.0, 161.7, 153.6, 149.5, 145.5, 137.1, 130.4, 129.6, 128.9, 128.0, 123.7, 114. 6, 106.5, 96.0, 55.3. HRMS m/z [M+H]^+^ calcd for C_15_H_13_N_2_O_2_: 253.0977, found 253.0970, LC t_R_ = 3.85 min, > 98% Purity.

##### (Z)-6-methoxy-3-(pyridin-3-ylmethylene)indolin-2-one (20)

Compound **20** was prepared by procedure A to afford a mustard solid (93 mg, 0.369 mmol, 40%, ratio of E:Z - 28:72). MP 177-179 °C; (*Z*)-6-Methoxy-3-(pyridin-3-ylmethylene)indolin-2-one. ^1^H NMR (600 MHz, DMSO-*d*_6_) δ 10.73 (bs, 1H), 8.84 (d, *J* = 2.3 Hz, 1H), 8.62 (dd, *J* = 4.9, 1.6 Hz, 1H), 8.08 (dt, *J* = 7.9, 2.1 Hz, 1H), 7.53 (dd, *J* = 7.9, 4.8 Hz, 1H), 7.43 (s, 1H), 7.32 (d, *J* = 8.5 Hz, 1H), 6.45 – 6.42 (m, 2H), 3.75 (s, 3H). 13C NMR (150 MHz, DMSO-*d*_6_) δ 168.9, 161.5, 149.77, 149.75, 145.1, 136.2, 130.9, 129.0, 128.6, 123.7, 123.1, 113.4, 106.8, 96.7, 55.4. (*E*)-6-Methoxy-3-(pyridin-3-ylmethylene)indolin-2-one. ^1^H NMR (600 MHz, DMSO-*d*_6_) δ 9.13 (d, *J* = 2.2 Hz, 1H), 8.82 (dt, *J* = 8.1, 2.0 Hz, 1H), 8.54 (dd, *J* = 4.8, 1.7 Hz, 1H), 7.62 (s, 1H), 7.62 (d, *J* = 8.1 Hz, 1H), 7.46 (dd, *J* = 8.1, 4.8 Hz, 1H), 6.57 (dd, *J* = 8.4, 2.4 Hz, 1H), 6.40 (d, *J* = 2.3 Hz, 1H), 3.78 (s, 3H). ^13^C NMR (150 MHz, DMSO-*d*_6_) δ 167.7, 161.0, 152.1, 149.7, 142.7, 137.5, 130.2, 129.9, 128.6, 123.5, 121.5, 117.1, 106.9, 96.0, 55.4. HRMS m/z [M+H]^+^ calcd for C_15_H_13_N_2_O_2_: 253.0977, found 253.0965, LC t_R_ = 2.70 min, > 98% Purity.

##### (Z)-3-(1H-imidazol-5-ylmethylidene)-5,6-dimethoxy-2,3-dihydro-1H-indol-2-one (21)

Compound **21** was prepared by procedure A to afford a bright red solid (160 mg, 0.590 mmol, 76%, ratio of E:Z - 50:50). MP 222-224 °C; (*Z*)-3-((1H*-*Imidazol-5-yl)methylene)-5,6-dimethoxyindolin-2-one. ^1^H NMR (600 MHz, DMSO-*d*_6_) δ 13.71 (bs, 1H), 7.94 (s, 1H), 7.68 (s, 1H), 7.56 (s, 1H), 7.38 (s, 1H), 6.53 (s, 1H), 3.77 (s, 3H), 3.77 (s, 3H). (*E*)-3-((1*H-*Imidazol-5-yl)methylene)-5,6-dimethoxyindolin-2-one. ^1^H NMR (600 MHz, DMSO-*d*_6_) δ 10.14 (s, 1H), 9.20 (s, 1H), 7.97 (s, 1H), 7.81 (d, *J* = 1.1 Hz, 1H), 7.30 (s, 1H), 6.47 (s, 1H), 3.78 (s, 3H), 3.76 (s, 3H). ^13^C NMR (150 MHz, DMSO-*d*_6_) δ 170.6, 150.0, 143.1, 137.9, 136.7, 125.1, 124.3, 122.2, 114.1, 113.1, 104.8, 94.6, 56.5, 55.6. HRMS m/z [M+H]^+^ calcd for C_14_H_14_N_3_O_3_: 272.1035, found 272.1023, LC t_R_ = 2.76 min, > 98% Purity.

##### (3Z)-5,6-dimethoxy-3-(1H-pyrrol-2-ylmethylidene)-2,3-dihydro-1H-indol-2-one (22)

Compound **22** was prepared by procedure A to afford a bright red solid (93 mg, 0.369 mmol, 40%). MP 167-169 °C; ^1^H NMR (400 MHz, DMSO-*d*_6_) δ 13.27 (s, 1H), 10.65 (s, 1H), 7.61 (s, 1H), 7.35 (s, 1H), 7.29 (q, *J* = 2.2 Hz, 1H), 6.73 (dt, *J* = 3.6, 1.7 Hz, 1H), 6.52 (s, 1H), 6.32 (dt, *J* = 4.0, 2.4 Hz, 1H), 3.77 (d, *J* = 2.2 Hz, 6H). ^13^C NMR (100 MHz, DMSO-*d*_6_) δ 169.6, 149.1, 144.4, 133.2, 129.7, 124.6, 124.4, 118.9, 117.9, 116.3, 111.1, 104.3, 95.4, 56.3, 55.8. HRMS m/z [M+H]^+^ calcd for C_15_H_15_N_2_O_3_: 271.1083, found 271.1072, LC t_R_ = 4.29 min, > 98% Purity.

##### (3Z)-3-[(3,5-dimethyl-1H-pyrrol-2-yl)methylidene]-5,6-dimethoxy-2,3-dihydro-1H-indol-2-one (23)

Compound **23** was prepared by procedure A to afford a dark red solid (167 mg, 0.560 mmol, 72%). MP 282-285 °C; ^1^H NMR (400 MHz, DMSO-*d*_6_) δ 13.28 (s, 1H), 10.53 (s, 1H), 7.45 (s, 1H), 7.42 (s, 1H), 6.51 (s, 1H), 5.96 (d, *J* = 2.5 Hz, 1H), 3.78 (s, 3H), 3.75 (s, 3H), 2.30 (s, 6H). 13C NMR (100 MHz, DMSO-*d*_6_) δ ^13^C NMR (101 MHz, DMSO-*d*_6_) δ 169.7, 148.4, 144.3, 134.3, 132.3, 130.1, 126.6, 121.9, 117.2, 114.0, 112.0, 104.3, 95.3, 56.6, 55.8, 13.5, 11.4. HRMS m/z [M+H]^+^ calcd for C_22_H_13_N_4_O: 349.1089, found 349.1088, LC t_R_ = 4.61 min, > 98% Purity.

##### (3Z)-5,6-dimethoxy-3-(1H-pyrazol-5-ylmethylidene)-2,3-dihydro-1H-indol-2-one (24)

Compound **24** was prepared by procedure A to afford a dark red solid (160 mg, 0.590 mmol, 76%). MP 230-232 °C; ^1^H NMR (400 MHz, DMSO-*d*_6_) δ 13.49 (s, 1H), 10.26 (s, 1H), 8.85 (s, 1H), 7.89 (d, *J* = 2.3 Hz, 1H), 7.29 (s, 1H), 6.75 (d, *J* = 2.3 Hz, 1H), 6.51 (s, 1H), 3.80 (s, 3H), 3.77 (s, 3H).^13^C NMR (100 MHz, DMSO-*d*_6_) δ 170.2, 150.9, 146.6, 143.2, 137.8, 129.7, 125.7, 122.6, 113.2, 112.6, 109.9, 94.9, 56.5, 55.6. HRMS m/z [M+H]^+^ calcd for C_14_H_14_N_3_O_3_: 272.1035, found 272.1028, LC t_R_ = 3.21 min, > 98% Purity.

##### (Z)-3-(1H-imidazol-2-ylmethylidene)-5,6-dimethoxy-2,3-dihydro-1H-indol-2-one (25)

Compound **25** was prepared by procedure A to afford a dark red solid (95 mg, 0.350 mmol, 45%, ratio of E:Z - 44:56). MP >300 °C; (*E*)-3-((1H-Imidazol-2-yl)methylene)-5,6-dimethoxyindolin-2-one. ^1^H NMR (500 MHz, DMSO-*d*_6_) δ 10.28 (bs, 1H), 9.26 (s, 1H), 7.41 (s, 2H), 7.21 (s, 1H), 6.50 (s, 1H), 3.80 (s, 3H), 3.77 (s, 3H). ^13^C NMR (125 MHz, DMSO-*d*_6_) δ 170.2, 151.1, 143.5, 143.2, 137.8, 132.1, 125.6, 117.5, 113.28, 113.26, 94.8, 56.5, 55.6. (*Z*)-3-((1H-Imidazol-2-yl)methylene)-5,6-dimethoxyindolin-2-one. ^1^H NMR (500 MHz, DMSO-*d*_6_) δ 14.01 (bs, 1H), 7.69 (s, 1H), 7.56 (s, 1H), 7.50 (bs, 1H), 7.30 (bs, 1H), 6.54 (s, 1H), 3.78 (s, 3H), 3.77 (s, 3H). ^13^C NMR (125 MHz, DMSO-*d*_6_) δ 169.4, 150.6, 144.7, 144.0, 134.5, 124.5, 122.2, 120.0, 115.1, 105.5, 95.5, 56.3, 55.8. HRMS m/z [M+H]^+^ calcd for C_14_H_14_N_3_O_3_: 272.1035, found 272.1022, LC t_R_ = 2.74 min, > 98% Purity.

##### (Z)-5-acetyl-3-(1H-pyrazol-5-ylmethylidene)-2,3-dihydro-1H-indol-2-one (26)

Compound **26** was prepared by procedure A to afford a bright orange solid (154 mg, 0.608 mmol, 71%, *E*:Z ratio 32:68). MP 262-265 °C; (*Z*)-3-((1*H-*Pyrazol-5-yl)methylene)-5-acetylindolin-2-one. ^1^H NMR (500 MHz, DMSO-*d*_6_) δ 10.80 (s, 1H), 10.12 (d, *J* = 1.8 Hz, 1H), 8.02 (s, 1H), 7.94 (d, *J* = 1.1 Hz, 1H), 7.82 (dd, *J* = 8.1, 1.9 Hz, 1H), 7.56 (s, 1H), 6.92 (d, *J* = 8.2 Hz, 1H), 2.57 (s, 3H). ^13^C NMR (150 MHz, DMSO-*d*_6_) δ 196.8, 170.5, 145.4, 139.7, 137.0, 130.4, 128.9, 127.5, 122.7, 119.7, 119.4, 108.6, 26.5. (*E*)-3-((1*H-*Pyrazol-5-yl)methylene)-5-acetylindolin-2-one. ^1^H NMR (600 MHz, DMSO-*d*_6_) δ 10.80 (s, 1H), 8.33 (d, *J* = 1.7 Hz, 1H), 8.06 (s, 1H), 8.01 (s, 1H), 7.85 (dd, *J* = 8.2, 1.7 Hz, 1H), 6.99 (d, *J* = 8.2 Hz, 1H), 2.58 (s, 3H). HRMS m/z [M+H]^+^ calcd for C_14_H_12_N_3_O_2_: 254.0930, found 254.0918, LC t_R_ = 2.74 min, > 98% Purity.

##### (3Z)-5-acetyl-3-(1H-pyrrol-2-ylmethylidene)-2,3-dihydro-1H-indol-2-one (27)

Compound **27** was prepared by procedure A to afford a dark orange solid (117 mg, 0.467 mmol, 54%). MP 218-220 °C; ^1^H NMR (400 MHz, DMSO-*d*_6_) δ 10.58 (s, 1H), 9.08 – 8.92 (m, 1H), 8.85 (ddd, *J* = 4.8, 1.9, 0.8 Hz, 1H), 7.92 (td, *J* = 7.7, 1.8 Hz, 1H), 7.84 (dt, *J* = 7.9, 1.1 Hz, 1H), 7.43 (ddd, *J* = 7.6, 4.7, 1.2 Hz, 1H), 7.25 (td, *J* = 7.6, 1.3 Hz, 1H), 6.95 (td, *J* = 7.7, 1.1 Hz, 1H), 6.84 (dt, *J* = 7.7, 0.8 Hz, 1H), 3.31 (s, 3H). ^13^C NMR (100 MHz, DMSO-*d*_6_) δ 169.3, 153.2, 149.7, 143.6, 137.3, 133.7, 130.8, 129.3, 128.4, 127.9, 124.1, 121.5, 121.2, 109.6, 39.3. HRMS m/z [M+H]^+^ calcd for C_15_H_13_N_2_O_2_: 253.0977, found 253.0963, LC t_R_ = 3.99 min, > 98% Purity.

##### (3Z)-5-acetyl-3-[(3,5-dimethyl-1H-pyrrol-2-yl)methylidene]-2,3-dihydro-1H-indol-2-one (28)

Compound **28** was prepared by procedure A to afford a dark orange solid (149 mg, 0.532 mmol, 62%). MP >300 °C; δ ^1^H NMR (400 MHz, DMSO-*d*_6_) δ 13.33 (s, 1H), 11.14 (s, 1H), 8.35 (d, *J* = 1.6 Hz, 1H), 7.82 – 7.68 (m, 2H), 6.96 (d, *J* = 8.2 Hz, 1H), 6.05 (d, *J* = 2.5 Hz, 1H), 2.59 (s, 3H), 2.35 (s, 3H), 2.33 (s, 3H). ^13^C NMR (100 MHz, DMSO-*d*_6_) δ 197.0, 169.8, 141.8, 136.8, 133.1, 130.5, 126.9, 126.6, 125.9, 124.7, 118.5, 113.0, 111.3, 108.9, 26.7, 13.6, 11.5. HRMS m/z [M+H]^+^ calcd for C_17_H_17_N_2_O_2_: 281.1290, found 281.1278, LC t_R_ = 4.62 min, > 98% Purity.

##### (3Z)-5-acetyl-3-(1H-pyrazol-5-ylmethylidene)-2,3-dihydro-1H-indol-2-one (29)

Compound **29** was prepared by procedure A to afford a mustard solid (33 mg, 0.130 mmol, 15%). MP 240-242 °C; ^1^H NMR (600 MHz, DMSO-*d*_6_) δ 13.76 (s, 1H), 10.94 (s, 1H), 9.75 (s, 1H), 7.96 (s, 1H), 7.92 (dd, *J* = 8.2, 1.8 Hz, 1H), 7.56 (s, 1H), 6.97 (d, *J* = 8.1 Hz, 1H), 6.86 (s, 1H), 2.58 (s, 3H). ^13^C NMR (150 MHz, DMSO-*d*_6_) δ 196.6, 170.0, 146.5, 146.2, 130.7, 130.1, 127.2, 127.0, 124.0, 121.8, 120.4, 111.0, 109.1, 26.5. HRMS m/z [M+H]^+^ calcd for C_14_H_12_N_3_O_2_: 254.0930, found 254.0920, LC t_R_ = 3.30 min, > 98% Purity.

##### (Z)-5-acetyl-3-(1H-imidazol-2-ylmethylidene)-2,3-dihydro-1H-indol-2-one (30)

Compound **30** was prepared by procedure A to afford a bright yellow solid (134 mg, 0.529 mmol, 62%, *E*:Z ratio 47:53 - not able to distinguish E/Z isomers). MP >300 °C; Major diastereoisomer. 3-((1*H*-Imidazol-2-yl)methylene)-5-acetylindolin-2-one. ^1^H NMR (600 MHz, DMSO-*d*_6_) δ 10.94 (bs, 1H), 10.14 (d, *J* = 1.8 Hz, 1H), 7.91 (dd, *J* = 8.2, 1.9 Hz, 1H), 7.56 (bs, 1H), 7.48 (bs, 1H), 7.46 (s, 1H), 6.96 (d, *J* = 8.2 Hz, 1H), 2.58 (s, 3H). Minor diastereoisomer. 3-((1*H*-Imidazol-2-yl)methylene)-5-acetylindolin-2-one. ^1^H NMR (600 MHz, DMSO-*d*_6_) δ 13.97 (bs, 1H), 10.95 (bs, 1H), 8.54 (d, *J* = 1.6 Hz, 1H), 8.06 (s, 1H), 7.88 (dd, *J* = 8.2, 1.7 Hz, 1H), 7.59 (d, *J* = 2.1 Hz, 1H), 7.39 (d, *J* = 1.3 Hz, 1H), 7.03 (d, *J* = 8.1 Hz, 1H), 2.60 (s, 3H). HRMS m/z [M+H]^+^ calcd for C_14_H_12_N_3_O_2_: 254.0930, found 254.0917, LC t_R_ = 2.86 min, > 98% Purity.

##### (3Z)-5-acetyl-3-(pyridin-2-ylmethylidene)-2,3-dihydro-1H-indol-2-one (31)

Compound **31** was prepared by procedure A to afford a mustard solid (190 mg, 0.719 mmol, 63%). MP 250-252 °C; ^1^H NMR (400 MHz, DMSO-*d*_6_) δ 11.00 (s, 1H), 9.77 (d, *J* = 1.8 Hz, 1H), 8.88 (dt, *J* = 4.8, 1.2 Hz, 1H), 7.96 (td, *J* = 7.6, 1.8 Hz, 1H), 7.93 – 7.80 (m, 2H), 7.62 (s, 1H), 7.49 (ddd, *J* = 7.5, 4.7, 1.3 Hz, 1H), 6.94 (d, *J* = 8.2 Hz, 1H), 2.55 (s, 3H). ^13^C NMR (100 MHz, DMSO-*d*_6_) δ 196.4, 169.7, 152.9, 149.7, 147.6, 137.5, 134.9, 131.5, 130.6, 128.9, 128.9, 128.4, 124.6, 121.3, 109.5, 26.4. HRMS m/z [M+H]^+^ calcd for C_16_H_13_N_2_O_2_: 265.0977, found 265.0966, LC t_R_ = 3.72 min, > 98% Purity.

##### (3Z)-5-acetyl-3-(pyridin-3-ylmethylidene)-2,3-dihydro-1H-indol-2-one (32)

Compound **32** was prepared by procedure A to afford a yellow solid (88 mg, 0.333 mmol, 39%, *E*:Z ratio 48:52 - not able to distinguish E/Z isomers). MP 209-211 °C; Major diastereoisomer. 5-Acetyl-3-(pyridin-3-ylmethylene)indolin-2-one. ^1^H NMR (600 MHz, DMSO-*d*_6_) δ 11.10 (bs, 1H), 9.24 (d, *J* = 2.2 Hz, 1H), 8.93 – 8.91 (m, 2H), 8.61 (dd, *J* = 4.8, 1.7 Hz, 1H), 8.08 (s, 1H), 7.94 – 7.91 (m, 1H), 7.51 (dd, *J* = 8.1, 4.8 Hz, 1H), 6.94 (d, *J* = 8.2 Hz, 1H), 2.58 (s, 3H). Minor diastereoisomer. 5-Acetyl-3-(pyridin-3-ylmethylene)indolin-2-one. ^1^H NMR (600 MHz, DMSO-*d*_6_) δ 11.08 (bs, 1H), 8.69 (dd, *J* = 4.8, 1.6 Hz, 1H), 8.38 (d, *J* = 1.6 Hz, 1H), 8.23 – 8.15 (m, 1H), 7.99 (d, *J* = 1.7 Hz, 1H), 7.94 – 7.91 (m, 1H), 7.73 (s, 1H), 7.59 (dd, *J* = 7.7, 5.0 Hz, 1H), 6.99 (d, *J* = 8.2 Hz, 1H), 2.42 (s, 3H). HRMS m/z [M+H]^+^ calcd for C_16_H_13_N_2_O_2_: 265.0977, found 265.0967, LC t_R_ = 2.46 min, > 98% Purity.

##### (3Z)-2-oxo-3-(1H-pyrrol-2-ylmethylidene)-2,3-dihydro-1H-indole-5-carboxylic acid (33)

Compound **33** was prepared by procedure A to afford a dark orange solid (90 mg, 0.354 mmol, 42%, E/Z ratio 30:70). MP >220 °C (decomp.); (*Z*)-3-((1*H-*pyrrol-2-yl)methylene)-2-oxoindoline-5-carboxylic acid. ^1^H NMR (600 MHz, DMSO-*d*_6_) δ 13.28 (bs, 1H), 11.07 (bs, 1H), 8.18 (d, *J* = 1.5 Hz, 1H), 7.77 (dd, *J* = 8.0, 1.4 Hz, 1H), 7.77 (s, 1H), 7.34 – 7.33 (m, 1H), 6.88 – 6.85 (m, 1H), 6.83 (d, *J* = 8.0 Hz, 1H), 6.34 (dt, *J* = 3.7, 2.3 Hz, 1H). ^13^C NMR (125 MHz, DMSO-*d*_6_) δ 170.3, 169.7, 140.1, 129.7 (s, 2C), 128.6, 126.1, 125.4, 124.1, 120.3, 119.5, 117.1, 111.3, 108.3. (*E*)-3-((1*H-*pyrrol-2-yl)methylene)-2-oxoindoline-5-carboxylic acid. ^1^H NMR (600 MHz, DMSO-*d*_6_) δ 11.91 (bs, 1H), 10.58 (bs, 1H), 8.59 (d, *J* = 1.5 Hz, 1H), 7.81 (dd, *J* = 8.0, 1.4 Hz, 1H), 7.52 (s, 1H), 7.21 – 7.20 (m, 1H), 7.15 – 7.05 (m, 1H), 6.80 (d, *J* = 8.0 Hz, 1H), 6.42 – 6.41 (m, 1H). ^13^C NMR (125 MHz, DMSO-*d*_6_) δ 170.3, 169.7 142.6, 132.2, 129.8 (s, 2C), 127.6, 124.8, 124.0, 123.1, 120.9, 113.5, 111.5, 108.2. HRMS m/z [M+H]^+^ calcd for C_14_H_11_N_2_O_3_: 255.0770, found 255.0762, LC t_R_ = 3.79 min, > 98% Purity.

##### (3Z)-3-(1H-imidazol-5-ylmethylidene)-2-oxo-2,3-dihydro-1H-indole-5-carboxylic acid (34)

Compound **34** was prepared by procedure A to afford a yellow solid (104 mg, 0.408 mmol, 48%). MP >300 °C; ^1^H NMR (600 MHz, DMSO-*d*_6_) δ 10.75 (s, 1H), 9.90 (s, 1H), 8.25 (s, 1H), 8.07 (s, 1H), 7.96 (s, 1H), 7.83 (dd, *J* = 8.1, 1.7 Hz, 1H), 7.54 (s, 1H), 6.89 (d, *J* = 8.1 Hz, 1H). ^13^C NMR (150 MHz, DMSO-*d*_6_) δ 170.3, 167.9, 145.3, 138.3, 130.6, 129.8, 128.2, 126.6, 124.4, 123.9, 122.4, 120.6, 108.6. HRMS m/z [M+H]^+^ calcd for C_13_H_10_N_3_O_3_: 256.0722, found 256.0716, LC t_R_ = 2.46 min, > 98% Purity.

##### (3Z)-3-(1H-imidazol-2-ylmethylidene)-2-oxo-2,3-dihydro-1H-indole-5-carboxylic acid (35)

Compound **35** was prepared by procedure A to afford a yellow solid (113 mg, 0.443 mmol, 52%). MP >260 °C (decomp.); ^1^H NMR (600 MHz, DMSO-*d*_6_) δ 14.34 (s, 1H), 8.15 (d, *J* = 1.4 Hz, 1H), 7.83 (dd, *J* = 8.0, 1.4 Hz, 1H), 7.57 (s, 1H), 7.53 (s, 1H), 7.36 – 7.25 (m, 1H), 6.81 (d, *J* = 7.9 Hz, 1H). ^13^C NMR (150 MHz, DMSO-*d*_6_) δ 174.4, 169.3, 143.9, 137.0, 134.7, 132.2, 130.5, 128.2, 125.0, 123.0, 122.8, 120.5, 108.8. HRMS m/z [M+H]^+^ calcd for C_13_H_10_N_3_O_3_: 256.0722, found 256.0714, LC t_R_ = 2.46 min, > 98% Purity.

##### 5-butyrylindolin-2-one (36)

Compound **36** was prepared by procedure B to afford a colourless solid (901 mg, 4.431 mmol, 59%). MP 130-132 °C; ^1^H NMR (400 MHz, DMSO-*d*_6_) δ 10.73 (s, 1H), 7.89 – 7.82 (m, 1H), 7.82 – 7.76 (m, 1H), 6.89 (d, *J* = 8.1 Hz, 1H), 3.54 (s, 2H), 2.90 (t, *J* = 7.2 Hz, 2H), 1.61 (h, *J* = 7.3 Hz, 2H), 0.91 (t, *J* = 7.4 Hz, 3H). ^13^C NMR (100 MHz, DMSO-*d*_6_) δ 198.5, 176.8, 148.2, 130.4, 129.0, 126.1, 124.1, 108.7, 39.4, 35.5, 17.6, 13.7. HRMS m/z [M+H]^+^ calcd for C_12_H_14_NO_2_: 204.1025,found 204.1017, LC t_R_ = 3.18 min, > 98% Purity.

##### 5-benzoylindolin-2-one (37)

Compound **37** was prepared by procedure C to afford a colourless solid (1.319 g, 5.555 mmol, 74%). MP 186-188 °C; ^1^H NMR (400 MHz, DMSO-*d*_6_) δ 10.81 (s, 1H), 7.73 – 7.59 (m, 5H), 7.57 – 7.51 (m, 2H), 6.95 (d, *J* = 8.6 Hz, 1H), 3.57 (s, 2H). ^13^C NMR (100 MHz, DMSO-*d*_6_) δ 194.7, 176.7, 148.3, 138.0, 131.9, 131.3, 130.0, 129.2 (s, 2C), 128.4 (s, 2C), 126.2, 126.0, 108.7, 35.6. HRMS m/z [M+H]^+^ calcd for C_15_H_12_NO_2_: 238.0868, found 238.0858, LC t_R_ = 3.36 min, > 98% Purity.

##### 5-(4-fluorobenzoyl)indolin-2-one (38)

Compound **38** was prepared by procedure B to afford a colourless/light green solid (571 mg, 2.237 mmol, 30%). MP 155-157 °C; ^1^H NMR (400 MHz, DMSO-*d*_6_) δ 10.86 (s, 1H), 7.76 – 7.55 (m, 3H), 7.51 (td, J = 7.4, 1.7 Hz, 1H), 7.40 – 7.31 (m, 2H), 6.95 (d, J = 8.6 Hz, 1H), 3.57 (s, 2H). 13C NMR (100 MHz, DMSO-*d*_6_) δ 191.4, 176. 8, 158.8 (d, J = 247.6 Hz), 149.3, 132.8 (d, J = 8.2 Hz), 131.2, 130.0 (d, J = 14.9 Hz), 129.9, 127.20 (d, J = 16.1 Hz), 126.5, 125.5, 124.7 (d, J = 3.4 Hz), 116.1 (d, J = 21.4 Hz), 109.0, 35.5. HRMS m/z [M+H]^+^ calcd for C_15_H_11_FNO_2_: 256.0774, found 256.0763, LC t_R_ = 3.40 min, > 98% Purity.

##### 5-(furan-2-carbonyl)indolin-2-one (39)

Compound **39** was prepared by procedure B to afford a beige solid (836 mg, 3.579 mmol, 49%). Consistent with previous report.^86^

##### (3Z)-5-butanoyl-3-(1H-pyrrol-2-ylmethylidene)-2,3-dihydro-1H-indol-2-one (40)

Compound **40** was prepared by procedure A to afford a mustard solid (43 mg, 0.153 mmol, 31%). MP 153-155 °C; ^1^H NMR (400 MHz, DMSO-*d*_6_) δ 13.22 (s, 1H), 11.23 (s, 1H), 8.30 (d, *J* = 1.6 Hz, 1H), 7.99 (s, 1H), 7.82 (dd, *J* = 8.2, 1.7 Hz, 1H), 7.40 (q, *J* = 1.8, 1.3 Hz, 1H), 6.97 (d, *J* = 8.2 Hz, 1H), 6.92 (dt, *J* = 3.6, 1.7 Hz, 1H), 6.39 (dt, *J* = 3.7, 2.3 Hz, 1H), 2.99 (t, *J* = 7.2 Hz, 2H), 1.66 (h, *J* = 7.3 Hz, 2H), 0.95 (t, *J* = 7.4 Hz, 3H). ^13^C NMR (100 MHz, DMSO-*d*_6_) δ 198.8, 169.6, 142.6, 130.6, 129.6, 127.8, 127.5, 126.5, 125.3, 121.3, 118.6, 115.6, 111.8, 109.2, 17.6 (s, 2C), 13.8. HRMS m/z [M+H]^+^ calcd for C_17_H_17_N_2_O_2_: 281.1290, found 281.1286, LC t_R_ = 4.85 min, > 98% Purity.

##### (3Z)-5-butanoyl-3-(1H-imidazol-5-ylmethylidene)-2,3-dihydro-1H-indol-2-one (41)

Compound **41** was prepared by procedure A to afford a bright yellow solid (58 mg, 0.207 mmol, 42%). MP >300 °C; ^1^H NMR (600 MHz, DMSO-*d*_6_) δ 10.52 (s, 1H), 10.36 (d, *J* = 1.9 Hz, 1H), 8.53 (s, 1H), 7.75 – 7.61 (m, 3H), 7.51 (s, 1H), 6.85 (d, *J* = 8.1 Hz, 1H), 2.96 (t, *J* = 7.3 Hz, 2H), 1.74 – 1.68 (m, 2H), 0.99 (t, *J* = 7.4 Hz, 3H). ^13^C NMR (150 MHz, DMSO-*d*_6_) δ 174.3, 170.5, 165.8, 156.3, 148.0, 145.3, 130.8, 124.3, 122.2, 121.8, 114.0, 111.9, 108.1, 48.6, 25.6, 14.0. HRMSm/z [M+H]^+^ calcd for C_16_H_16_N_3_O_2_: 282.1243, found 282.1233, LC t_R_ = 3.36 min, > 98% Purity.

##### (3Z)-5-butanoyl-3-(1H-imidazol-2-ylmethylidene)-2,3-dihydro-1H-indol-2-one (42)

Compound **42** was prepared by procedure A to afford a bright yellow solid (77 mg, 0.274 mmol, 56%). MP >300 °C; ^1^H NMR (600 MHz, DMSO-*d*_6_) δ 13.98 (s, 1H), 11.48 (s, 1H), 8.56 (d, *J* = 1.6 Hz, 1H), 8.08 (s, 1H), 7.89 (dd, *J* = 8.2, 1.7 Hz, 1H), 7.59 (d, *J* = 2.1 Hz, 1H), 7.39 (d, *J* = 1.1 Hz, 1H), 7.02 (d, *J* = 8.2 Hz, 1H), 3.03 (t, *J* = 7.2 Hz, 2H), 1.65 (q, *J* = 7.3 Hz, 2H), 0.96 (t, *J* = 7.4 Hz, 3H). ^13^C NMR (150 MHz, DMSO-*d*_6_) δ 199.0, 169.4, 143.6, 143.5, 132.9, 131.1, 129.2, 125.9, 124.1, 122.6, 121.2, 120.8, 109.9, 40.1, 17.6, 13.8. HRMS m/z [M+H]^+^ calcd for C_16_H_16_N_3_O_2_: 282.1243, found 282.1236, LC t_R_ = 3.29 min, > 98% Purity.

##### (3Z)-5-benzoyl-3-(1H-pyrrol-2-ylmethylidene)-2,3-dihydro-1H-indol-2-one (43)

Compound **43** was prepared by procedure A to afford a red solid (29 mg, 0.092 mmol, 22%). MP 232-234 °C; ^1^H NMR (400 MHz, DMSO-*d*_6_) δ 13.21 (s, 1H), 11.29 (s, 1H), 8.10 (d, *J* = 1.6 Hz, 1H), 7.97 (s, 1H), 7.78 – 7.70 (m, 2H), 7.69 – 7.64 (m, 1H), 7.61 – 7.53 (m, 3H), 7.41 (q, *J* = 1.8, 1.2 Hz, 1H), 7.03 (d, *J* = 8.1 Hz, 1H), 6.93 (dt, *J* = 3.6, 1.7 Hz, 1H), 6.38 (dt, *J* = 3.7, 2.3 Hz, 1H). ^13^C NMR (100 MHz, DMSO-*d*_6_) δ 195.1, 169.6, 142.6, 138.0, 132.1, 130.4, 130.0, 129.6, 129.4 (s, 2C), 128.5 (s, 2C), 128.1, 126.6, 125.4, 121.6, 119.9, 115.2, 111.8, 109.1. HRMS m/z [M+H]^+^ calcd for C_20_H_15_N_2_O_2_: 315.1134, found 315.1120, LC t_R_ = 4.90 min, > 98% Purity.

##### (3Z)-5-benzoyl-3-(1H-imidazol-5-ylmethylidene)-2,3-dihydro-1H-indol-2-one (44)

Compound **44** was prepared by procedure A to afford a dark orange solid (24 mg, 0.076 mmol, 24%). MP 255-257 °C; ^1^H NMR (400 MHz, DMSO-*d*_6_) δ 10.86 (s, 1H), 9.96 (s, 1H), 7.95 (d, *J* = 1.0 Hz, 1H), 7.78 – 7.71 (m, 4H), 7.69 – 7.55 (m, 5H), 6.99 (d, *J* = 8.1 Hz, 1H). ^13^C NMR (100 MHz, DMSO-*d*_6_) δ 194.9, 170.3, 145.6, 138.3, 137.8, 136.9, 131.7, 131.3, 129.9, 129.5, 129.3 (s, 2C), 128.6, 128.2 (s, 2C), 126.6, 122.34, 120.6, 108.9. HRMS m/z [M+H]^+^ calcd for C_19_H_14_N_3_O_2_: 316.1086, found 316.1076, LC t_R_ = 3.51 min, > 98% Purity.

##### (3Z)-5-[(4-fluorophenyl)carbonyl]-3-(1H-pyrrol-2-ylmethylidene)-2,3-dihydro-1H-indol-2-one (45)

Compound **45** was prepared by procedure A to afford a dark orange solid (37 mg, 0.111 mmol, 28%). MP 242-244 °C; ^1^H NMR (400 MHz, DMSO-*d*_6_) δ 13.19 (s, 1H), 11.35 (s, 1H), 8.14 (d, *J* = 1.6 Hz, 1H), 7.98 (s, 1H), 7.65 (ddd, *J* = 13.9, 5.5, 2.7 Hz, 1H), 7.62 – 7.51 (m, 2H), 7.46 – 7.30 (m, 3H), 7.02 (d, *J* = 8.1 Hz, 1H), 6.94 (dt, *J* = 3.7, 1.7 Hz, 1H), 6.39 (dt, *J* = 4.1, 2.3 Hz, 1H). ^13^C NMR (100 MHz, DMSO-*d*_6_) δ 191.7, 169.6, 159.0 (d, *J* = 248.1 Hz), 143.5, 132.9 (d, *J* = 8.2 Hz), 130.4, 130.2 (d, *J* = 3.0 Hz), 130.0, 129.6, 128.3, 127.3 (d, *J* = 15.7 Hz), 126.8, 125.7, 124.7 (d, *J* = 3.4 Hz), 121.8, 119.3, 116.2 (d, *J* = 21.4 Hz), 114.9, 111.8, 109.3. HRMS m/z [M+H]^+^ calcd for C_20_H_14_N_2_O_2_F: 333.1039, found 333.1035, LC t_R_ = 4.88 min, > 98% Purity.

##### (3Z)-5-[(4-fluorophenyl)carbonyl]-3-(1H-imidazol-5-ylmethylidene)-2,3-dihydro-1H-indol-2-one (46)

Compound **46** was prepared by procedure A to afford a bright yellow solid (51 mg, 0.153 mmol, 39%). MP >300 °C; ^1^H NMR (600 MHz, DMSO-*d*_6_) δ 10.04 (d, *J* = 1.6 Hz, 1H), 7.98 (d, *J* = 39.4 Hz, 1H), 7.75 (s, 1H), 7.70 (dd, *J* = 8.2, 1.9 Hz, 1H), 7.66 (tdd, *J* = 7.5, 5.3, 1.8 Hz, 1H), 7.53 (dd, *J* = 7.2, 1.9 Hz, 1H), 7.50 (d, *J* = 5.3 Hz, 1H), 7.42 (s, 1H), 7.39 (d, *J* = 3.9 Hz, 1H), 7.37 (d, *J* = 7.5 Hz, 1H), 6.94 (d, *J* = 8.2 Hz, 1H). ^13^C NMR (150 MHz, DMSO-*d*_6_) δ 192.1, 173.7, 170.8, 158.9 (d, *J* = 246.5 Hz), 145.3, 137.1, 132.1 (d, *J* = 8.5 Hz), 129.9, 129.8 (d, *J* = 3.6 Hz, 2C), 129.6, 129.6, 128.2 (d, *J* = 16.6 Hz, 2C), 128.1, 124.5 (d, *J* = 3.5 Hz), 123.6, 115.9 (d, *J* = 21.5 Hz), 108.6. HRMS m/z [M+H]^+^ calcd for C_19_H_13_N_3_O_2_F: 334.0992, found 334.0979, LC t_R_ = 3.57 min, > 98% Purity.

##### (Z)-3-((1H-pyrrol-2-yl)methylene)-5-(furan-2-carbonyl)indolin-2-one (47)

Compound **47** was prepared by procedure A to afford a yellow solid (19 mg, 0.062 mmol, 14%). MP >220 °C (decomp.); ^1^H NMR (600 MHz, DMSO-*d*_6_) δ 13.82 (s, 1H), 8.53 (s, 1H), 8.13 (s, 1H), 8.08 (d, *J* = 1.6 Hz, 1H), 7.83 (s, 1H), 7.78 (dd, *J* = 8.2, 1.7 Hz, 1H), 7.37 (d, *J* = 3.5 Hz, 1H), 7.33 (s, 1H), 6.95 (d, *J* = 8.1 Hz, 1H), 6.91 – 6.80 (m, 1H), 6.78 (dd, *J* = 3.6, 1.7 Hz, 1H), 6.34 (d, *J* = 3.0 Hz, 1H). ^13^C NMR (150 MHz, DMSO-*d*_6_) δ 180.6, 174.2, 165.6, 151.9, 147.6, 133.6, 130.0, 129.2, 126.6, 125.4, 119.8, 118.9, 116.2, 115.5, 114.1, 112.4, 111.3, 109.9. HRMS m/z [M+H]^+^ calcd for C_18_H_13_N_2_O_3_: 305.0926, found 305.0920, LC t_R_ = 4.44 min, > 98% Purity.

##### (3Z)-5-[(furan-2-yl)carbonyl]-3-(1H-imidazol-2-ylmethylidene)-2,3-dihydro-1H-indol-2-one (48)

Compound **48** was prepared by procedure A to afford a yellow solid (23 mg, 0.075 mmol, 17%). MP >270 °C (decomp.); ^1^H NMR (600 MHz, DMSO-*d*_6_) δ 10.73 (s, 2H), 10.55 (s, 1H), 8.53 (s, 1H), 8.09 (d, *J* = 1.6 Hz, 1H), 7.81 (dd, *J* = 8.1, 2.0 Hz, 1H), 7.73 (d, *J* = 3.6 Hz, 1H), 7.53 (s, 1H), 7.36 (s, 1H), 6.96 (d, *J* = 8.1 Hz, 1H), 6.85 (dd, *J* = 3.5, 1.7 Hz, 1H). ^13^C NMR (150 MHz, DMSO-*d*_6_) δ 180.4, 173.9, 165.5, 151.6, 147.5, 131.8, 129.7, 129.2, 127.6, 122.5, 120.1, 117.3, 114.9, 114.0, 112.4, 111.2, 108.6. HRMS m/z [M+H]^+^ calcd for C_17_H_12_N_3_O_3_: 306.0879, found 306.0867, LC t_R_ = 2.96 min, > 98% Purity.

##### *N,N*-dimethyl-2-oxoindoline-5-sulfonamide (50)

Compound **50** was prepared by procedure D to afford a beige solid (447 mg, 1.985 mmol, 55%). MP 198-200 °C; ^1^H NMR (400 MHz, DMSO-*d*_6_) δ 10.82 (s, 1H), 7.67 – 7.46 (m, 2H), 7.04 – 6.96 (m, 1H), 3.60 (s, 2H), 2.57 (s, 6H). ^13^C NMR (100 MHz, DMSO-*d*_6_) δ 176.4, 148.0, 128.4, 126.9, 126.9, 123.7, 109.1, 37.6 (s, 2C), 35.7. HRMS m/z [M+H]^+^ calcd for C_10_H_13_N_2_O_3_S: 241.0649, found 241.0633.

##### 5-((4-methylpiperazin-1-yl)sulfonyl)indolin-2-one (51)

Compound **51** was prepared by procedure D to afford a beige/brown solid (496 mg, 1.679 mmol, 39%). MP 170-172 °C; HRMS m/z [M+H]^+^ calcd for C_13_H_18_N_3_O_3_S: 296.1069, found 296.1054. Consistent with previous report.^88^

##### (3Z)-*N,N*-dimethyl-2-oxo-3-(1H-pyrrol-2-ylmethylidene)-2,3-dihydro-1H-indole-5-sulfonamide (52)

Compound **52** was prepared by procedure A to afford an orange solid (54 mg, 0.170 mmol, 41%). MP 258-260 °C; ^1^H NMR (400 MHz, DMSO-*d*_6_) δ 13.23 (s, 1H), 11.33 (s, 1H), 8.11 (s, 1H), 8.05 (d, *J* = 1.7 Hz, 1H), 7.53 (dd, *J* = 8.2, 1.8 Hz, 1H), 7.44 (q, *J* = 2.3 Hz, 1H), 7.09 (d, *J* = 8.2 Hz, 1H), 6.94 (dt, *J* = 3.7, 1.7 Hz, 1H), 6.41 (dt, *J* = 4.1, 2.3 Hz, 1H), 2.61 (s, 6H). ^13^C NMR (100 MHz, DMSO-*d*_6_) δ 169.3, 142.1, 129.6, 128.9, 127.1, 127.1, 126.6, 125.9, 122.0, 117.9, 114.7, 112.0, 109.5, 37.7 (2C, s). HRMS m/z [M+H]^+^ calcd for C_15_H_16_N_3_O_3_S: 318.0912, found 318.0897, LC t_R_ = 4.10 min, > 98% Purity.

##### (3Z)-3-(1H-imidazol-5-ylmethylidene)-*N,N*-dimethyl-2-oxo-2,3-dihydro-1H-indole-5-sulfonamide (53)

Compound **53** was prepared by procedure A to afford an orange solid (52 mg, 0.217 mmol, 52%). MP 275-277 °C; ^1^H NMR (400 MHz, DMSO-*d*_6_) δ 12.81 (s, 1H), 10.87 (s, 1H), 9.92 (d, *J* = 1.9 Hz, 1H), 8.07 (t, *J* = 0.9 Hz, 1H), 8.02 (d, *J* = 1.1 Hz, 1H), 7.62 (d, *J* = 0.7 Hz, 1H), 7.58 (dd, *J* = 8.2, 2.0 Hz, 1H), 7.04 – 6.99 (m, 1H), 2.65 (s, 6H). ^13^C NMR (100 MHz, DMSO-*d*_6_) δ 170.0, 145.1, 138.3, 136.9, 129.4, 128.4, 127.14, 127.09, 125.9, 123.0, 120.1, 108.9, 37.8. HRMS m/z [M+H]^+^ calcd for C_14_H_15_N_4_O_3_S: 319.0865, found 319.0850, LC t_R_ = 2.54 min, > 98% Purity.

##### (3Z)-5-[(4-methylpiperazin-1-yl)sulfonyl]-3-(1H-pyrrol-2-ylmethylidene)-2,3-dihydro-1H-indol-2-one (54)

Compound **54** was prepared by procedure A to afford a bright yellow solid (82 mg, 0.220 mmol, 65%). MP >260 °C (decomp.); ^1^H NMR (400 MHz, DMSO-*d*_6_) δ 13.23 (s, 1H), 11.35 (s, 1H), 8.09 (s, 1H), 8.02 (d, *J* = 1.7 Hz, 1H), 7.51 (dd, *J* = 8.2, 1.8 Hz, 1H), 7.44 (q, *J* = 2.2 Hz, 1H), 7.09 (d, *J* = 8.1 Hz, 1H), 6.95 (dt, *J* = 3.6, 1.7 Hz, 1H), 6.41 (dt, *J* = 4.0, 2.3 Hz, 1H), 2.89 (t, *J* = 4.7 Hz, 4H), 2.36 (t, *J* = 4.9 Hz, 4H), 2.13 (s, 3H). ^13^C NMR (100 MHz, DMSO-*d*_6_) δ 169.3, 142.2, 129.6, 129.0, 127.2, 127.2, 126.6, 125.9, 122.1, 117.8, 114.6, 112.0, 109.5, 53.5, 45.8 (s, 2C), 45.3 (s, 2C). HRMS m/z [M+H]^+^ calcd for C_18_H_21_N_4_O_3_S: 373.1334, found 373.1326, LC t_R_ = 3.10 min, > 98% Purity.

##### (3Z)-3-(1H-imidazol-5-ylmethylidene)-5-[(4-methylpiperazin-1-yl)sulfonyl]-2,3-dihydro-1H-indol-2-one (55)

Compound **55** was prepared by procedure A to afford a bright yellow solid (67 mg, 0.228 mmol, 67%). MP >260 °C (decomp.); ^1^H NMR (600 MHz, DMSO-*d*_*6*_) δ 10.85 (s, 1H), 9.96 (d, *J* = 1.9 Hz, 1H), 7.97 (s, 1H), 7.94 (s, 1H), 7.59 (s, 1H), 7.52 (dd, *J* = 8.1, 2.0 Hz, 1H), 6.99 (d, *J* = 8.1 Hz, 1H), 2.95 (s, 4H), 2.36 (d, *J* = 5.2 Hz, 4H), 2.12 (s, 3H). ^13^C NMR (150 MHz, DMSO-*d*_*6*_) δ 170.2, 144.7, 140.9, 137.0, 129.9, 127.8, 127.7, 127.0, 125.6, 123.4, 118.5, 108.6, 53.7 (s, 2C), 45.9 (s, 2C), 45.3. HRMS m/z [M+H]^+^ calcd for C_17_H_20_N_5_O_3_S: 374.1287, found 374.1276, LC t_R_ = 2.29 min, > 98% Purity.

##### (Z)-3-((1H-pyrrol-2-yl)methylene)-5-isopropoxyindolin-2-one (57)

Compound **57** was prepared by procedure A to afford a red solid (48 mg, 0.179 mmol, 68%). MP 149-151 °C; ^1^H NMR (400 MHz, DMSO-*d*_6_) δ 13.39 (s, 1H), 10.68 (s, 1H), 7.76 (s, 1H), 7.34 (td, *J* = 2.6, 1.3 Hz, 1H), 7.31 (d, *J* = 2.3 Hz, 1H), 6.79 (dt, *J* = 3.6, 1.7 Hz, 1H), 6.75 (d, *J* = 8.3 Hz, 1H), 6.70 (dd, *J* = 8.4, 2.3 Hz, 1H), 6.35 (dt, *J* = 3.6, 2.3 Hz, 1H), 4.54 (p, *J* = 6.0 Hz, 1H), 1.26 (s, 3H), 1.25 (s, 3H). ^13^C NMR (100 MHz, DMSO-*d*_6_) δ 169.3, 152.8, 132.9, 129.5, 126.6, 126.2, 125.5, 120.1, 117.4, 115.4, 111.4, 110.0, 107.0, 70.1, 22.0 (s, 2C). HRMS m/z [M+H]^+^ calcd for C_16_H_17_N_2_O_2_: 269.1290, found 269.1283, LC t_R_ = 4.97 min, > 98% Purity.

##### (3Z)-3-(1H-imidazol-5-ylmethylidene)-5-(propan-2-yloxy)-2,3-dihydro-1H-indol-2-one (58)

Compound **58** was prepared by procedure A to afford an orange solid (37 mg, 0.137 mmol, 53%). MP 211-213 °C; ^1^H NMR (400 MHz, DMSO-*d*_6_) δ 13.79 (s, 1H), 10.81 (s, 1H), 8.01 (s, 1H), 7.87 (s, 1H), 7.59 (s, 1H), 7.36 – 7.34 (m, 1H), 6.76 (d, *J* = 3.0 Hz, 2H), 4.57 – 4.51 (m, 1H), 1.27 (s, 3H), 1.25 (s, 3H). ^13^C NMR (100 MHz, DMSO-*d*_6_) δ ^13^C NMR (100 MHz, DMSO-*d*_6_) δ 169.1, 153.0, 139.5, 138.1, 133.6, 128.1, 125.4, 123.1, 120.6, 116.5, 110.3, 107.7, 70.1, 22.0. HRMS m/z [M+H]^+^ calcd for C_15_H_16_N_3_O_2_: 270.1243, found 270.1230, LC t_R_ = 3.42 min, > 98% Purity.

##### (3Z)-3-(1H-imidazol-2-ylmethylidene)-5-(propan-2-yloxy)-2,3-dihydro-1H-indol-2-one (59)

Compound **59** was prepared by procedure A to afford a bright orange solid (44 mg, 0.163 mmol, 62%). MP 256-258 °C; ^1^H NMR (400 MHz, DMSO-*d*_6_) δ 10.30 (s, 1H), 9.17 (d, *J* = 2.6 Hz, 1H), 7.85 (s, 1H), 7.55 – 7.34 (m, 3H), 6.79 (d, *J* = 1.4 Hz, 2H), 4.59 (p, *J* = 6.0 Hz, 1H), 1.30 – 1.25 (m, 6H). ^13^C NMR (100 MHz, DMSO-*d*_6_) δ 169.0, 153.2, 143.7, 133.7, 132.5, 125.0, 124.7, 124.2, 120.6, 117.4, 110.6, 108.3, 69.9, 22.0 (s, 2C). HRMS m/z [M+H]^+^ calcd for C_15_H_16_N_3_O_2_: 270.1243, found 270.1230, LC t_R_ = 3.29 min, > 98% Purity.

##### *N*-(2-oxoindolin-5-yl)butyramide (60)

Compound **60** was prepared by procedure F to afford a colourless solid (552 mg, 2.531 mmol, 75%). MP 165-167 °C; ^1^H NMR (600 MHz, DMSO-*d*_6_) δ 10.26 (s, 1H), 9.70 (s, 1H), 7.55 – 7.47 (m, 1H), 7.32 (dd, *J* = 8.3, 2.0 Hz, 1H), 6.71 (d, *J* = 8.3 Hz, 1H), 3.44 (s, 2H), 2.23 (t, *J* = 7.3 Hz, 2H), 1.59 (h, *J* = 7.4 Hz, 2H), 0.90 (t, *J* = 7.4 Hz, 3H). ^13^C NMR (150 MHz, DMSO-*d*_*6*_) δ 176.3, 170.6, 139.1, 133.5, 126.0, 118.4, 116.5, 108.8, 38.3, 36.1, 18.7, 13.6. HRMS m/z [M+H]^+^ calcd for C_12_H_15_N_2_O_2_: 219.1134, found 219.1125, LC t_R_ = 2.51 min, > 98% Purity.

##### 2-ethyl-*N*-(2-oxoindolin-5-yl)butanamide (61)

Compound **61** was prepared by procedure F to afford a colourless solid (590 mg, 2.395 mmol, 71%). MP 228-230 °C; ^1^H NMR (400 MHz, DMSO-*d*_6_) δ 10.26 (s, 1H), 9.68 (s, 1H), 7.54 (d, *J* = 2.0 Hz, 1H), 7.33 (dd, *J* = 8.4, 2.1 Hz, 1H), 6.72 (d, *J* = 8.3 Hz, 1H), 3.45 (s, 2H), 2.16 (tt, *J* = 9.4, 5.1 Hz, 1H), 1.59 – 1.48 (m, 2H), 1.47 – 1.36 (m, 2H), 0.84 (t, *J* = 7.4 Hz, 6H). ^13^C NMR (100 MHz, DMSO-d6) δ ^13^C NMR (100 MHz, DMSO-*d*_6_) δ 176.3, 173.4, 139.2, 133.3, 126.0, 118.69, 116.8, 108.8, 49.8, 36.0, 25.3 (s, 2C), 11.9 (s, 2C). HRMS m/z [M+H]^+^ calcd for C_14_H_19_N_2_O_2_: 247.1447, found 247.1436, LC t_R_ = 3.06 min, > 98% Purity.

##### *N*-(2-oxoindolin-5-yl)cyclohexanecarboxamide (62)

Compound **62** was prepared by procedure F to afford a colourless solid (780 mg, 2.632 mmol, 78%). MP 223-225 °C; ^1^H NMR (600 MHz, DMSO-*d*_6_) δ 10.25 (s, 1H), 9.62 (s, 1H), 7.57 – 7.45 (m, 1H), 7.33 (dd, *J* = 8.4, 2.1 Hz, 1H), 6.70 (d, *J* = 8.3 Hz, 1H), 3.44 (s, 2H), 2.27 (tt, *J* = 11.7, 3.3 Hz, 1H), 2.03 – 1.67 (m, 4H), 1.68 – 1.60 (m, 1H), 1.39 (qd, *J* = 13.4, 12.9, 3.7 Hz, 2H), 1.29 – 1.13 (m, 3H). ^13^C NMR (150 MHz, DMSO-*d*_6_) δ 176.3, 173.8, 139.0, 133.6, 125.9, 118.4, 116.5, 108.8, 44.8, 36.0, 29.2 (s, 2C), 25.42, 25.26 (s, 2C). HRMS m/z [M+H]^+^ calcd for C_15_H_19_N_2_O_2_: 259.1447, found 259.1435, LC t_R_ = 3.25 min, > 98% Purity.

##### *N*-(2-oxoindolin-5-yl)benzamide (63)

Compound **63** was prepared by procedure F to afford a colourless solid (647 mg, 2.565 mmol, 76%). MP 115-117 °C; ^1^H NMR (600 MHz, DMSO-*d*_6_) δ 10.37 (s, 1H), 10.14 (s, 1H), 7.99 – 7.90 (m, 2H), 7.67 (d, *J* = 2.0 Hz, 1H), 7.59 – 7.55 (m, 1H), 7.55 – 7.44 (m, 3H), 6.79 (d, *J* = 8.3 Hz, 1H), 3.49 (s, 2H). ^13^C NMR (150 MHz, DMSO-*d*_*6*_) 176.4, 165.2, 139.8, 135.1, 133.2, 131.4, 128.4 (s, 2C), 127.6 (s, 2C), 125.9, 119.9, 117.8, 108.8, 36.1. HRMS m/z [M+H]^+^ calcd for C_15_H_13_N_2_O_2_: 253.0977, found 253.0967, LC t_R_ = 2.93 min, > 98% Purity.

##### *N*-(2-oxoindolin-5-yl)furan-2-carboxamide (64)

Compound **55** was prepared by procedure F to afford a beige solid (605 mg, 2.498 mmol, 74%). ^1^H NMR (600 MHz, DMSO-*d*_6_) δ 10.35 (s, 1H), 10.07 (s, 1H), 7.90 (d, *J* = 1.6 Hz, 1H), 7.64 (d, *J* = 2.2 Hz, 1H), 7.49 (dd, *J* = 8.4, 2.1 Hz, 1H), 7.29 (d, *J* = 3.4 Hz, 1H), 6.78 (d, *J* = 8.3 Hz, 1H), 6.68 (dd, *J* = 3.5, 1.7 Hz, 1H), 3.49 (s, 2H). ^13^C NMR (150 MHz, DMSO-*d*_6_) δ 176.4, 156.0, 147.7, 145.5, 139.9, 132.4, 126.0, 120.0, 117.8, 114.3, 112.1, 108.8, 36.1. HRMS m/z [M+H]^+^ calcd for C_13_H_11_N_2_O_3_: 243.0770, found 243.0759, LC t_R_ = 2.51 min, > 98% Purity.

##### *N*-[(3Z)-2-oxo-3-(1H-pyrrol-2-ylmethylidene)-2,3-dihydro-1H-indol-5-yl]butanamide (65)

Compound **65** was prepared by procedure A to afford a bright yellow solid (85 mg, 0.288 mmol, 63%). MP 260-262 °C; ^1^H NMR (400 MHz, DMSO-*d*_6_) δ 13.34 (s, 1H), 10.80 (s, 1H), 9.74 (s, 1H), 7.95 (d, *J* = 2.0 Hz, 1H), 7.59 (s, 1H), 7.35 (q, *J* = 2.2 Hz, 1H), 7.17 (dd, *J* = 8.3, 2.0 Hz, 1H), 6.92 (dt, *J* = 3.5, 1.6 Hz, 1H), 6.80 (d, *J* = 8.3 Hz, 1H), 6.35 (dt, *J* = 3.7, 2.3 Hz, 1H), 2.27 (t, *J* = 7.3 Hz, 2H), 1.62 (h, *J* = 7.4 Hz, 2H), 0.93 (t, *J* = 7.4 Hz, 3H). ^13^C NMR (100 MHz, DMSO-*d*_6_) δ 170.7, 169.3, 134.8, 133.5, 129.5, 126.1, 125.7, 125.1, 120.7, 118.8, 117.0, 111.4, 110.5, 109.4, 38.2, 18.7, 13.7. HRMS m/z [M+H]^+^ calcd for C_17_H_18_N_3_O_2_: 296.1399, found 296.1392, LC t_R_ = 4.14 min, > 98% Purity.

##### 2-ethyl-*N*-[(3Z)-2-oxo-3-(1H-pyrrol-2-ylmethylidene)-2,3-dihydro-1H-indol-5-yl]butanamide (66)

Compound **66** was prepared by procedure A to afford a yellow solid (86 mg, 0.266 mmol, 66%). MP >300 °C; ^1^H NMR (400 MHz, DMSO-*d*_6_) δ 13.32 (s, 1H), 10.79 (s, 1H), 9.72 (s, 1H), 8.02 – 7.92 (m, 1H), 7.60 (s, 1H), 7.33 (td, *J* = 2.6, 1.4 Hz, 1H), 7.16 (dd, *J* = 8.3, 2.0 Hz, 1H), 6.90 (dt, *J* = 3.6, 1.7 Hz, 1H), 6.83 – 6.73 (m, 1H), 6.33 (dt, *J* = 3.7, 2.3 Hz, 1H), 2.18 (tt, *J* = 9.4, 5.1 Hz, 1H), 1.55 (ddq, *J* = 13.2, 9.0, 7.4 Hz, 2H), 1.48 – 1.35 (m, 2H), 0.86 (t, *J* = 7.4 Hz, 6H). ^13^C NMR (100 MHz, DMSO-*d*_6_) δ 173.5, 169.3, 134.9, 133.3, 129.5, 126.2, 125.7, 125.1, 120.7, 119.1, 117.0, 111.4, 110.8, 109.4, 49.8, 25.3 (2C, s), 11.9 (2C, s). HRMS m/z [M+H]^+^ calcd for C_19_H_22_N_3_O_2_: 324.1712, found 324.1702, LC t_R_ = 4.56 min, > 98% Purity.

##### 2-ethyl-*N*-[(3Z)-3-(1H-imidazol-5-ylmethylidene)-2-oxo-2,3-dihydro-1H-indol-5-yl]butanamide (67)

Compound **67** was prepared by procedure A to afford a yellow solid (70 mg, 0.216 mmol, 53%). MP 285-287 °C; ^1^H NMR (600 MHz, DMSO-*d*_*6*_) δ 13.74 (s, 1H), 9.84 (s, 1H), 8.02 (s, 1H), 7.98 (s, 1H), 7.85 (d, *J* = 22.8 Hz, 1H), 7.71 (s, 1H), 7.24 (dd, *J* = 8.2, 2.0 Hz, 1H), 6.82 (d, *J* = 8.2 Hz, 1H), 2.21 (tt, *J* = 9.5, 5.0 Hz, 1H), 1.56 (dddd, *J* = 13.5, 7.3, 5.8, 1.9 Hz, 2H), 1.44 (ddd, *J* = 13.0, 7.3, 5.2 Hz, 2H), 0.87 (t, *J* = 7.4 Hz, 6H). ^13^C NMR (150 MHz, DMSO-*d*_*6*_) δ 174.4, 173.6, 140.3, 135.8, 133.5, 132.5, 124.4, 121.2, 120.4, 120.0, 111.5, 109.6, 108.2, 49.7, 25.3 (s, 2C), 11.9 (s, 2C). HRMS m/z [M+H]^+^ calcd for C_18_H_21_N_4_O_2_: 325.1665, found 325.1660, LC t_R_ = 3.29 min, > 98% Purity.

##### 2-ethyl-*N*-[(3Z)-3-(1H-imidazol-2-ylmethylidene)-2-oxo-2,3-dihydro-1H-indol-5-yl]butanamide (68)

Compound **68** was prepared by procedure A to afford a yellow solid (74 mg, 0.228 mmol, 56%). MP >300 °C; ^1^H NMR (600 MHz, DMSO-*d*_6_) δ 14.06 (s, 1H), 11.08 (s, 1H), 9.80 (s, 1H), 7.99 (s, 1H), 7.56 (d, *J* = 4.2 Hz, 2H), 7.46 – 7.27 (m, 2H), 6.86 (d, *J* = 8.2 Hz, 1H), 2.20 (tt, *J* = 9.8, 5.0 Hz, 1H), 1.56 (dq, *J* = 15.5, 7.6 Hz, 2H), 1.45 (dq, *J* = 13.2, 6.7, 6.0 Hz, 2H), 0.87 (t, *J* = 7.5 Hz, 6H). ^13^C NMR (150 MHz, DMSO-*d*_*6*_) δ 173.7, 169.0, 143.4, 135.9, 133.8, 132.6, 124.0, 123.8, 123.7, 121.1, 120.9, 112.0, 110.0, 49.8, 25.3, 11.9. HRMS m/z [M+H]^+^ calcd for C_18_H_21_N_4_O_2_: 325.1665, found 325.1656, LC t_R_ = 3.24 min, > 98% Purity.

##### *N*-[(3Z)-2-oxo-3-(1H-pyrrol-2-ylmethylidene)-2,3-dihydro-1H-indol-5-yl]cyclohexanecarboxamide (69)

Compound **69** was prepared by procedure A to afford a yellow solid (71 mg, 0.274 mmol, 71%). MP >300 °C; ^1^H NMR (400 MHz, DMSO-*d*_6_) δ 13.37 – 13.27 (m, 1H), 10.78 (s, 1H), 9.66 (s, 1H), 7.96 (d, *J* = 1.9 Hz, 1H), 7.57 (s, 1H), 7.33 (td, *J* = 2.6, 1.3 Hz, 1H), 7.14 (dd, *J* = 8.3, 2.0 Hz, 1H), 6.89 (dt, *J* = 3.7, 1.7 Hz, 1H), 6.78 (d, *J* = 8.3 Hz, 1H), 6.33 (dt, *J* = 3.7, 2.3 Hz, 1H), 2.33 – 2.25 (m, 1H), 1.76 (td, *J* = 13.9, 3.4 Hz, 4H), 1.64 (d, *J* = 11.3 Hz, 1H), 1.41 (qd, *J* = 13.6, 13.1, 3.5 Hz, 2H), 1.31 – 1.14 (m, 3H). ^13^C NMR (100 MHz, DMSO-*d*_6_) δ 173.9, 169.3, 134.8, 133.6, 129.5, 126.1, 125.7, 125.0, 120.6, 118.8, 117.0, 111.4, 110.6, 109.4, 44.7, 29.2 (2C, s), 25.4, 25.3 (2C, s). HRMS m/z [M+H]^+^ calcd for C_20_H_22_N_3_O_2_: 336.1712, found 336.1703, LC t_R_ = 4.71 min, > 98% Purity.

##### *N*-[(3Z)-3-(1H-imidazol-5-ylmethylidene)-2-oxo-2,3-dihydro-1H-indol-5-yl]cyclohexanecarboxamide (70)

Compound **70** was prepared by procedure A to afford a yellow solid (89 mg, 0.265 mmol, 68%). >300 °C; ^1^H NMR (400 MHz, DMSO-*d*_6_) δ 13.70 (s, 1H), 10.89 (s, 1H), 9.71 (s, 1H), 8.00 (d, *J* = 8.4 Hz, 2H), 7.70 (s, 2H), 7.21 (d, *J* = 8.3 Hz, 1H), 6.81 (d, *J* = 8.3 Hz, 1H), 2.31 (tt, *J* = 11.7, 3.4 Hz, 1H), 2.08 – 1.67 (m, 4H), 1.65 (dd, *J* = 10.5, 4.0 Hz, 1H), 1.42 (qd, *J* = 13.7, 13.1, 3.5 Hz, 2H), 1.35 – 1.06 (m, 3H). ^13^C NMR (100 MHz, DMSO-*d*_6_) δ 173.9, 168.9, 146.2, 139.5, 138.6, 135.5, 133.8, 128.0, 124.3, 123.7, 119.8, 111.2, 109.7, 44.7, 29.2 (2C, s), 25.4, 25.3 (2C, s). HRMS m/z [M+H]^+^ calcd for C_19_H_21_N_4_O_2_: 337.1665, found 337.1657, LC t_R_ = 3.47 min, > 98% Purity.

##### *N*-[(3Z)-3-(1H-imidazol-2-ylmethylidene)-2-oxo-2,3-dihydro-1H-indol-5-yl]cyclohexanecarboxamide (71)

Compound **71** was prepared by procedure A to afford a yellow solid (81 mg, 0.241 mmol, 62%). MP >300 °C; ^1^H NMR (600 MHz, DMSO-*d*_6_) δ 14.05 (s, 1H), 11.07 (s, 1H), 9.74 (s, 1H), 7.98 (d, *J* = 2.0 Hz, 1H), 7.56 (d, *J* = 2.1 Hz, 1H), 7.51 (s, 1H), 7.38 (dd, *J* = 8.4, 2.0 Hz, 1H), 7.34 (t, *J* = 1.2 Hz, 1H), 6.85 (d, *J* = 8.3 Hz, 1H), 2.31 (tt, *J* = 11.7, 3.5 Hz, 1H), 1.78 (ddt, *J* = 23.2, 12.4, 3.3 Hz, 4H), 1.68 – 1.63 (m, 1H), 1.42 (qd, *J* = 12.5, 3.2 Hz, 2H), 1.27 (qt, *J* = 12.4, 3.1 Hz, 2H), 1.20 (tt, *J* = 12.5, 3.2 Hz, 1H). ^13^C NMR (150 MHz, DMSO-*d*_*6*_) δ 174.0, 168.9, 143.4, 135.7, 134.1, 132.6, 123.8, 123.8, 123.8, 120.9, 120.8, 111.6, 110.0, 44.8, 29.2 (2C, s), 25.4, 25.3 (2C, s). HRMS m/z [M+H]^+^ calcd for C_19_H_21_N_4_O_2_: 337.1665, found 337.1652, LC t_R_ = 2.88 min, > 98% Purity.

##### *N*-[(3Z)-2-oxo-3-(1H-pyrrol-2-ylmethylidene)-2,3-dihydro-1H-indol-5-yl]benzamide (72)

Compound **55** was prepared by procedure A to afford a mustard solid (60 mg, 0.182 mmol, 46%). MP 254-256 °C; ^1^H NMR (400 MHz, DMSO-*d*_6_) δ 13.34 (s, 1H), 10.87 (s, 1H), 10.17 (s, 1H), 8.08 (d, *J* = 1.9 Hz, 1H), 8.03 – 7.94 (m, 2H), 7.65 (s, 1H), 7.62 – 7.50 (m, 3H), 7.43 – 7.33 (m, 2H), 6.92 (dt, *J* = 3.6, 1.7 Hz, 1H), 6.87 (d, *J* = 8.3 Hz, 1H), 6.36 (dt, *J* = 3.7, 2.3 Hz, 1H). ^13^C NMR (100 MHz, DMSO-d_6_) δ 169.3, 165.1, 135.4, 135.0, 133.1, 131.4, 129.5, 128.4 (s, 2C), 127.5 (s, 2C), 126.2, 125.8, 125.1, 120.7, 120.3, 116.9, 112.0, 111.5, 109.4. HRMS m/z [M+H]^+^ calcd for C_20_H_16_N_3_O_2_: 330.1242, found 330.1234, LC t_R_ = 4.45 min, > 98% Purity.

##### *N*-[(3Z)-3-(1H-imidazol-5-ylmethylidene)-2-oxo-2,3-dihydro-1H-indol-5-yl]benzamide (73)

Compound **73** was prepared by procedure A to afford an orange solid (78 mg, 0.236 mmol, 60%, *E*:Z ratio 43:57 - not able to distinguish E/Z isomers). MP >300 °C; Major diastereoisomer. *N-*(3-((1*H-*imidazol-5-yl)methylene)-2-oxoindolin-5-yl)benzamide. ^1^H NMR (600 MHz, DMSO-*d*_6_) δ 10.25 (bs, 1H), 10.19 (bs, 1H), 8.11 (d, *J* = 2.1 Hz, 1H), 8.01 – 7.95 (m, 3H), 7.84 (s, 1H), 7.74 (s, 1H), 7.55 – 7.49 (m, 3H), 7.45 (dd, *J* = 8.3, 2.0 Hz, 1H), 6.89 (d, *J* = 8.3 Hz, 1H). Minor diastereoisomer. *N-*(3-((1*H-*imidazol-5-yl)methylene)-2-oxoindolin-5-yl)benzamide. ^1^H NMR (600 MHz, DMSO-*d*_6_) δ 10.32 (bs, 1H), 9.31 (bs, 1H), 8.00 – 7.98 (m, 1H), 7.81 (s, 1H), 7.59 – 7.55 (m, 2H), 7.53 – 7.50 (m, 3H), 7.49 (s, 1H), 7.40 (dd, *J* = 8.2, 2.2 Hz, 1H), 6.79 (d, *J* = 8.2 Hz, 1H). HRMS m/z [M+H]^+^ calcd for C_19_H_15_N_4_O_2_: 331.1195, found 331.1181, LC t_R_ = 3.20 min, > 98% Purity.

##### *N*-[(3Z)-3-(1H-imidazol-2-ylmethylidene)-2-oxo-2,3-dihydro-1H-indol-5-yl]benzamide (74)

Compound **74** was prepared by procedure A to afford an orange solid (89 mg, 0.269 mmol, 68%). MP >300 °C; ^1^H NMR (400 MHz, DMSO-*d*_6_) δ 11.14 (s, 1H), 10.22 (s, 1H), 8.11 (d, *J* = 2.0 Hz, 1H), 7.98 (dd, *J* = 7.0, 1.9 Hz, 2H), 7.62 – 7.51 (m, 6H), 7.35 (d, *J* = 1.4 Hz, 1H), 6.93 (d, *J* = 8.4 Hz, 1H). ^13^C NMR (100 MHz, DMSO-*d*_6_) δ 169.0, 165.3, 143.4, 136.4, 134.9, 133.6, 132.6, 131.5, 128.4 (s, 2C), 127.6 (s, 2C), 124.0, 123.8, 123.7, 122.3, 120.9, 113.3, 109.9. HRMS m/z [M+H]^+^ calcd for C_19_H_15_N_4_O_2_: 331.1195, found 331.1186, LC t_R_ = 3.23 min, > 98% Purity.

##### *N*-[(3Z)-2-oxo-3-(1H-pyrrol-2-ylmethylidene)-2,3-dihydro-1H-indol-5-yl]furan-2-carboxamide (75)

Compound **75** was prepared by procedure A to afford an orange solid (74 mg, 0.232 mmol, 56%). MP 226-228 °C; ^1^H NMR (600 MHz, DMSO-*d*_6_) δ 13.38 (s, 1H), 10.94 (s, 1H), 10.13 (s, 1H), 8.01 (d, *J* = 2.0 Hz, 1H), 7.94 (d, *J* = 1.7 Hz, 1H), 7.64 (s, 1H), 7.39 (dd, *J* = 8.3, 2.0 Hz, 1H), 7.37 (t, *J* = 1.9 Hz, 1H), 7.33 (d, *J* = 3.5 Hz, 1H), 6.91 (dd, *J* = 3.6, 1.4 Hz, 1H), 6.87 (d, *J* = 8.3 Hz, 1H), 6.71 (dd, *J* = 3.5, 1.8 Hz, 1H), 6.36 (dd, *J* = 3.7, 2.4 Hz, 1H). ^13^C NMR (150 MHz, DMSO-*d*_*6*_) δ 174.0, 169.4, 156.1, 147.7, 145.5, 135.8, 132.3, 129.5, 126.2, 125.8, 125.2, 120.6, 120.3, 117.0, 114.2, 112.1, 111.5, 109.5. HRMS m/z [M+H]^+^ calcd for C_18_H_14_N_3_O_3_: 320.1035, found 320.1018, LC t_R_ = 3.90 min, > 98% Purity.

##### *N*-[(3Z)-3-(1H-imidazol-2-ylmethylidene)-2-oxo-2,3-dihydro-1H-indol-5-yl]furan-2-carboxamide (76)

Compound **76** was prepared by procedure A to afford an orange solid (72 mg, 0.225 mmol, 54%). MP >300 °C; ^1^H NMR (600 MHz, DMSO-*d*_6_) δ 14.06 (s, 1H), 11.14 (s, 1H), 10.14 (s, 1H), 8.06 (d, *J* = 2.0 Hz, 1H), 7.94 (dd, *J* = 1.7, 0.8 Hz, 1H), 7.59 (s, 1H), 7.58 – 7.52 (m, 2H), 7.35 (t, *J* = 1.1 Hz, 1H), 7.31 (dd, *J* = 3.5, 0.8 Hz, 1H), 6.91 (d, *J* = 8.3 Hz, 1H), 6.71 (dd, *J* = 3.5, 1.7 Hz, 1H). ^13^C NMR (150 MHz, DMSO-*d*_*6*_) δ ^13^C NMR (150 MHz, DMSO) δ 169.0, 156.2, 147.6, 145.6, 143.4, 136.5, 132.9, 132.6, 124.1, 123.9, 123.6, 122.3, 121.0, 114.4, 113.3, 112.1, 110.0. HRMS m/z [M+H]^+^ calcd for C_17_H_13_N_4_O_3_: 321.0988, found 321.0979, LC t_R_ = 2.35 min, > 98% Purity.

##### 4-cyano-*N*-(2-oxoindolin-5-yl)benzamide (77)

Compound **77** was prepared by procedure F to afford a colourless solid (786 mg, 2.835 mmol, 84%). MP >240 °C (decomp.); ^1^H NMR (600 MHz, DMSO-*d*_6_) δ 10.36 (s, 1H), 10.35 (s, 1H), 8.08 (d, *J* = 8.1 Hz, 2H), 8.01 (d, *J* = 8.3 Hz, 2H), 7.66 (d, *J* = 2.1 Hz, 1H), 7.52 (dd, *J* = 8.3, 2.1 Hz, 1H), 6.80 (d, *J* = 8.3 Hz, 1H), 3.50 (s, 2H). ^13^C NMR (150 MHz, DMSO-*d*_6_) δ 176.3, 163.7, 140.1, 139.1, 132.7, 132.4 (s, 2C), 128.4 (s, 2C), 126.0, 120.0, 118.4, 117.8, 113.7, 108.9, 36.1. HRMS m/z [M+H]^+^ calcd for C_16_H_12_N_3_O_2_: 278.0929, found 278.0920, LC t_R_ = 2.90 min, > 98% Purity.

##### 3-cyano-*N*-(2-oxoindolin-5-yl)benzamide (78)

Compound **78** was prepared by procedure F to afford a colourless solid (814 mg, 2.936 mmol, 87%). MP 214-216 °C; ^1^H NMR (600 MHz, DMSO-*d*_6_) δ 10.36 (s, 1H), 10.30 (s, 1H), 8.37 (t, *J* = 1.7 Hz, 1H), 8.23 (dt, *J* = 7.9, 1.4 Hz, 1H), 8.04 (dt, *J* = 7.7, 1.4 Hz, 1H), 7.74 (t, *J* = 7.8 Hz, 1H), 7.66 (d, *J* = 2.1 Hz, 1H), 7.51 (dd, *J* = 8.4, 2.1 Hz, 1H), 6.81 (d, *J* = 8.3 Hz, 1H), 3.50 (s, 2H). ^13^C NMR (150 MHz, DMSO-*d*_*6*_) δ 176.4, 163.2, 140.1, 136.0, 134.8, 132.7, 132.4, 131.2, 129.8, 126.1, 120.0, 118.4, 117.7, 111.5, 108.9, 36.1. HRMS m/z [M+H]^+^ calcd for C_16_H_12_N_3_O_2_: 278.0929, found 278.0919, LC t_R_ = 2.92 min, > 98% Purity.

##### 2-cyano-*N*-(2-oxoindolin-5-yl)benzamide (79)

Compound **79** was prepared by procedure F to afford a colourless solid (664 mg, 2.395 mmol, 71%). MP 218-220 °C; ^1^H NMR (600 MHz, DMSO-*d*_6_) δ 10.53 (s, 1H), 10.18 (s, 1H), 8.23 (s, 1H), 7.91 (ddd, *J* = 33.5, 5.4, 3.1 Hz, 1H), 7.85 (q, *J* = 7.8 Hz, 2H), 7.77 (t, *J* = 7.5 Hz, 1H), 7.26 – 7.19 (m, 2H), 6.93 (d, *J* = 8.2 Hz, 1H), 3.55 (s, 2H). ^13^C NMR (150 MHz, DMSO-*d*_*6*_) δ 176.5, 167.3, 143.7, 134.6, 133.6, 132.5, 131.6, 127.8, 127.2, 126.4, 124.7, 124.0, 123.3, 122.8, 109.1, 35.9. HRMS m/z [M+H]^+^ calcd for C_16_H_12_N_3_O_2_: 278.0929, found 278.0921, LC t_R_ = 3.08 min, > 98% Purity.

##### 4-methoxy-*N*-(2-oxoindolin-5-yl)benzamide (80)

Compound **80** was prepared by procedure F to afford a colourless solid (738 mg, 2.614 mmol, 77%). MP 125-127 °C; ^1^H NMR (600 MHz, DMSO-*d*_6_) δ 10.33 (s, 1H), 9.96 (s, 1H), 7.97 – 7.91 (m, 2H), 7.68 – 7.64 (m, 1H), 7.51 (dd, *J* = 8.4, 2.2 Hz, 1H), 7.08 – 7.01 (m, 2H), 6.79 (d, *J* = 8.3 Hz, 1H), 3.83 (s, 3H), 3.49 (s, 2H). ^13^C NMR (100 MHz, DMSO-*d*_*6*_) δ ^13^C NMR (151 MHz, DMSO) δ 176.4, 164.6, 161.8, 139.6, 133.3, 129.5 (s, 2C), 127.1, 125.9, 120.0, 117.9, 113.6 (s, 2C), 108.8, 55.4, 36.1. HRMS m/z [M+H]^+^ calcd for C_16_H_15_N_2_O_3_: 283.1083, found 283.1072, LC t_R_ = 3.01 min, > 98% Purity.

##### 3-methoxy-*N*-(2-oxoindolin-5-yl)benzamide (81)

Compound **81** was prepared by procedure F to afford a colourless solid (820 mg, 2.905 mmol, 86%). MP 125-127 °C; ^1^H NMR (600 MHz, DMSO-*d*_6_) δ 10.34 (s, 1H), 10.08 (s, 1H), 7.65 (s, 1H), 7.52 (dq, *J* = 7.8, 1.8, 1.2 Hz, 2H), 7.48 – 7.46 (m, 1H), 7.43 (t, *J* = 7.9 Hz, 1H), 7.14 (ddd, *J* = 8.2, 2.6, 1.0 Hz, 1H), 6.79 (d, *J* = 8.3 Hz, 1H), 3.83 (s, 3H), 3.50 (s, 2H). ^13^C NMR (150 MHz, DMSO-*d*_*6*_) δ 176.4, 164.9, 159.2, 139.8, 136.5, 133.1, 129.5, 125.9, 120.0, 119.8, 117.9, 117.1, 112.8, 108.8, 55.3, 36.1. HRMS m/z [M+H]^+^ calcd for C_16_H_15_N_2_O_3_: 283.1083, found 283.1073, LC t_R_ = 3.08 min, > 98% Purity.

##### 2-methoxy-*N*-(2-oxoindolin-5-yl)benzamide (82)

Compound **82** was prepared by procedure F to afford a colourless oil (676 mg, 2.395 mmol, 71%). ^1^H NMR (600 MHz, DMSO-*d*_6_) δ 10.33 (s, 1H), 9.98 (s, 1H), 7.65 (d, *J* = 2.0 Hz, 1H), 7.63 (dd, *J* = 7.5, 1.8 Hz, 1H), 7.49 (tdd, *J* = 8.8, 6.9, 2.0 Hz, 2H), 7.17 (d, *J* = 8.4 Hz, 1H), 7.06 (td, *J* = 7.5, 0.9 Hz, 1H), 6.77 (d, *J* = 8.3 Hz, 1H), 3.89 (s, 3H), 3.49 (s, 2H). ^13^C NMR (150 MHz, DMSO-*d*_*6*_) δ 176.4, 164.1, 156.5, 139.6, 133.2, 132.0, 129.7, 126.1, 125.0, 120.5, 119.2, 117.0, 112.0, 108.9, 55.9, 36.1. HRMS m/z [M+H]^+^ calcd for C_16_H_15_N_2_O_3_: 283.1083, found 283.1072, LC t_R_ = 3.21 min, > 98% Purity.

##### 4-fluoro-*N*-(2-oxoindolin-5-yl)benzamide (83)

Compound **83** was prepared by procedure F to afford a colourless solid (812 mg, 3.005 mmol, 89%). MP 150-152 °C; ^1^H NMR (600 MHz, DMSO-*d*_6_) δ 10.38 (s, 1H), 10.17 (s, 1H), 8.04 – 7.99 (m, 2H), 7.65 (d, *J* = 2.0 Hz, 1H), 7.50 (dd, *J* = 8.4, 2.1 Hz, 1H), 7.35 (t, *J* = 8.8 Hz, 2H), 6.80 (d, *J* = 8.3 Hz, 1H), 3.49 (s, 2H). ^13^C NMR (150 MHz, DMSO-*d*_6_) δ 176.4, 164.1, 164.0 (d, *J* = 248.8 Hz), 139.9, 133.1, 131.5 (d, *J* = 3.0 Hz), 130.3 (d, *J* = 9.0 Hz, 2C), 126.0, 120.1, 117.9, 115.3 (d, *J* = 21.8 Hz, 2C), 108.8, 36.1. HRMS m/z [M+H]^+^ calcd for C_15_H_12_FN_2_O_2_: 271.0883, found 271.0872, LC t_R_ = 3.09 min, > 98% Purity.

##### 3-fluoro-*N*-(2-oxoindolin-5-yl)benzamide (84)

Compound **84** was prepared by procedure F to afford a colourless solid (830 mg, 3.071 mmol, 91%). MP 89-91 °C; ^1^H NMR (600 MHz, DMSO-*d*_6_) δ 10.40 (s, 1H), 10.23 (s, 1H), 7.80 (dd, *J* = 7.8, 1.3 Hz, 1H), 7.75 (dt, *J* = 9.8, 2.1 Hz, 1H), 7.66 (d, *J* = 2.0 Hz, 1H), 7.57 (td, *J* = 8.0, 5.8 Hz, 1H), 7.52 (dd, *J* = 8.4, 2.1 Hz, 1H), 7.43 (td, *J* = 8.5, 2.6 Hz, 1H), 6.80 (d, *J* = 8.3 Hz, 1H), 3.50 (s, 2H). ^13^C NMR (150 MHz, DMSO-*d*_6_) δ 176.4, 163.7 (d, *J* = 2.3 Hz), 161.9 (d, *J* = 244.2 Hz), 140.0, 137.4 (d, *J* = 6.8 Hz), 132.9, 130.6 (d, *J* = 8.1 Hz), 126.0, 123.8 (d, *J* = 2.7 Hz), 120.1, 118.3 (d, *J* = 21.2 Hz), 117.9, 114.4 (d, *J* = 22.9 Hz), 108.8, 36.1. HRMS m/z [M+H]^+^ calcd for C_15_H_12_FN_2_O_2_: 271.0883, found 271.0873, LC t_R_ = 3.12 min, > 98% Purity.

##### 2-fluoro-*N*-(2-oxoindolin-5-yl)benzamide (85)

Compound **85** was prepared by procedure F to afford a colourless solid (438 mg, 1.621 mmol, 48%). MP 85-87 °C; ^1^H NMR (600 MHz, DMSO-*d*_6_) δ 10.36 (s, 1H), 10.26 (s, 1H), 7.74 – 7.59 (m, 2H), 7.56 (d, *J* = 9.0 Hz, 1H), 7.48 (d, *J* = 8.2 Hz, 1H), 7.33 (t, *J* = 9.8 Hz, 2H), 6.79 (t, *J* = 6.5 Hz, 1H), 3.49 (s, 2H). ^13^C NMR (150 MHz, DMSO-*d*_6_) δ 176.4, 162.4, 158.9 (d, *J* = 248.2 Hz), 139.9, 133.0, 132.3 (d, *J* = 8.2 Hz), 129.9 (d, *J* = 2.8 Hz), 126.1, 125.21 (d, *J* = 15.3 Hz), 124.6 (d, *J* = 3.1 Hz), 119.2, 117.0, 116.2 (d, *J* = 21.8 Hz), 108.9, 36.1. HRMS m/z [M+H]^+^ calcd for C_15_H_12_FN_2_O_2_: 271.0883, found 271.0866, LC t_R_ = 3.14 min, > 98% Purity.

##### *N*-(2-oxoindolin-5-yl)-4-(trifluoromethyl)benzamide (86)

Compound **86** was prepared by procedure F to afford a colourless solid (897 mg, 2.801 mmol, 83%). MP 140-142 °C; ^1^H NMR (400 MHz, DMSO-*d*_6_) δ 10.39 (s, 1H), 10.34 (s, 1H), 7.84 (dd, J = 8.0, 1.3 Hz, 1H), 7.78 (t, J = 7.5 Hz, 1H), 7.75 – 7.63 (m, 2H), 7.60 (d, J = 2.0 Hz, 1H), 7.43 (dd, J = 8.4, 2.1 Hz, 1H), 6.79 (d, J = 8.3 Hz, 1H). ^13^C NMR (100 MHz, DMSO-*d*_6_) δ 176.4, 165.2, 139.9, 136.4 (q, J = 2.2 Hz), 133.0, 131.2 (d, J = 272.0 Hz), 128.5, 126.3 (m, 1C), 126.2, 125.8 (d, J = 31.3 Hz), 125.2, 122.5, 119.1, 116.9, 108.9, 36.1. HRMS m/z [M+H]^+^ calcd for C_16_H_12_F_3_N_2_O_2_: 321.0851, found 321.0841, LC t_R_ = 3.69 min, > 98% Purity.

##### *N*-(2-oxoindolin-5-yl)-2-(trifluoromethyl)benzamide (87)

Compound **87** was prepared by procedure F to afford a colourless solid (778 mg, 2.429 mmol, 72%). MP >240 °C (decomp.); ^1^H NMR (400 MHz, DMSO-*d*_6_) δ 10.39 (s, 1H), 10.34 (s, 1H), 7.84 (dd, *J* = 7.9, 1.2 Hz, 1H), 7.81 – 7.75 (m, 1H), 7.73 – 7.64 (m, 2H), 7.60 (d, *J* = 2.0 Hz, 1H), 7.44 (dd, *J* = 8.4, 2.1 Hz, 1H), 6.79 (d, *J* = 8.3 Hz, 1H), 3.50 (s, 2H). ^13^C NMR (100 MHz, DMSO-*d*_6_) δ 176.4, 165.2, 139.9, 136.5 (q, *J* = 2.4 Hz), 133.0, 132.6, 129.9, 128.5, 126.3 (q, *J* = 4.5 Hz), 126.2, 125.8 (d, *J* = 31.3 Hz), 123.8 (d, *J* = 273.9 Hz), 119.1, 116.9, 108.9, 36.1. HRMS m/z [M+H]^+^ calcd for C_16_H_12_F_3_N_2_O_2_: 321.0851, found 321.0839, LC t_R_ = 3.23 min, > 98% Purity.

##### 4-cyano-*N*-[(3Z)-3-(1H-imidazol-5-ylmethylidene)-2-oxo-2,3-dihydro-1H-indol-5-yl]benzamide (88)

Compound **88** was prepared by procedure A to afford a red/orange solid (68 mg, 0.191 mmol, 53%, *E*:Z ratio 50:50 - not able to distinguish E/Z isomers). MP >300 °C; Diastereoisomer 1. *N-*(3-((1*H-*imidazol-5-yl)methylene)-2-oxoindolin-5-yl)-4-cyanobenzamide. ^1^H NMR (600 MHz, DMSO-*d*_6_) δ 10.50 (s, 1H), 9.48 (d, *J* = 2.2 Hz, 1H), 8.14 – 8.12 (m, 2H), 8.05 – 8.02 (m, 2H), 7.60 (s, 1H), 7.58 (s, 1H), 7.35 (bs, 1H), 6.94 (d, *J* = 8.3 Hz, 1H). Diastereoisomer 2. *N-* (3-((1*H-*imidazol-5-yl)methylene)-2-oxoindolin-5-yl)-4-cyanobenzamide. ^1^H NMR (600 MHz, DMSO-*d*_6_) δ 10.55 (s, 1H), 10.46 (s, 1H), 8.14 – 8.12 (m, 2H), 8.12 – 8.10 (m, 1H), 8.05 – 8.02 (m, 2H), 7.56 (s, 2H), 7.52 (bs, 1H), 7.41 (s, 1H), 7.39 (bs, 1H), 6.86 (d, *J* = 8.3 Hz, 1H). HRMS m/z [M+H]^+^ calcd for C_20_H_14_N_5_O_2_: 356.1148, found 356.1138, LC t_R_ = 3.13 min, > 98% Purity.

##### 3-cyano-*N*-[(3Z)-3-(1H-imidazol-5-ylmethylidene)-2-oxo-2,3-dihydro-1H-indol-5-yl]benzamide (89)

Compound **89** was prepared by procedure A to afford a red/orange solid (73 mg, 0.205 mmol, 73%, *E*:Z ratio 43:57 - not able to distinguish E/Z isomers). MP >250 °C (decomp.); Major diastereoisomer. *N-*(3-((1*H-* imidazol-5-yl)methylene)-2-oxoindolin-5-yl)-3-cyanobenzamide. ^1^H NMR (600 MHz, DMSO-*d*_6_) δ 12.94 (bs, 1H), 10.55 (s, 1H), 9.48 (d, *J* = 2.2 Hz, 1H), 8.43 (s, 1H), 8.29 – 8.26 (m, 1H), 8.07 (tt, *J* = 7.9, 1.4 Hz, 2H), 7.78 – 7.74 (m, 1H), 7.58 (m, 1H), 7.58 – 7.56 (m, 1H), 7.41 (s, 1H), 6.86 (d, *J* = 8.3 Hz, 1H). Minor diastereoisomer. *N-*(3-((1*H-*imidazol-5-yl)methylene)-2-oxoindolin-5-yl)-3-cyanobenzamide. ^1^H NMR (600 MHz, DMSO-*d*_6_) δ 14.06 (bs, 1H), 11.18 (bs, 1H), 10.45 (s, 1H), 8.43 – 8.42 (m, 1H), 8.29 – 8.26 (m, 1H), 8.10 (d, *J* = 2.1 Hz, 1H), 8.06 – 7.98 (m, 1H), 7.78 – 7.74 (m, 1H), 7.60 (s, 1H), 7.57 – 7.56 (m, 1H), 7.41 (s, 1H), 7.36 (s, 1H), 6.95 (d, *J* = 8.3 Hz, 1H). HRMS m/z [M+H]^+^ calcd for C_20_H_14_N_5_O_2_: 356.1148, found 356.1142, LC t_R_ = 3.13 min, > 98% Purity.

##### 3-cyano-*N*-[(3Z)-3-(1H-imidazol-2-ylmethylidene)-2-oxo-2,3-dihydro-1H-indol-5-yl]benzamide (90)

Compound **90** was prepared by procedure A to afford a yellow/mustard solid (69 mg, 0.194 mmol, 54%). MP >250 °C (decomp.); ^1^H NMR (600 MHz, DMSO-*d*_6_) δ 13.70 (s, 1H), 11.02 (s, 1H), 10.40 (s, 1H), 8.42 (t, *J* = 1.7 Hz, 1H), 8.27 (dt, *J* = 8.0, 1.5 Hz, 1H), 8.13 (d, *J* = 2.0 Hz, 1H), 8.07 (dt, *J* = 7.7, 1.4 Hz, 1H), 8.03 (s, 1H), 7.79 (d, *J* = 3.7 Hz, 1H), 7.76 (d, *J* = 7.8 Hz, 1H), 7.73 (s, 1H), 7.43 (dd, *J* = 8.4, 2.0 Hz, 1H), 6.91 (d, *J* = 8.3 Hz, 1H). ^13^C NMR (150 MHz, DMSO-*d*_*6*_) δ 169.1, 163.3, 139.7, 138.9, 136.4, 135.9, 134.9, 133.0, 132.4, 131.2, 129.9, 128.1, 124.4, 123.0, 121.2, 120.0, 118.4, 112.6, 111.5, 109.8. HRMS m/z [M+H]^+^ calcd for C_20_H_14_N_5_O_2_: 356.1148, found 356.1140, LC t_R_ = 3.18 min, > 98% Purity.

##### 2-cyano-*N*-[(3Z)-3-(1H-imidazol-2-ylmethylidene)-2-oxo-2,3-dihydro-1H-indol-5-yl]benzamide (91)

Compound **91** was prepared by procedure A to afford an orange solid (76 mg, 0.214 mmol, 59%, *E*:Z ratio 25:75 - not able to distinguish E/Z isomers ). MP >300 °C; (*Z*)*-N-*(3-((1*H-*imidazol-2-yl)methylene)-2-oxoindolin-5-yl)-2-cyanobenzamide. ^1^H NMR (600 MHz, DMSO-*d*_6_) δ 13.28 (bs, 1H), 11.07 (bs, 1H), 8.18 (s, 1H), 7.78 – 7.77 (m, 1H), 7.77 (s, 1H), 7.33 (s, 1H), 6.87 (s, 1H), 6.82 (d, *J* = 8.0 Hz, 1H), 6.35 – 6.34 (m, 1H). (*E*)*-N-*(3-((1*H-*imidazol-2-yl)methylene)-2-oxoindolin-5-yl)-2-cyanobenzamide. ^1^H NMR (600 MHz, DMSO-*d*_6_) δ 11.91 (bs, 1H), 10.59 (bs, 1H), 8.59 (s, 1H), 7.81 (dd, *J* = 8.0, 1.4 Hz, 1H), 7.52 (s, 1H), 7.20 (s, 1H), 7.10 (s, 1H), 6.80 (d, *J* = 8.0 Hz, 1H), 6.41 (s, 1H). HRMS m/z [M+H]^+^ calcd for C_20_H_14_N_5_O_2_: 356.1148, found 356.1135, LC t_R_ = 2.45 min, > 98% Purity.

##### *N*-[(3Z)-3-(1H-imidazol-5-ylmethylidene)-2-oxo-2,3-dihydro-1H-indol-5-yl]-4-methoxybenzamide (92)

Compound **92** was prepared by procedure A to afford a yellow solid (70 mg, 0.194 mmol, 55%). MP 275-277 °C; ^1^H NMR (600 MHz, DMSO-*d*_6_) δ 13.69 (s, 1H), 10.91 (s, 1H), 10.04 (s, 1H), 8.10 (s, 1H), 8.00 (s, 1H), 7.99 – 7.95 (m, 2H), 7.74 (s, 2H), 7.43 (dd, *J* = 8.3, 2.0 Hz, 1H), 7.09 – 7.05 (m, 2H), 6.87 (d, *J* = 8.3 Hz, 1H), 3.84 (s, 3H). ^1^H NMR (400 MHz, DMSO-*d*_*6*_) δ ^13^C NMR (150 MHz, DMSO-*d*_*6*_) δ 176.3, 168.9, 164.6, 161.8, 136.0, 133.5, 129.4 (s, 2C), 127.0, 124.4, 121.3, 120.4, 119.9, 117.8, 113.6 (s, 2C), 112.7, 109.6, 108.7, 55.4. HRMS m/z [M+H]^+^ calcd for C_20_H_17_N_4_O_3_: 361.1301, found 361.1291, LC t_R_ = 2.74 min, > 98% Purity.

##### *N*-[(3Z)-3-(1H-imidazol-2-ylmethylidene)-2-oxo-2,3-dihydro-1H-indol-5-yl]-4-methoxybenzamide (93)

Compound **93** was prepared by procedure A to afford a bright orange solid (91 mg, 0.253 mmol, 71%). MP >300 °C; ^1^H NMR (400 MHz, DMSO-*d*_6_) δ 11.14 (s, 1H), 10.06 (s, 1H), 8.10 (d, *J* = 2.0 Hz, 1H), 8.02 – 7.94 (m, 2H), 7.66 – 7.49 (m, 3H), 7.36 (s, 1H), 7.12 – 7.04 (m, 2H), 6.95 – 6.87 (m, 1H), 3.84 (s, 3H). ^13^C NMR (100 MHz, DMSO-*d*_6_) δ 169.0, 164.7, 161.8, 143.4, 136.2, 133.8, 132.6, 129.5 (2C, s), 126.9, 123.9, 123.8, 123.8, 122.3, 120.9, 113.6 (2C, s), 113.2, 109.9, 55.4. HRMS m/z [M+H]^+^ calcd for C_20_H_17_N_4_O_3_: 361.1301, found 361.1292, LC t_R_ = 3.25 min, > 98% Purity.

##### *N*-[(3Z)-3-(1H-imidazol-5-ylmethylidene)-2-oxo-2,3-dihydro-1H-indol-5-yl]-3-methoxybenzamide (94)

Compound **94** was prepared by procedure A to afford a yellow solid (98 mg, 0.272 mmol, 70%). MP 138-140 °C; ^1^H NMR (400 MHz, DMSO-*d*_6_) δ 13.71 (s, 1H), 10.99 (s, 1H), 10.18 (s, 1H), 8.12 (d, *J* = 2.0 Hz, 1H), 8.03 (s, 1H), 7.78 (s, 1H), 7.72 (s, 1H), 7.58 – 7.55 (m, 1H), 7.52 (dd, *J* = 2.7, 1.5 Hz, 1H), 7.48 – 7.44 (m, 1H), 7.44 – 7.41 (m, 1H), 7.16 (ddd, *J* = 8.2, 2.6, 1.0 Hz, 1H), 6.89 (d, *J* = 8.4 Hz, 1H), 3.85 (s, 3H). ^13^C NMR (100 MHz, DMSO-*d*_6_) δ 169.1, 164.9, 159.2, 139.7, 138.8, 136.3, 136.2, 133.3, 129.6, 128.1, 124.3, 122.9, 121.4, 120.1, 119.8, 117.2, 112.8, 109.7, 55.3. HRMS m/z [M+H]^+^ calcd for C_20_H_17_N_4_O_3_: 361.1301, found 361.1288, LC t_R_ = 2.78 min, > 98% Purity.

##### *N*-[(3Z)-3-(1H-imidazol-2-ylmethylidene)-2-oxo-2,3-dihydro-1H-indol-5-yl]-3-methoxybenzamide (95)

Compound **95** was prepared by procedure A to afford a red solid (55 mg, 0.153 mmol, 43%, *E*:Z ratio 37:63 - not able to distinguish E/Z isomers). MP >300 °C; Major diastereoisomer. *N-*(3-((1*H-*imidazol-2-yl)methylene)-2-oxoindolin-5-yl)-3-methoxybenzamide. 1H NMR (600 MHz, DMSO-*d*6) δ 12.90 (bs, 1H), 11.15 (bs, 1H), 10.53 (s, 1H), 10.21 (s, 1H), 9.44 (d, *J* = 2.2 Hz, 1H), 7.58 – 7.40 (m, 5H), 7.40 (s, 1H), 7.15 (tdd, *J* = 8.2, 2.7, 0.9 Hz, 1H), 6.85 (d, *J* = 8.2 Hz, 1H), 3.85 (s, 3H). Minor diastereoisomer. *N-*(3-((1*H-* imidazol-2-yl)methylene)-2-oxoindolin-5-yl)-3-methoxybenzamide. 1H NMR (600 MHz, DMSO-*d*6) δ 14.07 (s, 1H), 10.19 (s, 1H), 8.10 (d, *J* = 2.0 Hz, 1H), 7.60 (s, 1H), 7.58 – 7.40 (m, 5H), 7.35 (d, *J* = 1.2 Hz, 1H), 7.17 – 7.14 (m, 1H), 6.93 (d, *J* = 8.3 Hz, 1H), 3.84 (s, 3H). HRMS m/z [M+H]^+^ calcd for C_20_H_17_N_4_O_3_: 361.1301, found 361.1291, LC t_R_ = 3.28 min, > 98% Purity.

##### *N*-[(3Z)-3-(1H-imidazol-5-ylmethylidene)-2-oxo-2,3-dihydro-1H-indol-5-yl]-2-methoxybenzamide (96)

Compound **96** was prepared by procedure A to afford a mustard/yellow solid (59 mg, 0.164 mmol, 46%). MP >250 °C (decomp.); ^1^H NMR (600 MHz, DMSO-*d*_6_) δ 13.70 (s, 1H), 10.94 (d, *J* = 74.7 Hz, 1H), 10.44 (s, 1H), 8.21 – 7.99 (m, 6H), 7.76 (s, 2H), 7.52 – 7.39 (m, 1H), 6.90 (d, *J* = 8.3 Hz, 1H), 3.34 (s, 3H). ^13^C NMR (150 MHz, DMSO-*d*_*6*_) δ 206.5, 169.0, 163.8, 139.7, 139.0, 136.5, 132.9, 132.5 (s, 2C) 128.4 (s, 2C) 124.5, 123.0, 121.3, 120.1, 118.4, 113.8, 112.6, 109.7, 30.7. HRMS m/z [M+H]^+^ calcd for C_20_H_17_N_4_O_3_: 361.1301, found 361.1290, LC t_R_ = 2.96 min, > 98% Purity.

##### *N*-[(3Z)-3-(1H-imidazol-2-ylmethylidene)-2-oxo-2,3-dihydro-1H-indol-5-yl]-2-methoxybenzamide (97)

Compound **97** was prepared by procedure A to afford a bright red solid (49 mg, 0.136 mmol, 38%). MP >300 °C; ^1^H NMR (600 MHz, DMSO-*d*_6_) δ 14.08 (s, 1H), 11.13 (s, 1H), 10.08 (s, 1H), 8.07 (d, *J* = 2.0 Hz, 1H), 7.74 (dd, *J* = 7.6, 1.8 Hz, 1H), 7.68 (dd, *J* = 8.4, 2.0 Hz, 1H), 7.65 (s, 1H), 7.57 (d, *J* = 2.2 Hz, 1H), 7.53 – 7.51 (m, 1H), 7.36 (t, *J* = 1.2 Hz, 1H), 7.20 (dd, *J* = 8.5, 0.9 Hz, 1H), 7.09 (td, *J* = 7.5, 1.0 Hz, 1H), 6.91 (d, *J* = 8.4 Hz, 1H), 3.96 (s, 3H). ^13^C NMR (150 MHz, DMSO-*d*_*6*_) δ 169.0, 163.9, 156.6, 143.5, 136.2, 133.7, 132.6, 132.3, 130.0, 124.2, 124.2, 124.0, 123.7, 121.3, 120.9, 120.6, 112.3, 112.1, 110.0, 56.0. HRMS m/z [M+H]^+^ calcd for C_20_H_17_N_4_O_3_: 361.1301, found 361.1290, LC t_R_ = 2.84 min, > 98% Purity.

##### 4-fluoro-*N*-[(3Z)-3-[(1H-imidazol-5-yl)methylidene]-2-oxo-2,3-dihydro-1H-indol-5-yl]benzamide (98)

Compound **98** was prepared by procedure A to afford a yellow/orange solid (57 mg, 0.164 mmol, 44%). MP >250 °C (decomp.); ^1^H NMR (600 MHz, DMSO-*d*_6_) δ 13.72 (s, 1H), 11.00 (s, 1H), 10.22 (s, 1H), 8.12 (s, 1H), 8.07 – 8.04 (m, 2H), 8.03 (s, 1H), 7.75 (d, *J* = 32.5 Hz, 2H), 7.48 – 7.39 (m, 1H), 7.39 – 7.36 (m, 2H), 6.89 (d, *J* = 8.3 Hz, 1H). ^13^C NMR (150 MHz, DMSO-*d*_6_) δ 169.1, 164.1, 164.0 (d, *J* = 250.0 Hz), 139.7, 138.8, 136.3, 133.3, 131.4 (d, *J* = 2.8 Hz), 130.3 (d, *J* = 8.9 Hz, 2C), 128.1, 124.4, 122.9, 121.4, 120.2, 115.4 (d, *J* = 21.8 Hz, 2C), 112.8, 109.7. HRMS m/z [M+H]^+^ calcd for C_19_H_14_FN_4_O_2_: 349.1101, found 349.1092, LC t_R_ = 2.80 min, > 98% Purity.

##### 4-fluoro-*N*-[(3Z)-3-[(1H-imidazol-2-yl)methylidene]-2-oxo-2,3-dihydro-1H-indol-5-yl]benzamide (99)

Compound **99** was prepared by procedure A to afford an orange solid (87 mg, 0.250 mmol, 68%). MP >300 °C (decomp.); ^1^H NMR (400 MHz, DMSO-*d*_6_) δ 11.15 (s, 1H), 10.23 (s, 1H), 8.14 – 8.01 (m, 3H), 7.59 (s, 1H), 7.56 (dd, *J* = 8.4, 2.0 Hz, 1H), 7.45 – 7.27 (m, 3H), 6.93 (d, *J* = 8.3 Hz, 1H). ^13^C NMR (100 MHz, DMSO-*d*_6_) δ 169.0, 164.2, 164.0 (d, *J* = 249.0 Hz), 143.4, 136.4, 133.5, 132.6, 131.3 (d, *J* = 3.0 Hz), 130.3, 130.2, 124.0, 123.8 (2C, d, *J* = 19.9 Hz), 122.3, 121.0, 115.3 (2C, d, *J* = 21.8 Hz), 113.3, 109.9. HRMS m/z [M+H]^+^ calcd for C_19_H_14_FN_4_O_2_: 349.1101, found 349.1091, LC t_R_ = 2.74 min, > 98% Purity.

##### 3-fluoro-*N*-[(3Z)-3-[(1H-imidazol-5-yl)methylidene]-2-oxo-2,3-dihydro-1H-indol-5-yl]benzamide (100)

Compound **100** was prepared by procedure A to afford an orange solid (67 mg, 0.192 mmol, 52%). MP >300 °C; ^1^H NMR (600 MHz, DMSO-*d*_6_) δ 13.71 (s, 1H), 11.00 (s, 1H), 10.28 (s, 1H), 8.12 (s, 1H), 8.03 (s, 1H), 7.84 (dt, *J* = 7.8, 1.2 Hz, 1H), 7.83 – 7.75 (m, 2H), 7.73 (s, 1H), 7.60 (td, *J* = 8.0, 5.8 Hz, 1H), 7.50 – 7.38 (m, 2H), 6.90 (d, *J* = 8.3 Hz, 1H). ^13^C NMR (150 MHz, DMSO-*d*_6_) δ 169.1, 163.8 (d, *J* = 2.7 Hz), 162.0 (d, *J* = 244.4 Hz), 139.7, 138.9, 137.2 (d, *J* = 6.8 Hz), 136.4, 133.1, 130.6 (d, *J* = 8.1 Hz), 128.1, 124.4, 123.8 (d, *J* = 2.6 Hz), 123.0, 121.4, 120.1, 118.4 (d, *J* = 21.1 Hz), 114.4 (d, *J* = 22.8 Hz), 112.8, 109.8. HRMS m/z [M+H]^+^ calcd for C_19_H_14_FN_4_O_2_: 349.1101, found 349.1090, LC t_R_ = 2.80 min, > 98% Purity.

##### 3-fluoro-*N*-[(3Z)-3-[(1H-imidazol-2-yl)methylidene]-2-oxo-2,3-dihydro-1H-indol-5-yl]benzamide (101)

Compound **101** was prepared by procedure A to afford a yellow solid (75 mg, 0.215 mmol, 58%, *E*:Z ratio 35:65), MP >210 °C (decomp.); (*Z*)*-N-*(3-((1*H-*imidazol-2-yl)methylene)-2-oxoindolin-5-yl)-3-fluorobenzamide. ^1^H NMR (600 MHz, DMSO-*d*_6_) δ 10.54 (s, 1H), 10.32 (s, 1H), 9.46 (d, *J* = 2.2 Hz, 1H), 7.86 – 7.83 (m, 1H), 7.81 – 7.77 (m, 1H), 7.59 – 7.56 (m, 1H), 7.53 (dd, *J* = 8.3, 2.2 Hz, 1H), 7.47 – 7.42 (m, 2H), 7.41 (s, 1H), 6.85 (d, *J* = 8.3 Hz, 1H). (*E*)*-N-* (3-((1*H-*imidazol-2-yl)methylene)-2-oxoindolin-5-yl)-3-fluorobenzamide. ^1^H NMR (600 MHz, DMSO-*d*_6_) δ 14.06 (s, 1H), 10.29 (s, 1H), 8.10 (d, *J* = 2.0 Hz, 1H), 7.86 – 7.83 (m, 1H), 7.81 – 7.77 (m, 1H), 7.60 (s, 1H), 7.59 – 7.56 (m, 1H), 7.47 – 7.42 (m, 2H), 7.35 (s, 1H), 6.94 (d, *J* = 8.3 Hz, 1H). HRMS m/z [M+H]^+^ calcd for C_19_H_14_FN_4_O_2_: 349.1101, found 349.1088, LC t_R_ = 3.33 min, > 98% Purity.

##### 2-fluoro-*N*-[(3Z)-3-[(1H-imidazol-2-yl)methylidene]-2-oxo-2,3-dihydro-1H-indol-5-yl]benzamide (102)

Compound **102** was prepared by procedure A to afford an orange solid (48 mg, 0.138 mmol, 37%). MP >300 °C; ^1^H NMR (400 MHz, DMSO-*d*_6_) δ 11.15 (s, 1H), 10.33 (s, 1H), 8.09 (d, *J* = 2.0 Hz, 1H), 7.69 (td, *J* = 7.4, 1.8 Hz, 1H), 7.62 – 7.57 (m, 2H), 7.54 (dd, *J* = 8.4, 2.0 Hz, 1H), 7.45 – 7.22 (m, 3H), 6.92 (d, *J* = 8.3 Hz, 1H). ^13^C NMR (100 MHz, DMSO-*d*_6_) δ 169.0, 162.5, 158.9 (d, *J* = 248.8 Hz), 143.4, 136.4, 133.4, 132.5 (d, *J* = 8.4 Hz), 129.9 (d, *J* = 3.0 Hz), 125.0, 124.9, 124.6 (d, *J* = 3.4 Hz), 124.1, 123.9, 123.6, 121.5, 121.1-120.9 (m), 116.2 (d, *J* = 22.0 Hz), 112.4, 110.1. HRMS m/z [M+H]^+^ calcd for C_19_H_14_FN_4_O_2_: 349.1101, found 349.1084, LC t_R_ = 2.83 min, > 98% Purity.

##### *N*-[(3Z)-3-[(1H-imidazol-5-yl)methylidene]-2-oxo-2,3-dihydro-1H-indol-5-yl]-4-(trifluoromethyl)benzamide (103)

Compound **100** was prepared by procedure A to afford a yellow solid (58 mg, 0.146 mmol, 47%). MP >300 °C; ^1^H NMR (600 MHz, DMSO-*d*_6_) δ 13.71 (s, 1H), 11.01 (s, 1H), 10.44 (s, 1H), 8.17 (d, *J* = 8.0 Hz, 2H), 8.14 (d, *J* = 2.0 Hz, 1H), 8.03 – 8.02 (m, 1H), 7.93 (d, *J* = 8.2 Hz, 2H), 7.80 (s, 1H), 7.73 (s, 1H), 7.43 (dd, *J* = 8.3, 2.0 Hz, 1H), 6.91 (d, *J* = 8.3 Hz, 1H). ^13^C NMR (150 MHz, DMSO-*d*_6_) δ 169.1, 164.0, 139.7, 138.9, 138.7, 136.4, 133.0, 131.3 (q, *J* = 32.8, 31.9 Hz), 128.5 (s, 2C), 128.1, 125.5 (dd, *J* = 4.9, 2.2 Hz, 2C), 124.9, 124.4, 123.0 (d, *J* = 6.1 Hz), 121.4, 120.1, 112.8, 109.8. HRMS m/z [M+H]^+^ calcd for C_20_H_14_F_3_N_4_O_2_: 399.1069, found 399.1058, LC t_R_ = 3.25 min, > 98% Purity.

##### (Z)-*N*-(3-((1H-imidazol-2-yl)methylene)-2-oxoindolin-5-yl)-4- (trifluoromethyl)benzamide (104)

Compound **104** was prepared by procedure A to afford a yellow solid (51 mg, 0.128 mmol, 41%, *E*:Z ratio 34:66 - not able to distinguish E/Z isomers). MP >210 °C (decomp.); Major diastereoisomer. *N-*(3-((1*H-*imidazol-2-yl)methylene)-2-oxoindolin-5-yl)-4- (trifluoromethyl)benzamide. ^1^H NMR (600 MHz, DMSO-*d*_6_) δ 12.87 (bs, 1H), 10.55 (bs, 1H), 10.48 (s, 1H), 9.48 (d, *J* = 2.2 Hz, 1H), 8.18 – 8.17 (m, 2H), 7.92 – 7.91 (m, 2H), 7.61 – 7.56 (m, 2H), 7.41 (s, 1H), 6.86 (d, *J* = 8.3 Hz, 1H). Minor diastereoisomer. *N-*(3-((1*H-*imidazol-2-yl)methylene)-2-oxoindolin-5-yl)-4- (trifluoromethyl)benzamide. ^1^H NMR (600 MHz, DMSO-*d*_6_) δ 14.06 (bs, 1H), 11.16 (bs, 1H), 10.45 (s, 1H), 8.18 – 8.17 (m, 2H), 7.97 – 7.92 (m, 2H), 7.61 (s, 1H), 7.46 (bs, 2H), 7.35 (s, 1H), 6.95 (d, *J* = 8.3 Hz, 1H). HRMS m/z [M+H]^+^ calcd for C_20_H_14_F_3_N_4_O_2_: 399.1069, found 399.1058, LC t_R_ = 3.20 min, > 98% Purity.

##### *N*-[(3Z)-3-[(1H-imidazol-5-yl)methylidene]-2-oxo-2,3-dihydro-1H-indol-5-yl]-2-(trifluoromethyl)benzamide (105)

Compound **105** was prepared by procedure A to afford a yellow solid (46 mg, 0.116 mmol, 37%, *E*:Z ratio 38:62 - not able to distinguish E/Z isomers). MP 225-227 °C; Major Diastereoisomer. *N-*(3-((1*H-*imidazol-5-yl)methylene)-2-oxoindolin-5-yl)-2- (trifluoromethyl)benzamide. ^1^H NMR (600 MHz, DMSO-*d*_6_) δ 10.47 (bs, 1H), 10.42 (bs, 1H), 9.29 (d, *J* = 2.1 Hz, 1H), 7.88 (s, 2H), 7.86 – 7.68 (m, 4H), 7.60 (dd, *J* = 8.3, 2.2 Hz, 1H), 7.50 (s, 1H), 6.82 (d, *J* = 8.3 Hz, 1H). Minor Diastereoisomer. *N-*(3-((1*H-*imidazol-5-yl)methylene)-2-oxoindolin-5-yl)-2- (trifluoromethyl)benzamide. ^1^H NMR (600 MHz, DMSO-*d*_6_) δ 10.53 (bs, 1H), 8.09 (d, *J* = 2.0 Hz, 1H), 8.00 (s, 1H), 7.95 (s, 1H), 7.86 – 7.68 (m, 5H), 7.34 (dd, *J* = 8.3, 2.0 Hz, 1H), 6.89 (d, *J* = 8.3 Hz, 1H). HRMS m/z [M+H]^+^ calcd for C_20_H_14_F_3_N_4_O_2_: 399.1069, found 399.1058, LC t_R_ = 2.89 min, > 98% Purity.

##### *N*-[(3Z)-3-[(1H-imidazol-2-yl)methylidene]-2-oxo-2,3-dihydro-1H-indol-5-yl]-2-(trifluoromethyl)benzamide (106)

Compound **106** was prepared by procedure A to afford a yellow solid (56 mg, 0.141 mmol, 45%, *E*:Z ratio 34:66 - not able to distinguish E/Z isomers). MP >300 °C; Major diastereoisomer. *N-*(3-((1*H-*imidazol-2-yl)methylene)-2-oxoindolin-5-yl)-2- (trifluoromethyl)benzamide. ^1^H NMR (600 MHz, DMSO-*d*_6_) δ 12.93 (bs, 1H), 10.54 (bs, 1H), 10.50 (bs, 1H), 9.45 (d, *J* = 2.1 Hz, 1H), 7.87 – 7.83 (m, 1H), 7.81 – 7.77 (m, 1H), 7.73 – 7.68 (m, 3H), 7.50 (s, 1H), 7.41 (s, 1H), 7.36 (s, 1H), 6.85 (d, *J* = 8.3 Hz, 1H). Minor diastereoisomer. *N-*(3-((1*H-*imidazol-2-yl)methylene)-2-oxoindolin-5-yl)-2- (trifluoromethyl)benzamide. ^1^H NMR (600 MHz, DMSO-*d*_6_) δ 14.07 (bs, 1H), 11.17 (bs, 1H), 10.51 (bs, 1H), 8.04 (d, *J* = 2.0 Hz, 1H), 7.81 – 7.77 (m, 1H), 7.70 – 7.68 (m, 3H), 7.58 – 7.57 (m, 2H), 7.52 (d, *J* = 2.0 Hz, 1H), 7.37 – 7.34 (m, 1H), 6.93 (d, *J* = 8.3 Hz, 1H). HRMS m/z [M+H]^+^ calcd for C_20_H_14_F_3_N_4_O_2_: 399.1069, found 399.1057, LC t_R_ = 2.86 min, > 98% Purity.

##### *N*-(2-oxoindolin-5-yl)thiophene-2-carboxamide (107)

Compound **107** was prepared by procedure F to afford a red solid (514 mg, 1.991 mmol, 59%). MP 105-107 °C; ^1^H NMR (600 MHz, DMSO-*d*_6_) δ 10.34 (s, 1H), 10.10 (s, 1H), 7.97 (dd, *J* = 3.7, 1.1 Hz, 1H), 7.82 (dd, *J* = 5.0, 1.1 Hz, 1H), 7.63 – 7.57 (m, 1H), 7.46 (dd, *J* = 8.4, 2.1 Hz, 1H), 7.21 (dd, *J* = 5.0, 3.7 Hz, 1H), 6.79 (d, *J* = 8.3 Hz, 1H), 3.50 (s, 2H). ^13^C NMR (150 MHz, DMSO-*d*_*6*_) δ 176.3, 159.6, 140.3, 139.9, 132.6, 131.5, 128.7, 128.0, 126.1, 120.1, 117.9, 108.9, 36.1. HRMS m/z [M+H]^+^ calcd for C_13_H_11_N_2_O_2_S: 259.0541, found 259.0532, LC t_R_ = 2.84 min, > 98% Purity.

##### 3-methoxy-*N*-(2-oxoindolin-5-yl)thiophene-2-carboxamide (108)

Compound **108** was prepared by procedure F to afford a beige solid (662 mg, 2.296 mmol, 68%). MP 75-77 °C; ^1^H NMR (600 MHz, DMSO-*d*_6_) δ 10.37 (s, 1H), 9.14 (s, 1H), 7.80 (d, *J* = 5.5 Hz, 1H), 7.59 – 7.55 (m, 1H), 7.43 (dd, *J* = 8.3, 2.2 Hz, 1H), 7.18 (d, *J* = 5.6 Hz, 1H), 6.77 (d, *J* = 8.3 Hz, 1H), 4.06 (s, 3H), 3.48 (s, 2H). ^13^C NMR (150 MHz, DMSO-*d*_*6*_) δ 176.3, 159.1, 156.7, 139.8, 132.2, 130.4, 126.2, 119.6, 117.4, 117.1, 115.6, 108.9, 59.5, 36.0. HRMS m/z [M+H]^+^ calcd for C_14_H_13_N_2_O_3_S: 289.0647, found 289.0637, LC t_R_ = 3.05 min, > 98% Purity.

##### 3-ethoxy-*N*-(2-oxoindolin-5-yl)thiophene-2-carboxamide (109)

Compound **109** was prepared by procedure F to afford a beige solid (796 mg, 2.633 mmol, 78%). MP 98-100 °C; ^1^H NMR (600 MHz, DMSO-*d*_6_) δ 10.33 (s, 1H), 9.20 (s, 1H), 7.78 (d, *J* = 5.5 Hz, 1H), 7.56 – 7.51 (m, 1H), 7.39 (dd, *J* = 8.4, 2.1 Hz, 1H), 7.16 (d, *J* = 5.5 Hz, 1H), 6.78 (d, *J* = 8.3 Hz, 1H), 4.33 (q, *J* = 7.0 Hz, 2H), 3.49 (s, 2H), 1.45 (t, *J* = 7.0 Hz, 3H). ^13^C NMR (150 MHz, DMSO-*d*_*6*_) δ 176.3, 159.0, 155.8, 139.8, 132.3, 130.4, 126.4, 119.0, 117.5, 116.9, 115.9, 109.1, 68.0, 36.1, 14.8. HRMS m/z [M+H]^+^ calcd for C_15_H_15_N_2_O_3_S: 303.0803, found 303.0795, LC t_R_ = 3.56 min, > 98% Purity.

##### 3-fluoro-*N*-(2-oxoindolin-5-yl)thiophene-2-carboxamide (110)

Compound **110** was prepared by procedure F to afford a beige solid (532 mg, 1.926 mmol, 57%). MP 101-103 °C; HRMS m/z [M+H]^+^ calcd for C_13_H_10_FN_2_O_2_S: 277.0447, found 277.0438, LC t_R_ = 3.35 min, > 98% Purity.

##### *N*-(2-oxoindolin-5-yl)thieno[3,2-*b*]thiophene-2-carboxamide (111)

Compound **111** was prepared by procedure F to afford a beige solid (366 mg, 1.164 mmol, 69%). MP 125-127 °C; HRMS m/z [M+H]^+^ calcd for C_15_H_11_N_2_O_2_S_2_: 315.0262, found 315.0251, LC t_R_ = 3.45 min, > 98% Purity.

##### 2-fluoro-*N*-(2-oxoindolin-5-yl)-5-(trifluoromethyl)benzamide (112)

Compound **112** was prepared by procedure F to afford a beige solid (651 mg, 1.925 mmol, 57%). MP 96-97 °C; ^1^H NMR (600 MHz, DMSO-*d*_6_) δ 10.46 (s, 1H), 10.38 (s, 1H), 8.03 (dd, *J* = 6.2, 2.5 Hz, 1H), 7.97 (dt, *J* = 7.7, 3.2 Hz, 1H), 7.65 – 7.54 (m, 2H), 7.46 (dd, *J* = 8.3, 2.1 Hz, 1H), 6.81 (d, *J* = 8.3 Hz, 1H), 3.50 (s, 2H). ^13^C NMR (150 MHz, DMSO-*d*_6_) δ 176.4, 162.0-160.0 (m, 1C), 140.1, 132.6, 129.6 (dd, *J* = 9.6, 3.9 Hz), 127.3 (p, *J* = 3.9 Hz), 126.2, 126.1 (d, *J* = 17.0 Hz), 125.4 (qd, *J* = 32.8, 3.2 Hz), 124.5, 122.7, 119.4, 117.7 (d, *J* = 23.5 Hz), 117.2, 109.0, 36.1. HRMS m/z [M+H]^+^ calcd for C_16_H_11_F_4_N_2_O_2_: 339.0757, found 339.0740, LC t_R_ = 3.80 min, > 98% Purity.

##### *N*-(2-oxoindolin-5-yl)benzo[*d*][1,3]dioxole-4-carboxamide (113)

Compound **113** was prepared by procedure F to afford a beige solid (750 mg, 2.531 mmol, 75%). MP 105-107 °C; ^1^H NMR (600 MHz, DMSO-*d*_6_) δ 10.38 (s, 1H), 9.70 (s, 1H), 7.60 (d, *J* = 2.0 Hz, 1H), 7.47 (dd, *J* = 8.4, 2.0 Hz, 1H), 7.22 (d, *J* = 8.1 Hz, 1H), 7.10 (d, *J* = 7.7 Hz, 1H), 6.96 (t, *J* = 7.9 Hz, 1H), 6.78 (d, *J* = 8.3 Hz, 1H), 6.15 (s, 2H), 3.49 (s, 2H). ^13^C NMR (150 MHz, DMSO-*d*_*6*_) δ 176.3, 161.8, 147.8, 145.2, 139.9, 132.7, 126.1, 121.7, 121.1, 119.4, 117.7, 117.2, 110.9, 108.90, 101.7, 36.1. HRMS m/z [M+H]^+^ calcd for C_16_H_13_N_2_O_4_: 297.0875, found 297.0865, LC t_R_ = 3.18 min, > 98% Purity.

##### *N*-(2-oxoindolin-5-yl)thiazole-2-carboxamide (114)

Compound **114** was prepared by procedure F to afford a beige solid (735 mg, 2.835 mmol, 84%). MP 108-110 °C; ^1^H NMR (600 MHz, DMSO-*d*_6_) δ 10.71 (s, 1H), 10.39 (s, 1H), 8.38 (s, 1H), 7.68 (d, *J* = 2.0 Hz, 1H), 7.57 (dd, *J* = 8.4, 2.1 Hz, 1H), 7.53 (s, 1H), 6.79 (d, *J* = 8.4 Hz, 1H), 3.50 (s, 2H). ^13^C NMR (150 MHz, DMSO-*d*_6_) δ 176.4, 155.1, 152.9, 142.5, 140.4, 131.8, 128.3, 126.1, 120.3, 117.8, 108.9, 36.0. HRMS m/z [M+H]^+^ calcd for C_12_H_10_N_3_O_2_S: 260.0494, found 260.0484, LC t_R_ = 2.72 min, > 98% Purity.

##### *N*-(2-oxoindolin-5-yl)oxazole-2-carboxamide (115)

Compound **115** was prepared by procedure F to afford a beige solid (345 mg, 1.418 mmol, 70%). MP 87-89 °C; ^1^H NMR (600 MHz, DMSO-*d*_6_) δ 10.65 (s, 1H), 10.39 (s, 1H), 8.13 – 8.07 (m, 2H), 7.72 (d, *J* = 2.0 Hz, 1H), 7.63 (dd, *J* = 8.4, 2.1 Hz, 1H), 6.80 (d, *J* = 8.4 Hz, 1H), 3.49 (s, 2H). ^13^C NMR (150 MHz, DMSO-*d*_*6*_) δ 176.8, 164.6, 157.9, 144.4, 140.7, 132.4, 126.7, 126.5, 120.7, 118.3, 109.3, 36.5. HRMS m/z [M+H]^+^ calcd for C_12_H_10_N_3_O_3_: 244.0722, found 244.0709, LC t_R_ = 2.25 min, > 98% Purity.

##### *N*-(2-oxoindolin-5-yl)isothiazole-5-carboxamide (116)

Compound **116** was prepared by procedure A to afford a beige solid (328 mg, 1.265 mmol, 75%). MP 178-180 °C; ^1^H NMR (400 MHz, DMSO-*d*_6_) δ 10.47 (s, 1H), 10.38 (s, 1H), 8.71 (d, *J* = 1.8 Hz, 1H), 8.09 (d, *J* = 1.8 Hz, 1H), 7.61 (d, *J* = 2.0 Hz, 1H), 7.47 (dd, *J* = 8.4, 2.1 Hz, 1H), 6.82 (d, *J* = 8.3 Hz, 1H), 3.51 (s, 2H). ^13^C NMR (100 MHz, DMSO-*d*_6_) δ 176.33, 163.32, 159.18, 157.40, 140.49, 131.77, 126.22, 123.87, 120.32, 117.97, 108.97, 36.03. HRMS m/z [M+H]^+^ calcd for C_12_H_10_N_3_O_2_S: 260.0494, found 260.0485, LC t_R_ = 2.57 min, > 98% Purity.

##### *N*-[(3Z)-3-[(1H-imidazol-5-yl)methylidene]-2-oxo-2,3-dihydro-1H-indol-5-yl]thiophene-2-carboxamide (117)

Compound **117** was prepared by procedure A to afford an orange solid (89 mg, 0.265 mmol, 68%). MP>300 °C; ^1^H NMR (600 MHz, DMSO-*d*_6_) δ 13.71 (s, 1H), 11.00 (s, 1H), 10.22 (s, 1H), 8.06 (d, *J* = 2.0 Hz, 1H), 8.03 (s, 1H), 8.01 – 8.00 (m, 1H), 7.84 (dd, *J* = 5.0, 1.1 Hz, 1H), 7.80 (s, 1H), 7.71 (s, 1H), 7.39 (dd, *J* = 8.3, 2.0 Hz, 1H), 7.23 (dd, *J* = 5.0, 3.7 Hz, 1H), 6.90 (d, *J* = 8.3 Hz, 1H). ^13^C NMR (150 MHz, DMSO -*d*_6_) δ 169.1, 159.7, 140.2, 139.7, 138.8, 136.3, 132.8, 131.6, 128.8, 128.1 (s, 2C), 124.5, 123.0, 121.4, 120.1, 112.9, 109.8. HRMS m/z [M+H]^+^ calcd for C_17_H_13_N_4_O_2_S: 337.0759, found 327.0721, LC t_R_ = 2.78 min, > 98% Purity.

##### *N*-[(3Z)-3-[(1H-imidazol-2-yl)methylidene]-2-oxo-2,3-dihydro-1H-indol-5-yl]thiophene-2-carboxamide (118)

Compound **118** was prepared by procedure A to afford an orange solid (92 mg, 0.274 mmol, 71%, *E*:Z ratio 34:66 - not able to distinguish E/Z isomers). MP >300 °C; Major diastereoisomer. *N-*(3-((1*H-*imidazol-2-yl)methylene)-2-oxoindolin-5-yl)thiophene-2-carboxamide. ^1^H NMR (600 MHz, DMSO-*d*_6_) δ 12.94 (bs, 1H), 10.55 (bs, 1H), 10.26 (bs, 1H), 9.42 (d, *J* = 2.2 Hz, 1H), 8.05 (d, *J* = 3.9 Hz, 1H), 7.83 (dd, *J* = 5.0, 1.1 Hz, 1H), 7.49 (dd, *J* = 8.3, 2.2 Hz, 1H), 7.50 – 7.44 (bs, 2H), 7.41 (s, 1H), 7.22 (dd, *J* = 5.0, 3.7 Hz, 1H), 6.85 (d, *J* = 8.3 Hz, 1H). Minor diastereoisomer. *N-*(3-((1*H-*imidazol-2-yl)methylene)-2-oxoindolin-5-yl)thiophene-2-carboxamide. ^1^H NMR (600 MHz, DMSO-*d*_6_) δ 14.07 (bs, 1H), 11.16 (bs, 1H), 10.22 (bs, 1H), 8.06 (d, *J* = 2.0 Hz, 1H), 8.01 (dd, *J* = 3.8, 1.2 Hz, 1H), 7.85 (dd, *J* = 5.0, 1.1 Hz, 1H), 7.62 (s, 1H), 7.57 (d, *J* = 2.1 Hz, 1H), 7.52 (dd, *J* = 8.4, 2.1 Hz, 1H), 7.36 (bt, *J* = 1.2 Hz, 1H), 7.23 (dd, *J* = 5.0, 3.7 Hz, 1H), 6.93 (d, *J* = 8.3 Hz, 1H). HRMS m/z [M+H]^+^ calcd for C_17_H_13_N_4_O_2_S: 337.0759, found 337.0753, LC t_R_ = 3.07 min, > 98% Purity.

##### (Z)-*N*-(3-((1H-imidazol-5-yl)methylene)-2-oxoindolin-5-yl)-3-methoxythiophene-2-carboxamide (119)

Compound **119** was prepared by procedure A to afford an orange solid (45 mg, 0.123 mmol, 35%). MP 268-270 °C; ^1^H NMR (400 MHz, DMSO-*d*_6_) δ 13.72 (s, 1H), 10.99 (s, 1H), 9.20 (s, 1H), 8.03 (s, 1H), 7.96 (d, J = 2.0 Hz, 1H), 7.84 – 7.80 (m, 2H), 7.69 (s, 1H), 7.50 – 7.47 (m, 1H), 7.20 (d, J = 5.6 Hz, 1H), 6.87 (d, J = 8.3 Hz, 1H), 4.09 (s, 3H). ^13^C NMR (100 MHz, DMSO-*d*_6_) δ 169.1, 159.2, 156.8, 139.7, 138.7, 136.2, 132.5, 130.5, 128.1, 124.6, 123.1, 120.7, 120.0, 117.1, 115.5, 112.3, 109.7, 59.6. HRMS m/z [M+H]^+^ calcd for C_18_H_15_N_4_O_3_S: 367.0865, found 367.0854, LC t_R_ = 2.78 min, > 98% Purity.

##### 3-ethoxy-*N*-[(3Z)-3-[(1H-imidazol-5-yl)methylidene]-2-oxo-2,3-dihydro-1H-indol-5-yl]thiophene-2-carboxamide (120)

Compound **120** was prepared by procedure A to afford an orange solid (52 mg, 0.137 mmol, 41%, *E*:Z ratio 28:72 - not able to distinguish E/Z isomers). MP Decomp. >250 °C; Major diastereoisomer. *N-*(3-((1*H-*imidazol-5-yl)methylene)-2-oxoindolin-5-yl)-3-ethoxythiophene-2-carboxamide. ^1^H NMR (600 MHz, DMSO-*d*_6_) δ 12.74 (bs, 1H), 10.39 (bs, 1H), 9.32 (s, 1H), 9.23 (bs, 1H), 7.96 – 7.95 (m, 2H), 7.87 (dd, *J* = 8.3, 2.2 Hz, 1H), 7.81 (d, *J* = 5.5 Hz, 1H), 7.51 (s, 1H), 7.21 (d, *J* = 5.5 Hz, 1H), 6.81 (d, *J* = 8.3 Hz, 1H), 4.41 – 4.37 (m, 2H), 1.59 (t, *J* = 6.9 Hz, 3H). Minor diastereoisomer. *N-*(3-((1*H-*imidazol-5-yl)methylene)-2-oxoindolin-5-yl)-3-ethoxythiophene-2-carboxamide. ^1^H NMR (600 MHz, DMSO-*d*_6_) δ 13.73 (bs, 1H), 10.99 (bs, 1H), 9.25 (s, 1H), 8.02 (d, *J* = 14.3 Hz, 2H), 7.83 (s, 1H), 7.81 – 7.79 (m, 1H), 7.71 (s, 1H), 7.37 (dd, *J* = 8.4, 2.1 Hz, 1H), 7.19 (d, *J* = 5.5 Hz, 1H), 6.89 (d, *J* = 8.3 Hz, 1H), 4.39 – 4.35 (m, 2H), 1.47 (t, *J* = 7.0 Hz, 3H). HRMS m/z [M+H]^+^ calcd for C_19_H_17_N_4_O_3_S: 381.1021, found 381.1007, LC t_R_ = 3.03 min, > 98% Purity.

##### 3-fluoro-*N*-[(3Z)-3-[(1H-imidazol-5-yl)methylidene]-2-oxo-2,3-dihydro-1H-indol-5-yl]thiophene-2-carboxamide (121)

Compound **121** was prepared by procedure A to afford an orange solid (37 mg, 0.104 mmol, 29%). MP >300 °C; ^1^H NMR (400 MHz, DMSO-*d*_6_) δ 13.69 (s, 1H), 11.00 (s, 1H), 9.83 – 9.72 (m, 1H), 8.03 (s, 1H), 7.98 (d, *J* = 2.0 Hz, 1H), 7.86 (dd, *J* = 5.5, 4.0 Hz, 1H), 7.81 (s, 1H), 7.70 (s, 1H), 7.38 (dd, *J* = 8.3, 2.0 Hz, 1H), 7.16 (d, *J* = 5.5 Hz, 1H), 6.88 (d, *J* = 8.3 Hz, 1H). ^13^C NMR (100 MHz, DMSO-*d*_6_) δ 169.10, 157.67, 156.86, 154.22, 139.70, 138.82, 136.49, 132.43, 129.77 (d, *J* = 9.9 Hz), 128.08, 124.46, 123.10, 121.51, 119.97, 118.30 (d, *J* = 26.4 Hz), 113.03, 109.72. HRMS m/z [M+H]^+^ calcd for C_17_H_12_FN_4_O_2_S: 355.0665, found 355.0652, LC t_R_ = 2.68 min, > 98% Purity.

##### (Z)-*N*-(3-((1H-imidazol-5-yl)methylene)-2-oxoindolin-5-yl)thieno[3,2-b]thiophene-2-carboxamide (122)

Compound **122** was prepared by procedure A to afford a light orange solid (45 mg, 0.110 mmol, 36%). MP >250 °C (decomp.); ^1^H NMR (600 MHz, DMSO-*d*_6_) δ 13.68 (s, 1H), 10.95 (s, 1H), 10.34 (s, 1H), 8.34 (s, 1H), 8.09 (d, *J* = 2.0 Hz, 1H), 8.01 (s, 1H), 7.89 (d, *J* = 5.2 Hz, 1H), 7.78 (s, 1H), 7.54 (d, *J* = 5.2 Hz, 1H), 7.40 (dd, *J* = 8.2, 2.0 Hz, 1H), 6.90 (d, *J* = 8.3 Hz, 1H). ^13^C NMR (150 MHz, DMSO-*d*_*6*_) δ 169.0, 160.1, 142.0, 141.6, 139.4, 138.5, 136.3, 132.7, 131.8, 124.5, 121.4 (s, 2C), 121.2, 120.6, 120.4 (s, 2C), 120.3, 112.7, 109.7. HRMS m/z [M+H]^+^ calcd for C_19_H_13_N_4_O_2_S_2_: 393.0480, found 393.0468, LC t_R_ = 3.08 min, > 98% Purity.

##### 2-fluoro-*N*-[(3Z)-3-[(1H-imidazol-5-yl)methylidene]-2-oxo-2,3-dihydro-1H-indol-5-yl]-5-(trifluoromethyl)benzamide (123)

Compound **123** was prepared by procedure A to afford an orange solid (21 mg, 0.062 mmol, 21%). MP 202-204 °C; ^1^H NMR (600 MHz, DMSO-*d*_6_) δ 13.73 (s, 1H), 10.99 (s, 1H), 10.75 (s, 1H), 8.22 (s, 1H), 8.03 (s, 1H), 7.85 (d, *J* = 2.3 Hz, 1H), 7.79 – 7.75 (m, 2H), 7.35 (dd, *J* = 9.4, 5.8 Hz, 2H), 6.91 (d, *J* = 8.4 Hz, 1H). ^13^C NMR (150 MHz, DMSO) δ 169.1, 164.6, 154.3, 139.7, 139.0, 136.1, 133.3, 128.9, 128.1, 126.7, 125.3, 124.6, 123.5, 123.0, 121.7, 121.4, 120.1, 119.8, 111.4, 110.1. HRMS m/z [M+H]^+^ calcd for C_20_H_13_F_4_N_4_O_2_: 417.0975, found 417.0962, LC t_R_ = 3.22 min, > 98% Purity.

##### (Z)-*N*-(3-((1H-imidazol-5-yl)methylene)-2-oxoindolin-5-yl)benzo[d][1,3]dioxole-4-carboxamide (124)

Compound **124** was prepared by procedure A to afford a brown/red solid (47 mg, 0.126 mmol, 37%). MP 189-191 °C; ^1^H NMR (600 MHz, DMSO-*d*_6_) δ 13.71 (s, 1H), 10.96 (s, 1H), 9.72 (s, 1H), 8.30 (s, 1H), 8.06 – 8.03 (m, 1H), 8.00 (d, *J* = 4.2 Hz, 1H), 7.85 – 7.66 (m, 2H), 7.45 (dt, *J* = 8.3, 3.1 Hz, 1H), 7.27 (dd, *J* = 8.1, 1.1 Hz, 1H), 7.12 (dd, *J* = 7.7, 1.1 Hz, 1H), 6.98 (t, *J* = 7.9 Hz, 1H), 6.88 (d, *J* = 8.3 Hz, 1H), 6.18 (s, 2H). ^13^C NMR (150 MHz, DMSO-*d*_*6*_) δ 169.0, 161.9, 147.8, 145.3, 139.5, 136.3, 134.6, 132.9, 128.5, 124.6, 121.7, 121.1, 120.6, 120.3, 117.5, 112.0, 111.0, 109.7, 101.7, 43.8. HRMS m/z [M+H]^+^ calcd for C_20_H_15_N_4_O_4_: 375.1093, found 375.1081, LC t_R_ = 2.85 min, > 98% Purity.

##### *N*-[(3Z)-3-[(1H-imidazol-5-yl)methylidene]-2-oxo-2,3-dihydro-1H-indol-5-yl]-1,3-thiazole-2-carboxamide (125)

Compound **125** was prepared by procedure A to afford a mustard green solid (60 mg, 0.178 mmol, 46%). MP 271-273 °C; ^1^H NMR (600 MHz, DMSO-*d*_6_) δ 13.70 (s, 1H), 11.01 (s, 1H), 10.70 (s, 1H), 8.17 (s, 1H), 8.13 – 8.10 (m, 2H), 8.03 (s, 1H), 7.74 (d, *J* = 24.9 Hz, 2H), 7.55 (d, *J* = 8.0 Hz, 1H), 6.89 (d, *J* = 8.3 Hz, 1H). ^13^C NMR (150 MHz, DMSO-*d*_6_) δ 169.1, 164.1, 157.6, 157.6, 144.0, 139.7, 138.9, 136.6, 132.2, 126.4, 124.5, 123.0, 121.5, 120.1, 112.7, 109.8. HRMS m/z [M+H]^+^ calcd for C_16_H_12_N_5_O_2_S: 338.0712, found 338.0700, LC t_R_ = 2.52 min, > 98% Purity.

##### *N*-[(3Z)-3-[(1H-imidazol-5-yl)methylidene]-2-oxo-2,3-dihydro-1H-indol-5-yl]-1,3-oxazole-2-carboxamide (126)

Compound **126** was prepared by procedure A to afford a dark orange solid (57 mg, 0.177 mmol, 43%). MP >300 °C; ^1^H NMR (400 MHz, DMSO-*d*_6_) δ 13.68 (s, 1H), 10.99 (s, 1H), 10.73 (s, 1H), 8.38 (dd, *J* = 6.0, 0.8 Hz, 1H), 8.15 – 7.96 (m, 2H), 7.73 (d, *J* = 14.6 Hz, 2H), 7.52 (dd, *J* = 4.0, 0.8 Hz, 1H), 7.47 (d, *J* = 7.7 Hz, 1H), 6.83 (dd, *J* = 29.0, 8.3 Hz, 1H). ^13^C NMR (100 MHz, DMSO-*d*_6_) δ 169.1, 155.1, 153.0, 142.6, 139.7, 138.9, 136.7, 132.0, 128.4, 128.0, 124.4, 123.1, 121.5, 119.9, 112.7, 109.8. HRMS m/z [M+H]^+^ calcd for C_16_H_12_N_5_O_3_: 322.0940, found 322.0930, LC t_R_ = 2.21 min, > 98% Purity.

##### *N*-[(3Z/E)-3-[(1H-imidazol-5-yl)methylidene]-2-oxo-2,3-dihydro-1H-indol-5-yl]-1,2-thiazole-5-carboxamide (127)

Compound **127** was prepared by procedure A to afford an orange solid (48 mg, 0.142 mmol, 37%, *E*:Z ratio 30:70). MP >300 °C; (*Z*)*-N-*(3-((1*H-*imidazol-5-yl)methylene)-2-oxoindolin-5-yl)isothiazole-5-carboxamide. ^1^H NMR (600 MHz, DMSO-*d*_6_) δ 12.74 (s, 1H), 10.60 (s, 1H), 10.46 (s, 1H), 9.41 (d, *J* = 2.1 Hz, 1H), 8.72 (dd, *J* = 3.5, 1.7 Hz, 1H), 8.16 (d, *J* = 1.8 Hz, 1H), 8.03 (s, 1H), 7.95 (s, 1H), 7.54 (s, 1H), 7.50 – 7.44 (m, 1H), 6.86 (d, *J* = 8.2 Hz, 1H). (*E*)*-N-*(3-((1*H-*imidazol-5-yl)methylene)-2-oxoindolin-5-yl)isothiazole-5-carboxamide. ^1^H NMR (600 MHz, DMSO-d_6_) δ 13.71 (s, 1H), 11.05 (s, 1H), 10.59 (s, 1H), 8.12 (s, 0H), 8.07 (s, 1H), 8.04 (s, 1H), 7.82 (s, 1H), 7.73 (s, 1H), 7.42 (d, *J* = 7.5 Hz, 1H), 6.93 (d, *J* = 8.3 Hz, 1H).HRMS m/z [M+H]^+^ calcd for C_16_H_12_N_5_O_2_S: 338.0712, found 338.0699, LC t_R_ = 2.41 min, > 98% Purity.

##### *N*-[(3Z)-3-[(4-methyl-1H-imidazol-5-yl)methylidene]-2-oxo-2,3-dihydro-1H-indol-5-yl]benzamide (128)

Compound **128** was prepared by procedure A to afford a light orange solid (59 mg, 0.171 mmol, 43%). MP >270 °C (decomp.); ^1^H NMR (400 MHz, DMSO-*d*_6_) δ 13.91 (s, 1H), 11.02 (s, 1H), 10.23 (s, 1H), 8.15 (d, *J* = 2.0 Hz, 1H), 8.07 – 8.02 (m, 2H), 7.99 (s, 1H), 7.71 (d, *J* = 0.7 Hz, 1H), 7.67 – 7.62 (m, 1H), 7.62 – 7.56 (m, 2H), 7.51 (dd, *J* = 8.4, 2.0 Hz, 1H), 6.94 (d, *J* = 8.3 Hz, 1H), 2.50 (s, 3H). ^13^C NMR (100 MHz, DMSO-*d*_6_) δ 169.5, 165.2, 147.6, 138.2, 135.9, 134.8, 133.1, 131.5, 128.4 (s, 2C), 127.5 (s, 2C), 124.8, 124.4, 122.3, 121.4, 118.1, 113.4, 109.5, 13.1. HRMS m/z [M+H]^+^ calcd for C_20_H_17_N_4_O_2_: 345.1352, found 345.1342, LC t_R_ = 2.76 min, > 98% Purity.

##### *N*-[(3Z)-2-oxo-3-(1H-pyrazol-5-ylmethylidene)-2,3-dihydro-1H-indol-5-yl]benzamide (129)

Compound **129** was prepared by procedure A to afford a dark red/purple solid (87 mg, 0.263 mmol, 66%). MP 218-220 °C; ^1^H NMR (600 MHz, DMSO-*d*_6_) δ 13.54 (s, 1H), 10.53 (s, 1H), 10.21 (s, 1H), 7.99 – 7.97 (m, 2H), 7.91 (d, *J* = 2.2 Hz, 1H), 7.72 – 7.44 (m, 6H), 6.88 (s, 1H), 6.86 (d, *J* = 8.3 Hz, 1H). ^13^C NMR (150 MHz, DMSO-*d*_*6*_) δ 169.6, 165.3, 165.2, 139.2, 136.8, 135.0, 134.9, 132.5, 131.5, 131.4, 128.42, 128.36, 127.59, 127.55, 125.3, 123.9, 121.7, 109.8, 109.1. HRMS m/z [M+H]^+^ calcd for C_19_H_15_N_4_O_2_: 331.1195, found 331.1185, LC t_R_ = 3.56 min, > 98% Purity.

## Notes

The authors declare no competing financial interests.

## Acknowledgement

The SGC is a registered charity (number 1097737) that receives funds from AbbVie, Bayer Pharma AG, Boehringer Ingelheim, Canada Foundation for Innovation, Eshelman Institute for Innovation, Genome Canada, Innovative Medicines Initiative (EU/EFPIA) [ULTRA-DD grant no. 115766], Janssen, Merck KGaA Darmstadt Germany, MSD, Novartis Pharma AG, Ontario Ministry of Economic Development and Innovation, Pfizer, São Paulo Research Foundation-FAPESP, Takeda, and Wellcome [106169/ZZ14/Z]. The US National Institutes of Health (NIH) is acknowledged for support (1U24DK11604-01). We thank Biocenter Finland/DDCB for financial support and the CSC-IT Center for Science Ltd. (Finland) for allocation of computational resources. We also thank Dr. Brandie Ehrmann and Ms. Diane E. Wallace for LC-MS/HRMS support provided by in the Mass Spectrometry Core Laboratory at the University of North Carolina at Chapel Hill. The core is supported by the National Science Foundation under Grant No. (CHE-1726291). In addition, we thank the University of North Carolina’s Department of Chemistry NMR Core Laboratory for the use of their NMR spectrometers, along with Dr. Marc ter Horst and Dr. Andrew Camp for their assistance with NMR spectroscopy. The core is supported by the National Science Foundation under Grant No. (CHE-1828183).

## Appendix A. Supplementary data

Supplementary data to this article can be found online at https://doi.org/10.1016/j.ejmech.xxxxxx

